# A relative-motion method for parsing spatio-temporal behaviour of dyads using GPS relocation data

**DOI:** 10.1101/2021.01.04.425238

**Authors:** Ludovica Luisa Vissat, Jason K. Blackburn, Wayne M. Getz

**Affiliations:** Dept. ESPM, University of California, Berkeley, 130 Hilgard Way, Berkeley, CA 94720-3114, USA; Spatial Epidemiology and Ecology Res. Lab., Department of Geography, University of Florida, 330 Newell Dr, Gainesville, FL 32611, USA; Emerging Pathogens Institute, University of Florida, 2055 Mowry Road, Gainesville, FL 32610, USA; School of Mathematical Sciences, University of KwaZulu-Natal, 238 Mazisi Kunene Rd, Glenwood, Durban, 4041, South Africa

**Keywords:** African elephant, approach and retreat, biased random walk, dyadic interactions, *Loxodonta africana*

## Abstract

1. In this paper, we introduce a novel method for classifying and computing the frequencies of movement modes of intra and interspecific dyads, focusing in particular on distance-mediated approach, retreat, following and side by side movement modes.
2. Besides distance, the method includes factors such as sex, age, time of day, or season that cause frequencies of movement modes to deviate from random.
3. We demonstrate and validate our method using both simulated and empirical data. Our simulated data were obtained from a relative-motion, biased random-walk (RM-BRW) model with attraction and repulsion circumferences. Our empirical data were GPS relocation time series collected from African elephants in Etosha National Park, Namibia. The simulated data were primarily used to validate our method while the empirical data analysis were used to illustrate the types of behavioral assessment that our methodology reveals.
4. Our methodology facilitates automated, observer-bias-free analysis of the locomotive interactions of dyads using GPS relocation data, which is becoming increasingly ubiquitous as telemetry and related technologies improve. Our method should open up a whole new vista of behavioral-interaction type analyses to movement and behavioral ecologists.

## 1. Introduction

Most studies of dyadic interactions involve making observations of individuals at close quarters (Whitehead and Dufault, 1999); for example, agonistic interactions in social insects (Getz and Smith, 1986, Breed, 2003), grooming networks in primates (Voelkl et al., 2011), and dominance behavior in elephants (Archie et al., 2006, Wittemyer and Getz, 2007). An exception though are a new class of methods that use global positioning system (GPS) telemetry data to assess the joint movement of individuals that may be some distance apart and not simultaneously directly observable to a visual recorder (human or camera) (Joo et al., 2018). Of course, the assumption is that individuals not in visual contact with one another may still have auditory (Hulse, 2002, Erbe et al., 2016), olfactory (Shorey, 2013), or even low frequency vibratory cues (McComb et al., 2003, O’Connell-Rodwell, 2007) regarding the location of other individuals within a radius and direction salient to the perceptual modality involved (with wind direction playing a critical role in olfactory communication).

In this paper, we continue to develop the opportunities for evaluating dyadic movement interactions using GPS relocation data by presenting a technique that allows us to classify dyadic modes of movement in terms of distance-dependent approach, retreat, following, and side by side modes of movement. Other salient factors such as the sexes and ages of the individuals, the time of day or year, the internal state of the individuals and the state of the environment may also be included in an extended version of our method. Specifically, our method is based on the analysis of pairs of animal trajectories with overlapping time periods, comparable frequencies and without major gaps in the data collection. By considering individual directions and locations, our approach allows us to extract both individual and dyadic behaviors, whether symmetric (e.g., both individual moving towards each other) or asymmetric (e.g., one advancing and one retreating). This new methodology extends existing analysis: it allows us to understand details of dyad interactions by classifying different behaviour types and grouping the results based on distance apart, which can be up to several kilometers for individuals using auditory communication (McComb et al., 2003, O’Connell-Rodwell, 2007).

In the case of two individuals moving in the same general direction, whether they are moving side by side or one is pursuing the other will come down to an assessment based on relative speeds and headings, and, ultimately, the terminal behavior at the end of the event or even the identity of the individuals involved (e.g., two predators pursuing the same unknown prey versus a predator pursuing a known prey).

Since our method is novel, we need to demonstrate both its validity and its utility. We undertake the former by applying our method to simulated data with known dyadic interactions to see how well the method uncovers these interactions embedded in movement data. We undertake the latter by analysing GPS data obtained from GPS collared African elephants (*Loxodonta africana*), in Etosha National Park, Namibia (Abrahms et al., 2017, Tsalyuk et al., 2019). Our simulation data are generated by a novel relative-motion, biased random-walk (RM-BRW) model with attraction and repulsion circumferences constructed and implemented using the Numerus Model Builder (NMB) simulation platform (Getz et al., 2018). Although we use our simulation data to demonstrate the ability of our method to correctly identify known behaviors embedded in the model, it can also be used to test various hypotheses about the structure of empirical data, particularly in the context of evaluating whether or not particular movement patterns differ significantly from patterns that may be generated at random.

In the rest of this paper, in Section 2 we first introduce our general method and we next report details of the method’s implementation. This is followed by our description of the model used to generate simulated data and a description of our empirical data. We then present a report in Section 3 of the application of our method to the simulated and to the empirical data. Finally, we present a discussion of related work in Section 4 and then end with concluding remarks in Section 5.

## 2. Materials and Methods

### 2.1. Method

We developed our method to study, in particular, how approach and retreat behavior in pairs of individuals may depend on the current distance between them, using locations recorded at the same, or close to the same, times. Since direction of movement and speed are needed to classify the different behaviour types, consecutive relocation points are needed of the type obtained using GPS telemetry data (Calenge et al., 2009). In addition, the frequency of location sampling should be sufficiently high to ensure that estimates of direction and speed are relevant to the scale of the analysis (Codling and Hill, 2005). GPS collar battery life creates a trade-off between the frequency of GPS point collection and the total tracking period for an animal. The empirical study undertaken here includes data in the 15-30 minute sampling range, which is frequently reported in terrestrial animal movement studies. Hence this range will be the focus of our discussion.

To begin, the relocation data from pairs of individuals are time-matched so that consecutive relocation points obtained from these individuals can be used to compute heading directions and speed for each. Although these computed directions and speeds for each individual may not be perfectly matched in time, they are nominally labeled as occurring at common times *t*, *t* + 1, *t* + 2, and so on if deviations from these times are sufficiently small compared to the size of inter-sampling interval: e.g., if points in both relocation sets are collected every 15 minutes, then we may decide that points collected in different sets within a threshold of 2 or 3 minutes of one another can be matched up, where this threshold may be varied to see how results are affected. Our method then proceeds by a considering time series vector set 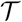 containing the following information at each time point *t* regarding two individuals labeled A and B and their positions at time *t* and *t* + 1 : viz., the absolute headings (headA, headB), the relative headings (dirAB, dirBA) (which we define as direction towards the other individual in the dyad), the difference between absolute and relative headings (diffA, diffB), the speed (sA, sB), and the pair distance (*d*_AB_). Either some or all of these values will be used in the different analysis (individual, dyadic, extended) proposed in this paper. Thus our time series is:

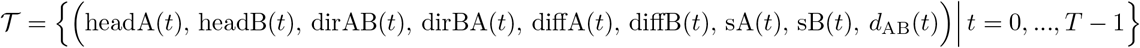

In the following, we describe how we compute the entries for individual A with respect to individual B. For the *t*^*th*^ entry in 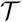, this computation requires the positions of A and B at time *t* and A at time *t* + 1, which we denote by *A*(*t*), *B*(*t*) and *A*(*t* + 1) (Fig. 1, left panel).

**Figure 1:**
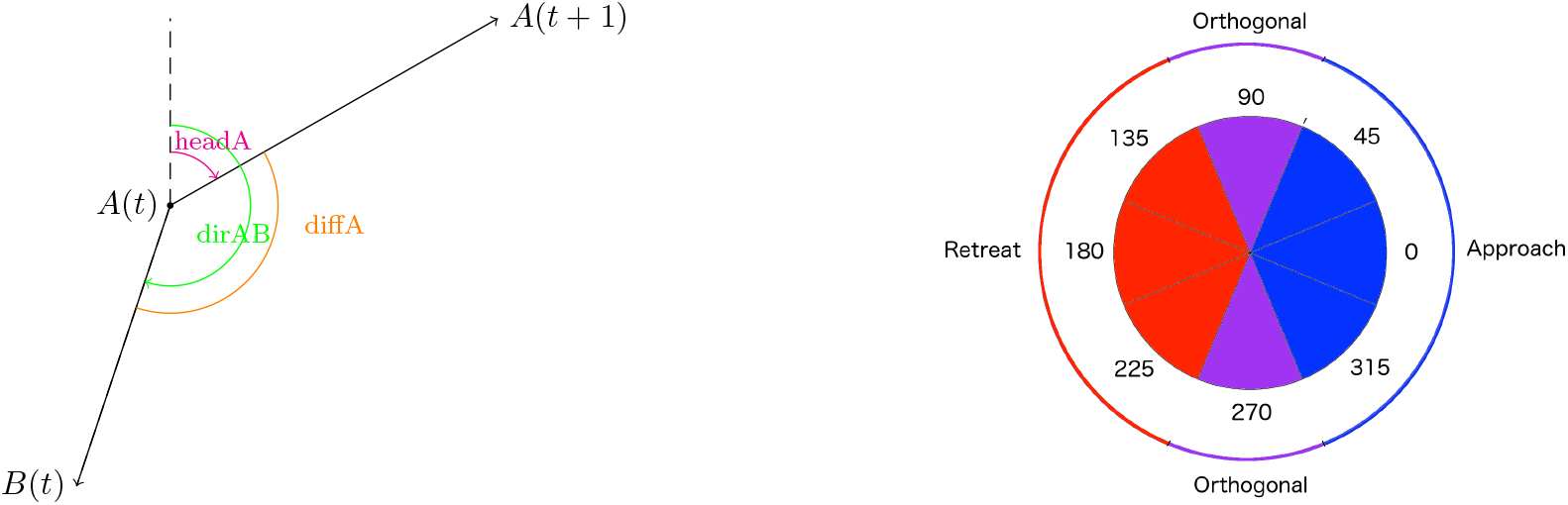
Left panel: absolute heading (headA), relative heading (dirAB) and difference between these two angles (diffA) for individuals A in relation to individual B. Right panel: classification of absolute value of the difference among the individual absolute and relative heading, based on an 8 equal tranche segmentation of the unit circle (numbers represent the central angle value of each tranche, where 0 ≡ 360). Other levels of segmentation, say twelve, can be used and results obtained can be compared for sensitivity of the results obtained to a more demanding scale of segmentation (in the 12 segment case, the purple areas encompasses two 30 rather than 45 degree tranches). Blue represents approach behaviour, red represents retreat behaviour while purple represents orthogonal behaviour.

In calculating angles (e.g., headA, dirAB and diffA for individual A), we use the convention that north is 0 and measure clockwise, following the convention used by the function `bearing` in the R package `geosphere`. The speed of individual A at time *t* is calculated as the ratio of the distance *d(A*(*t*)*, A*(*t* + 1)) and the time interval between *t* and *t* + 1. The entry *d*_AB_ is the pair distance *d(A*(*t*)*, B*(*t*)). The values of the first two calculated angles (e.g., headA and dirAB for individual A) lie in [0, 360) and therefore the absolute value of their difference (e.g., diffA for individual A) lies in the same interval. We divide the turning circle into 8 sections (Fig. 1, right panel), and consider individual A to be *approaching* individual B if diffA is below 67.5 or above 292.5 (area shown in blue), while if diffA is between 112.5 and 247.5 (area shown in red) A is considered to be *retreating* from B. If diffA lies between 67.5 and 112.5 or between 247.5 and 292.5, we refer to the movement as *orthogonal*. Of course, this partitioning of the turning circle can be varied and the sensitivity of results to this partition evaluated.

### 2.2. Individual behavior

We first focus on the movement of an individual A, though in the context of individual B, as represented by that time values headA(*t*), dirAB(*t*), and diffA(*t*). We begin by identifying the following three behavioral modes (Fig. 1, with examples shown in Fig. 2):

**Figure 2:**
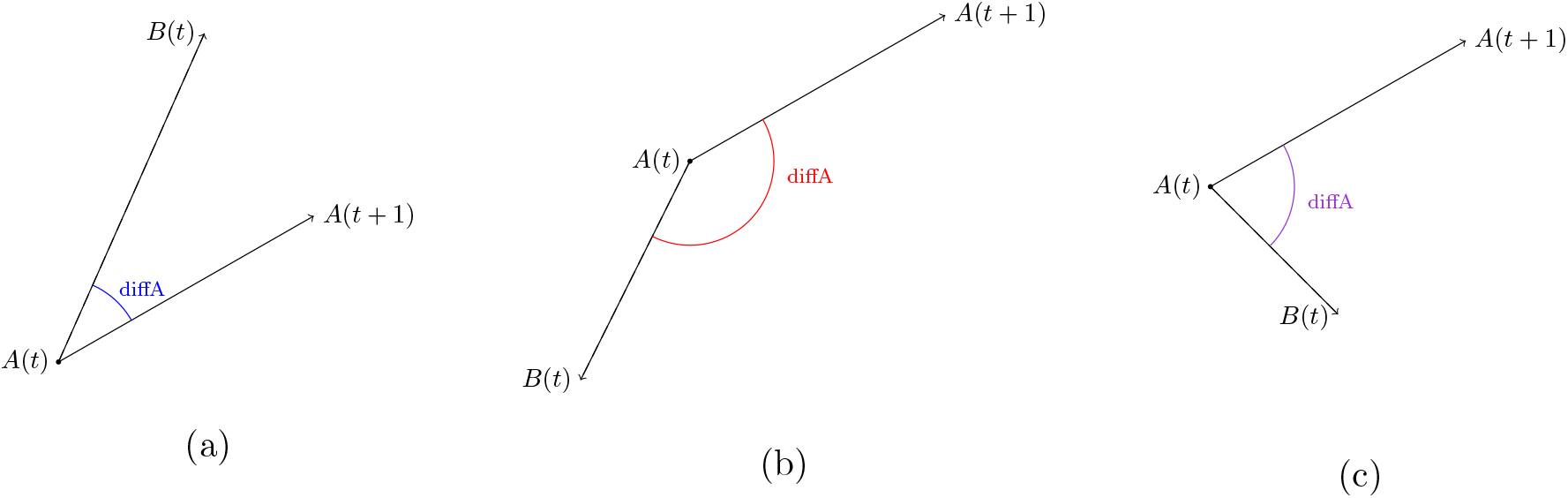
Example of behaviour classification for individual A: (a) approach, (b) retreat, (c) orthogonal individual behaviour, in relation to individual B.

a. A approaches B: blue segments in Fig. 1
b. A retreats from B: red segments in Fig. 1
c. A moves orthogonally to B (i.e., around 90 or 270 degrees): purple segments in Fig. 1

In particular, we are interested in how these approach, retreat and orthogonal modes may depend on the distance between A and B, as well as influenced by temporal (diurnal, seasonal) and local environmental (landscape features, other individuals present, weather conditions, etc.) factors. Thus, in the first instance, we bin the classification of individual behaviors in relation to the distance between pairs of individuals. Specifically, for each individual in the dyad and for each chosen distance interval, we count the number of points at which approach, retreat and orthogonal movements are detected. We then evaluate whether or not the proportion of approaches to retreats are significantly different from random: given the symmetry of the problem, once the orthogonal movement points are removed, this proportion should be not be significantly different from 0.5. If it is, we can conclude that individual A’s behavior with respect to be B is one of approach or retreat, as the case may be.

However, it is important to note that in the case where the number of retreats and approaches do not differ significantly from each other, they may still differ from random once the orthogonal points have been taken into account. Essentially, in this case we can conclude that approach and retreat behaviors, though they may be equally likely, are directed and hence intended when they occur. Thus the analysis when orthogonal movement designations are considered can be used to see to what extent approach and retreat behaviors are directed. Note that the larger the purple area in Fig. 1 the more the movement points are directed when the approach or retreat points are significantly different from random.

### 2.3. Dyadic behaviour

Our method also includes the identification of dyadic behavioral modes beyond the classification of how individuals move with respect to one another. In particular, at matching points for each dyad, we assign one of the following six dyadic modes of behavior:

1. both individuals approach each other
2. both individuals retreat from each other
3. one individual approaches while the other individual retreats
4. one individual moves orthogonally, the other approaches
5. one individual moves orthogonally, the other retreats
6. both individuals move orthogonally

**Figure 3:**
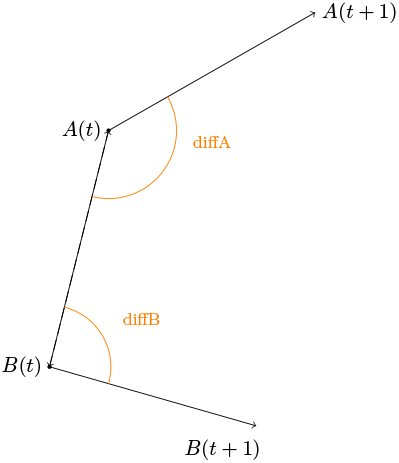
diffA and diffB: angle difference between the absolute and the relative headings for individuals A and B respectively.

As with the individual behavioral modes, we can use sample sizes and the expected proportions of each of the behavioral modes to assess whether or not a particular dyadic mode occurs significantly more often than expected at random over a particular set of intervals of time or in particular locations on the landscape. Specifically, if we exclude dyadic time-matched points at which one of the individuals is moving orthogonally, we expect the proportion of modes 1-3 to be 0.25, 0.25 and 0.5—the latter when the roles of A and B are interchangeable. Otherwise, we expect the proportion to all be equal to 0.25 if we consider mode 3 in the context of “A approaches while B retreats” compared with “A retreats while B approaches.”

### 2.4. Extended analysis involving absolute headings and relative speed

We build on the previous analysis to extract dyadic behaviour such as *A following B* or *A and B walking side by side*. To achieve this, we consider also the *relative speed* of the individuals in the pair and the difference between their *absolute headings*. The speed of an individual at a particular point is calculated in terms of its distance to the next consecutive point, for all its relocation points before these points are time matched with the other’s points, as described above. Time-matched points then inherit the speed calculation associated with each of these matched points. We then categorize the *relative speed* of a dyad (A,B) as *similar speed*, *A faster than B* or *B faster than A*. To each of the six dyadic modes described in the previous section, we can now assign an additional designator: a.) similar speed, b.) A faster than B, 3.) B faster than A. This yields a total of 18 dyadic movement modes: 1a, 1b, …, 6c. This additional designator, combined with the following heading analysis, will give us the opportunity to extract specific behaviours of interest, as elaborated in Section 3.3. In the SOF, we provide a table with all the dyadic movement modes and the description of the behaviours of interest.

For an *a priori* threshold value *θ* (which may vary in a sensitivity analysis of results to this parameter value) in the context of individual headings, we classify a dyad as having *a similar heading* if the absolute value of the difference is below *θ* or above 360 *− θ* (shown in cyan in Fig. 4), while we classify it as having *the opposite heading* if the difference lies between 180 *− θ* and 180 + *θ* degrees (shown in orange in Fig. 4).

**Figure 4:**
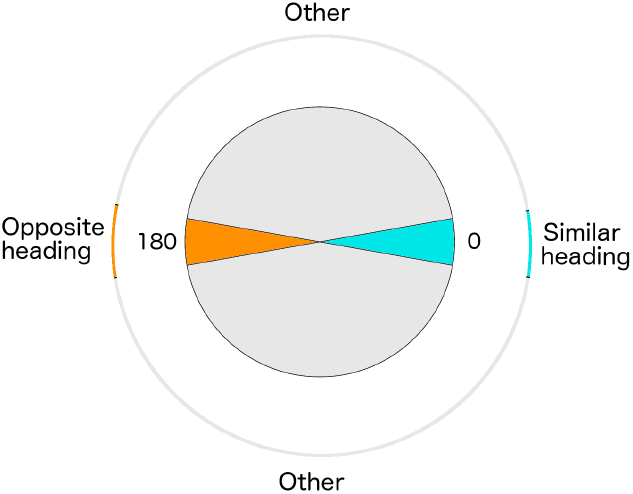
Heading difference for a dyad where, in this case, the angle *θ* referred to in the text is 10 degrees. Cyan values are classified as a similar heading dyad, orange values as an opposite heading dyad, with all other cases shown in gray.

Given this additional classification, we can now combine all the different information—the dyadic behaviour, the relative speed and the heading difference—to extract behaviour of interest and compare its frequency of occurrence to that of other behaviours or, if desired, values expected at random. We can also compare the frequencies of the behaviours of interest, looking at difference among seasons, sex-type of individuals in the pair, or time of the day. For example, we may be interested in the following behaviour types:

- *Following behaviour*: dyadic behaviour of type 3a (dyadic behaviour of type 3, similar speed) and similar heading.
- *Side by side movement*: dyadic behaviour of type 6a (dyadic behaviour of type 6, similar speed) and similar heading.

These behaviours refine the previous dyadic behaviour of type 3 and 6, where the individuals also have similar speed and similar absolute heading.

### 2.5. Implementation

We perform this novel analysis using the R programming language, through the integrated development environment RStudio, and different R packages. We used the function `bearing` and the function `distm` from the R package `geosphere` to calculate the angles and the dyadic distance respectively; we used the function `binom.confint` from the R package `binom` to obtain the confidence intervals. Since the function `bearing` allows us to calculate the angle between the North direction and a given vector, expressed with longitude and latitude coordinates, the calculation of the angle diffA, between the two vectors shown in Fig. 1, was defined as the difference between the absolute (headA) and relative (dirAB) heading, as described in Section 2.1. Algorithm 1 describes the analysis of individual behaviour for both individual A and B, while the procedures for dyadic and extended analysis are provided in the SOF. For the sake of simplicity, we assume that the trajectories have location points that where recorded at almost the same time or have been interpolated to match up in time.

**Algorithm 1:**
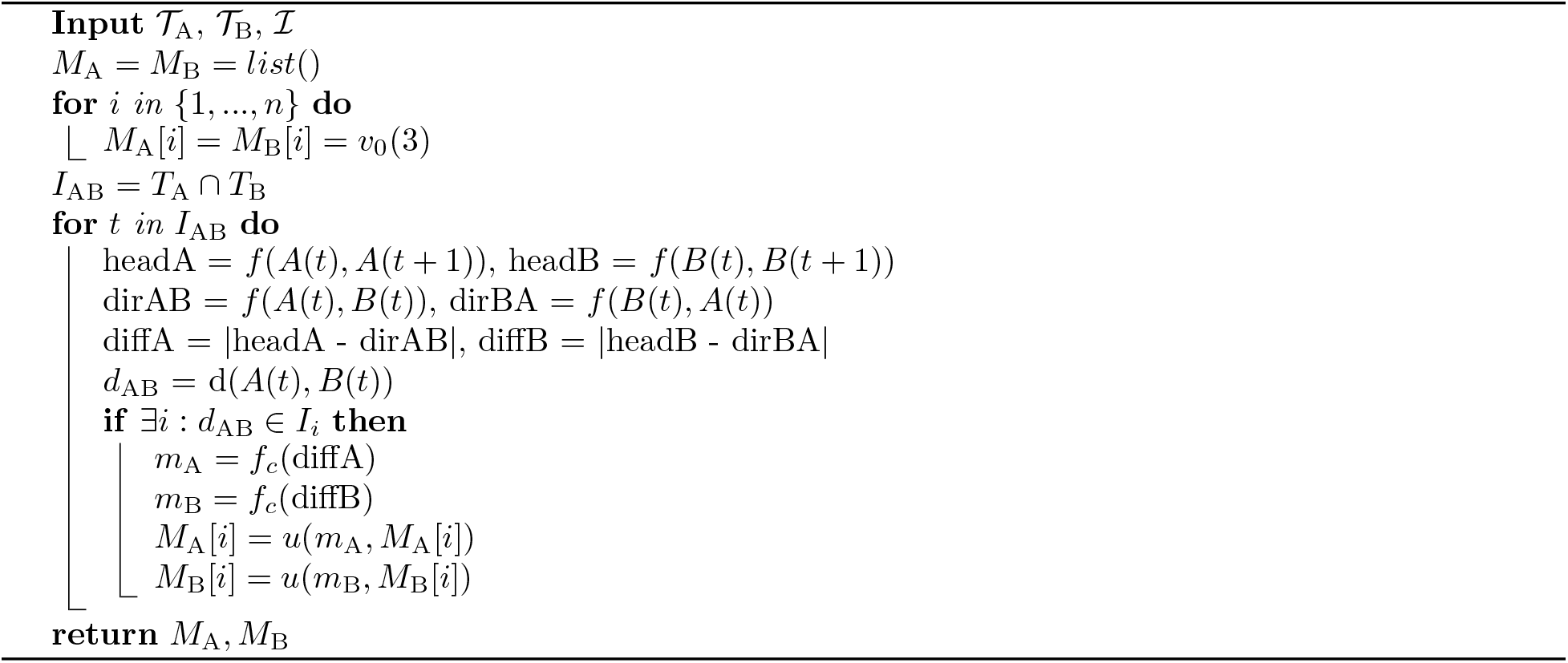
Individual behaviour analysis

The required inputs for the individual behaviour analysis are the trajectories for individual A and B (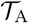 and 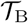), and the *n* distance intervals 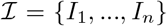 used to bin the results. We use the lists *M*_A_ and *M*_B_ to record the counts related to the different individual behaviour types and to the different *n* distance intervals. *M*_A_ and *M*_B_ are initiated as lists of *n* vectors, indicated with *v*_0_(3), which are vectors of length 3 and entry all equal to 0. Each of these vectors is used to keep track of the counts for each of the 3 behaviour types for the *n* different distance intervals.

The algorithm then calculates the set *I*_AB_, the intersection between the time point sets *T*_A_ and *T*_B_, thereby identifying the sets of time points in both trajectories 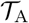 and in 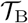. For each of points in *I*_AB_, the procedure evaluates the three different angles for each individual: the function *f* calculates the angle between the North direction and either the absolute heading vector (angles: headA and headB) or the relative heading vector (angles: dirAB and dirBA). The algorithm then calculates the angles *difference* diffA and diffB, as well as the distances *d*_AB_. If *d*_AB_ lies within any of the chosen distance intervals *I*_*i*_, then the algorithm proceeds by classifying the angle *difference* as shown in Fig. 1 through the function *f*_*c*_. The function *u* then updates the lists *M*_A_ and *M*_B_, as a function of the angle classification and the corresponding distance interval *I*_*i*_. The outputs of this procedure are the lists *M*_A_ and *M*_B_, which are used to construct barplots and perform the statistical analysis. In particular, the statistical analysis uses the entries corresponding to the approach and retreat behaviours for the calculation of the confidence intervals related to each *I*_*i*_. The R code for this analysis, the dyadic behaviour and the extended analysis can be found at https://ludovicalv.github.io/Dyadic_behaviour_method/.

### 2.6. Simulated data

To validate our individual and dyadic methods of analysis, we applied them to simulated data with known properties to see how well our methods could capture these properties. Our simulation models were constructed using the Numerus Model Builder (NMB) platform (Getz et al., 2018). These NMB models and simulations used in the analysis can be found at https://ludovicalv.github.io/Dyadic_behaviour_method/. Beyond providing test data for this study, our NMB relative-motion, biased random-walk (RM-BRW), as described below, can also be used to explore theoretical question or help design empirical studies. For example, our simulator can be used to evaluate how easily behaviors of different durations can be detected with data collected at particular frequencies. This information would then inform the choice of data collection frequency, battery use, or collar deployment. Additionally, our simulator can be used to test how sensitive different algorithms may be to detecting movement path structures that have been defined at various spatio-temporal scales (Getz et al., 2020).

To generate the set of simulated data used to validate our novel method, we model a relative-motion, biased random-walk (RM-BRW) model with approach and retreat circumferences. In particular, for distances less then *d*_*R*_, A and B repulse one another, between *d*_*R*_ and *d*_*I*_ they attract one another and above *d*_*I*_ they behave independently. Additionally, the approach and retreat behaviours only occur at each time step with a given probability *p*_eff_ and with noise introduced by a coefficient *ρ ∈* [0, 1]. A complete mathematical description of the model is provided in our SOF. The NMB modelling panel is shown in Fig. 5.

**Figure 5:**
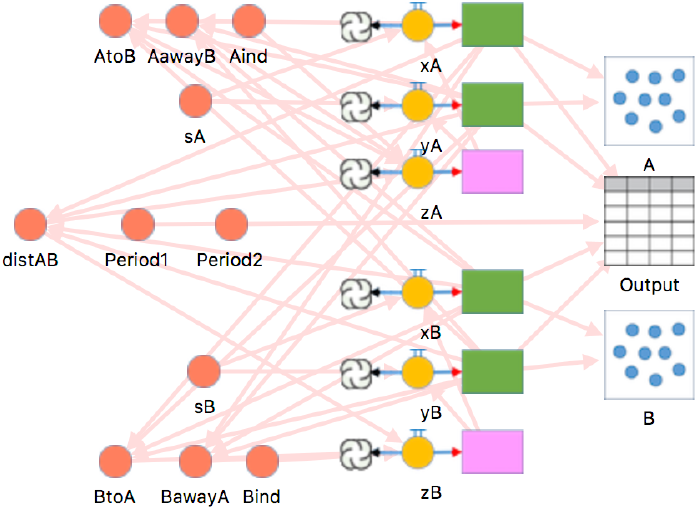
NMB panel: RM-BRW model

We also extended the model to include a time-dependent component to the attraction and repulsion behaviour. Specifically, we allowed attraction to operate only during the first quarter of the day, repulsion during the second quarter of the day, and independent movement for the remaining half day.

### 2.7. Empirical data

We applied our method to GPS relocation data from 39 African elephants, collected between 2008 and 2015 in Etosha National Park, Namibia (Abrahms et al., 2017, Tsalyuk et al., 2019). The intervals at which these data were recorded varied among 4, 3 and 2 points per hour (10, 15, and 14 individuals respectively). As mentioned in Section 2.1, these data collection frequencies are commonly reported in the GPS movement literature. Data were collected over different periods ranging from 2.5 months to 4.5 years. The time line of data collection for each individual and for each different frequency is provided in our SOF.

## 3. Results

### 3.1. Simulations

We simulated paths for A and B over a 10 day period, using a time step of 1 minute. Our unit spatial measure was set to 10 m and the repulsion and attracting circle diameters were set to *d*_*R*_ = 30 and *d*_*I*_ = 60 (i.e., 300 and 600 m respectively). The results reported in the first two subsections are for the time-independent version of the model, while those reported in the third subsection are for the time-dependent version of the model.

#### 3.1.1. Individual behaviour

To show the distribution of the three different individual behaviour types, we use barplots and the 3-colours legend introduced in Section 2.1 (Fig. 1). Specifically, we identified and binned the frequencies of approach, retreat and orthogonal behaviours using equal-width classes of pair distance, starting with a 0-5 unit bin and ending with a 75-80 unit bin, as shown in Fig. 6. Note that here we classify these approach/retreat/orthogonal behaviours per individual, with the results for the dyad as a whole reported in the next subsection. In Fig. 6 (left panel) we show the results for individual A only, since individuals A and B are symmetric in the model (see SOF for further details). In addition, in Fig. 6 (central panel), we display the ratio between the approach count and the total of the approach and retreat counts, and the calculated 95% confidence intervals for each distance interval. We colored the results that are significantly larger than 0.5 in blue (approach) or less then 0.5 in red (retreat). As expected, we observe more retreats (red) for smaller distances, while the number of approaches (blue) increases with pair distance. Since we set *d*_*I*_ = 60 the pair distance tends to remain below *d*_*I*_ most of the time.

**Figure 6:**
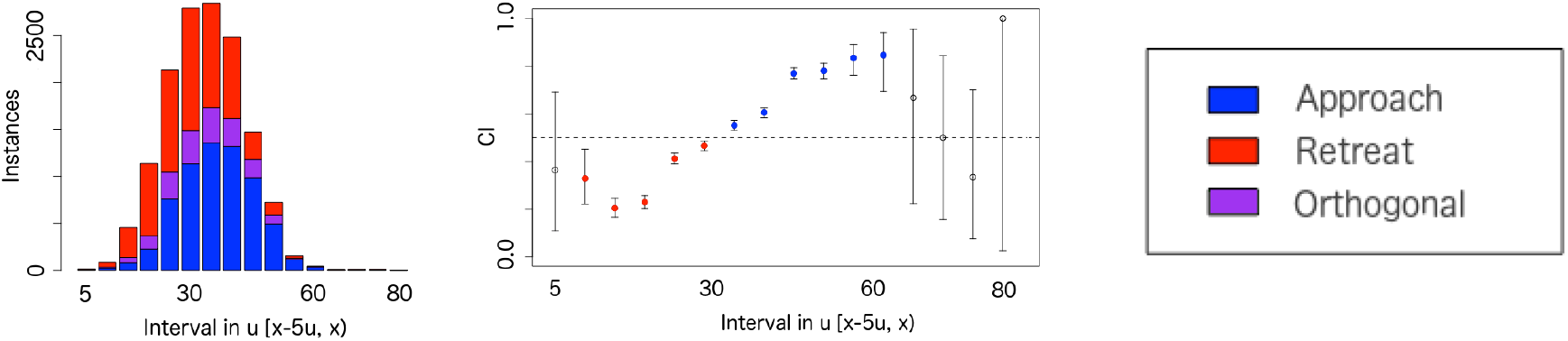
Analysis barplot for individual A (left). Estimated confidence intervals (CI) for individual A, colored if showing a statistically significant result (center), according to the legend (right).

As introduced before, in this particular example *d*_*R*_ was chosen to be equal to 30 and *d*_*I*_ equal to 60 units. We evaluate the statistical analysis results for the distance intervals: [0,30), [30,60) and [60,80). We provide the results of this analysis in Table 1, where we indicate statistically significant results above 0.5 (approach) or below 0.5 (retreat) according to the results. As expected, we observe that for distance below 30 units, the results are statistically significant and show retreat behaviour, while for distance in [30,60) the results show approach behaviour. We do not observe a high number of instances and any statistically significant results for distance above 60 units.

**Table 1:** Results of the individual analysis. We report the results grouped by distance intervals, providing the number of approaches and considering the total number of approach and retreat behaviours. We then show the bound of the confidence interval (CI) and check the result that are statistically significant at the 95% level.

| Individual | Distance interval (units) | Total | Approach | Lower CI | Upper CI | Approach | Retreat |
| --- | --- | --- | --- | --- | --- | --- | --- |
| A | $[0,30)$ | 5779 | 2236 | 0.3743 | 0.3996 | | ✓ |
| B |  | 5821 | 2196 | 0.3648 | 0.3899 |  | ✓ |
| A | $[30,60)$ | 6744 | 4311 | 0.6276 | 0.6507 | ✓ | |
| B |  | 6798 | 4367 | 0.6309 | 0.6538 | ✓ |  |
| A | $[60,80)$ | 24 | 12 | 0.2912 | 0.7088 | | |
| B |  | 16 | 10 | 0.3543 | 0.848 |  |  |

#### 3.1.2. Dyadic behaviour

By considering the behaviour of both individuals A and B, we compute the number of incidences of dyadic type 3 mode of behavior separately for A and B: viz.,

- 3(A,B): individual A approaches while individual B retreats
- 3(B,A): individual A retreats while individual B approaches

We also compute the number of instances of dyadic type 1 and 2 behaviors and exclude cases where orthogonal movements are involved (types 4-6). Our expectation, under a purely random movement hypothesis, is that each of these four instances (1, 2, 3(A,B), 3(B,A)) should occur with frequency 0.25. The number of instances of the various modes is illustrated in the barplot shown in Fig. 7, while the statistical analysis of the significance of the deviations from 0.25 at the various distances can be found in the table in SOF. As expected retreats occur significantly more often than random below 300 m (30 units) while approaches occur significantly more often than random between 300 and 600 m (30-60 units).

**Figure 7:**
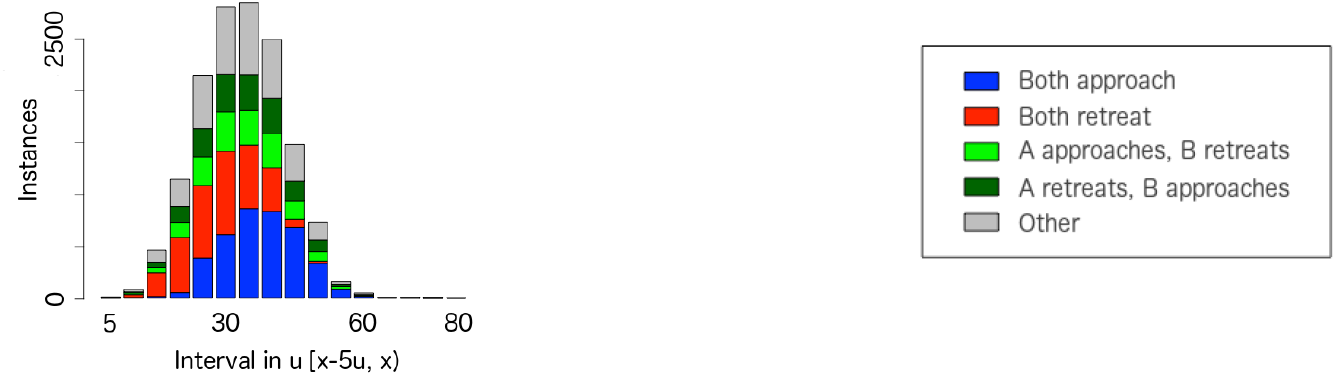
Analysis barplot for pair behaviour with legend provided on the right. We note that we observe significantly more than expected retreat behaviours (red) for distance below 30 (300 m), and significantly more approach behaviors than expected (blue) for distances above 30.

The results present in this and the previous subsection individuates that our method extracts the expected deviations from random that are a result of the biases that we have built into our model in terms of retreating behavior at *<*30 units and attracting behaviour at *>*30 units Our analysis also allows to observe dyadic behaviour over time. For instance, in Fig. 8, we show the pair distance, coloring each point in time with the color corresponding to the dyadic behaviour type. We observe mostly red points at shorter distances, where the retreat behaviours predominate, and mostly blue points at larger distances, where the approach behaviours predominate.

**Figure 8:**
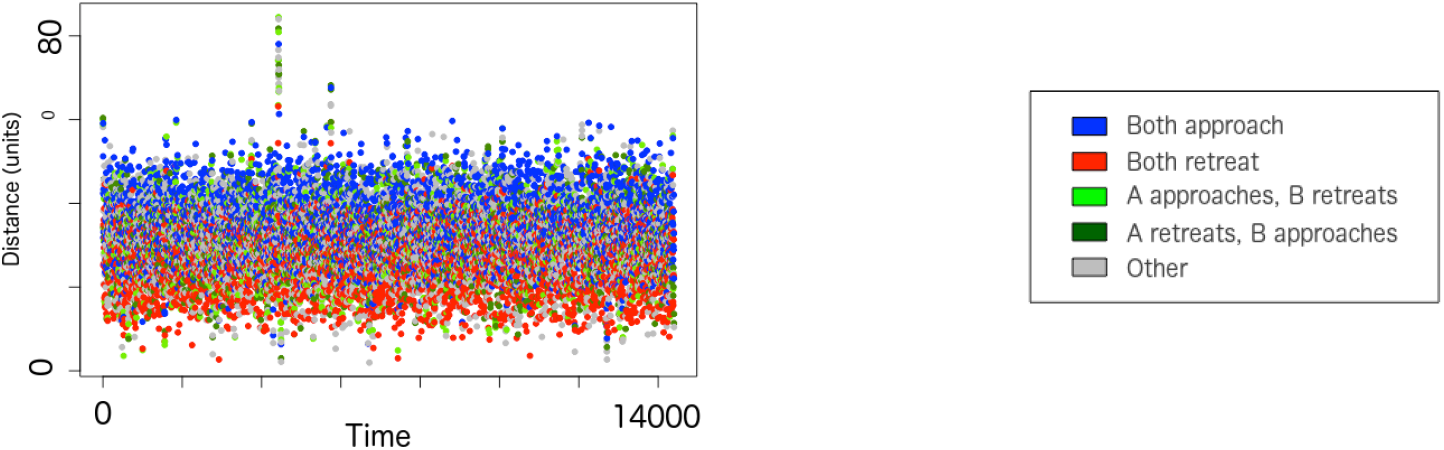
Distance between the pair (left), colored according to the dyadic behaviour type as shown in the legend (right). Also in this case, we observe mostly retreat behaviours (red) for shorter distance while mostly approach behaviours (blue) for bigger distances.

#### 3.1.3. Time-dependent case

In the case of the time-dependent model, we observe (Fig. 9) as expected that during the repulsion period both individuals move away from each other (indicated in red), while during the attraction period they both move towards each other (indicated in blue). When the individuals move independently we observe a less substantial change of the pair distance. A distance-dependent behaviour analysis across all time is not be able to capture the temporal patterns. By subdividing the data according to time of the day, however, we are able to observe time-dependent behaviour patterns, as documented in our SOF.

**Figure 9:**
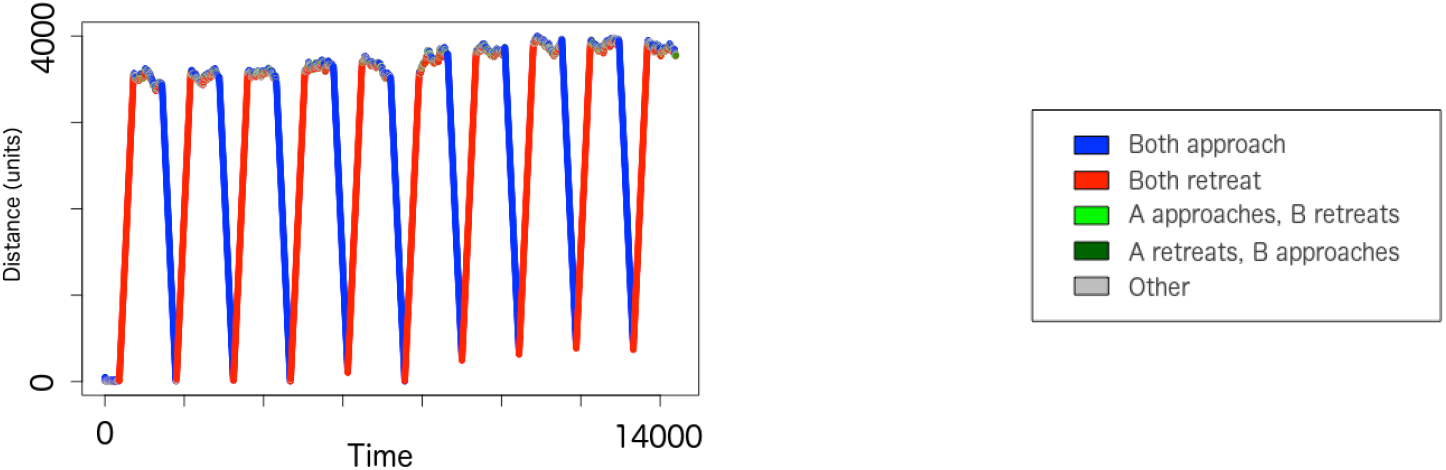
Distance between the pair (left), colored according to the dyadic behaviour type as shown in the legend (right). We observe that the distance between individuals increases with retreat behaviour (red) and decreases with approach (blue).

### 3.2. Empirical results

We first identified among the full set of elephant data those dyads that had matching frequencies within overlapping time periods and were close enough to one another for our method to yield interpretable results. In particular, we selected dyads that presented at least 500 instances at a distance below 1 km and had matching frequencies. In the first subsection below, we present the analysis results for a specific female-male dyad of interest. The male of this chosen pair had been tagged (Tsalyuk et al., 2019) as “with breeding herd” of the female. Given the repeated interactions, it is likely that the male was the son of one of the females in the same herd. In subsection 3.2.3, we provide a general overview of the results for all pairs that fit our matching criteria.

#### 3.2.1. Individual behaviour

As presented in Section 3.1.1, we use barplots to show the distribution of the three different individual behaviour types, grouped by pair distance. In this case, we identified and binned the frequencies of approach and retreat behaviours using 9 different size classes, starting with a 0-50 m bin and ending with a 5-10 km bin, as shown in Figs. 10a and b. In addition, in Fig. 10c and d, we show the value of the ratio between the approach count and the total of approach and retreat counts, and the calculated confidence intervals. These results were colored if statistically significant: in blue for approach and in red for retreat individual behaviour. We provide the values of the analysis in Table 2, where we indicate statistical significant results (approach and retreat) with a check mark. We observe that the female retreats (Figs. 10a and c) significantly more often then random at distances between 50m-3 km while the male approaches significantly more often than random at distances *<* 3 km.

**Table 2:** Results of the individual analysis for female and male. We report the results grouped by distance intervals, providing the total number of approach and retreat behaviours and the number of approach ones. We then show the bound of the confidence interval (CI) and if the result was statistically significant showing approach or retreat behaviours. Approach behaviours are mostly shown by the female while the male shows mostly retreat behaviours.

| Ind. in the pair | Distance interval (m) | Total | Approach | Lower CI | Upper CI | Approach | Retreat |
| --- | --- | --- | --- | --- | --- | --- | --- |
| Female | [0,50) | 584 | 289 | 0.4536 | 0.5362 |  |  |
| Male |  | 567 | 313 | 0.51 | 0.5935 | ✓ |  |
| Female | [50,100) | 613 | 275 | 0.4088 | 0.489 |  | ✓ |
| Male |  | 603 | 326 | 0.4999 | 0.581 |  |  |
| Female | [100,200) | 1040 | 498 | 0.4481 | 0.5097 |  |  |
| Male |  | 1060 | 567 | 0.5043 | 0.5653 | ✓ |  |
| Female | [200,500) | 2115 | 967 | 0.4358 | 0.4787 |  | ✓ |
| Male |  | 2125 | 1197 | 0.5419 | 0.5845 | ✓ |  |
| Female | [500,1000) | 1691 | 653 | 0.3629 | 0.4098 |  | ✓ |
| Male |  | 1763 | 1115 | 0.6094 | 0.655 | ✓ |  |
| Female | [1000,2000) | 1308 | 494 | 0.3513 | 0.4046 |  | ✓ |
| Male |  | 1361 | 877 | 0.6183 | 0.6698 | ✓ |  |
| Female | [2000,3000) | 924 | 362 | 0.3601 | 0.4241 |  | ✓ |
| Male |  | 938 | 529 | 0.5315 | 0.596 | ✓ |  |
| Female | [3000,5000) | 1266 | 635 | 0.4737 | 0.5295 |  |  |
| Male |  | 1319 | 646 | 0.4625 | 0.5171 |  |  |
| Female | [5000,10000) | 2764 | 1375 | 0.4787 | 0.5163 |  |  |
| Male |  | 2809 | 1458 | 0.5004 | 0.5377 | ✓ |  |

**Figure 10:**
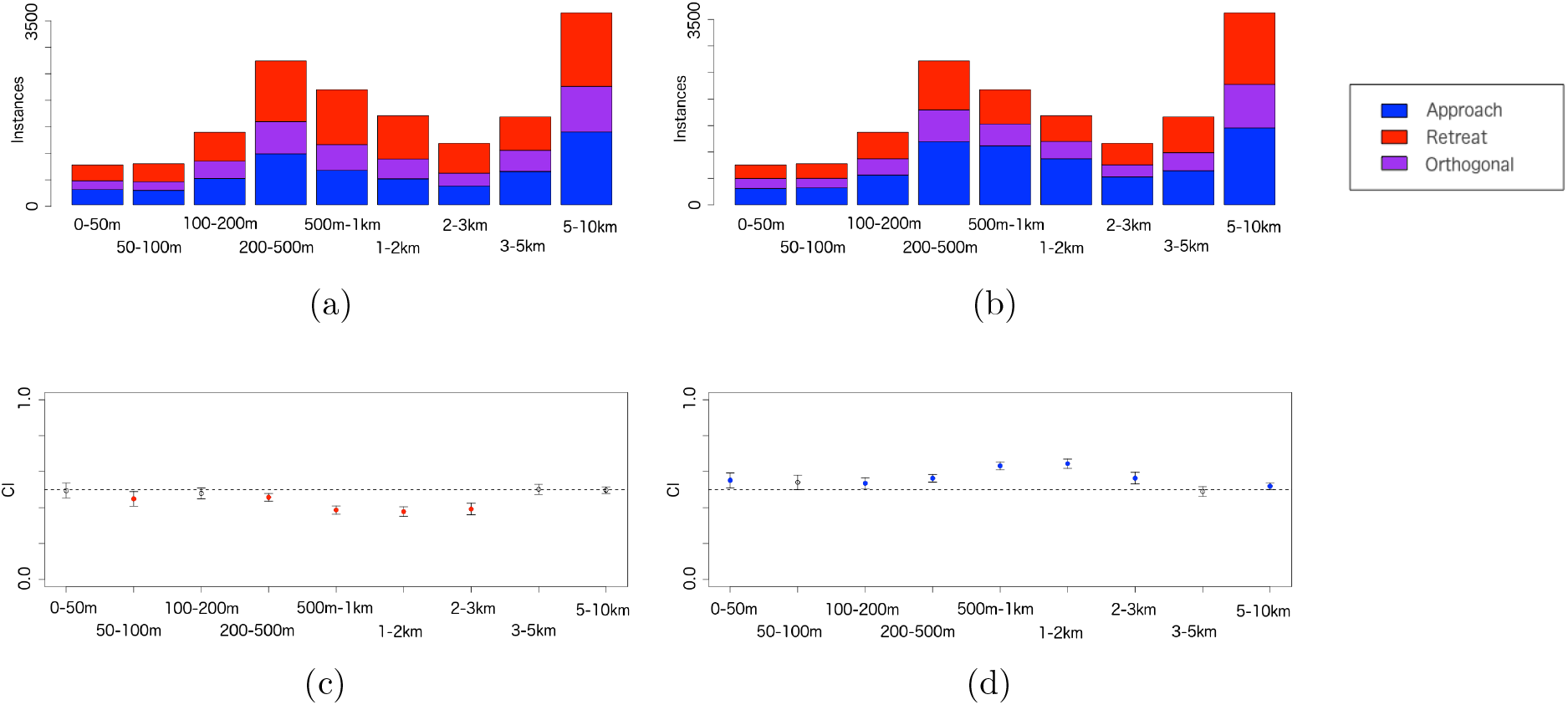
Individual behaviour classification for female (a) and male (b) and legend (right). Estimated CI, colored if providing statistically significant results, for female (c) and male (d). We can observe that the female presents mostly retreat behaviours (in red) while the male mostly approach behaviours (in blue).

#### 3.2.2. Dyadic behaviour

The results of our behavior analysis of our focal dyad is depicted in Fig. 11. We indicate statistically significant results (if above 0.25) in the upper part of the figure, corresponding to the different distance intervals, using the colors representing the different dyadic behaviour types. The table showing the results of the analysis is provided in our SOF. We observe that the statistically significant results correspond to pair behaviours of type 3(A,B) and 3(B,A). In particular, the most common pair behaviour corresponds to the female retreating while the male approaches (dyadic behaviour of type 3(B,A), A: female, B: male).

**Figure 11:**
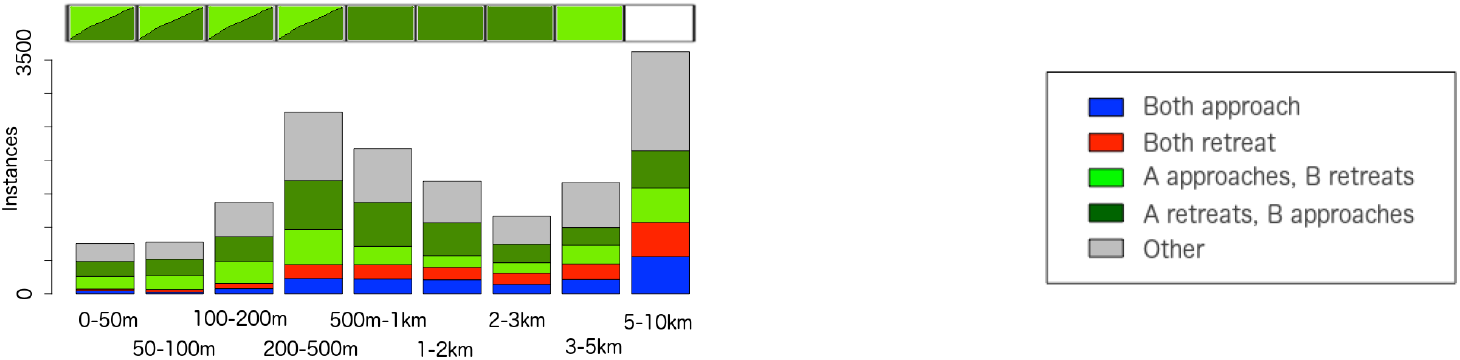
Analysis barplot for dyadic behaviour (right). The section above the barplot shows which of the four behaviours was significantly greater than 0.25 (when ignoring other), using the legend colors; two colored triangles were used if both behaviours of type 3 were statistically significant.

In Fig. 12 we show the results partitioned by season: hot-wet (Jan-Apr), cold-dry (May-Aug) and hot-dry (Sep-Dec). The hot-wet and hot-dry seasons contain significantly more female retreats from male interactions than the cold-dry season. Further, the hot dry period contains significantly more than random interactions at distances *<* 0.5 km where male retreats while female approaches, and female retreats while male approaches at distances up to 10 km (tables detailing these results are presented in the SOF, as well as the pair distance showing the various dyadic behaviour types over time).

**Figure 12:**
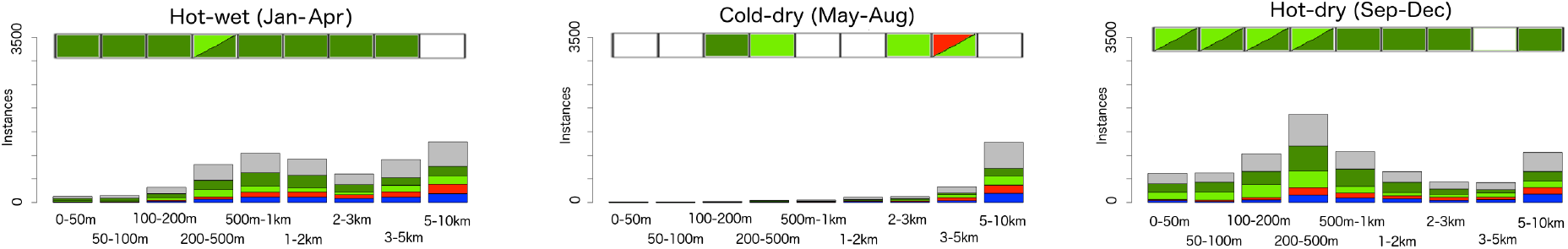
Analysis barplot for dyadic behaviour for hot-wet, cold-dry and hot-dry seasons. Same information above the barplots and same legend as Figure 11.

#### 3.2.3. Analysis of pair of interest - dyadic behaviour

From the GPS data of the 39 individuals, we extracted 53 pairs of interest which satisfied the criteria mentioned in Section 3.2. The 15-min frequency data were collected from 10 male individuals and we extracted 17 male-male dyads of interest, while the 30-min data were collected from 14 female individuals and we extracted 13 female-female dyads. The 20-min data were collected from 7 male and 8 female individuals: the 23 pairs chosen from this data were 11 male-male, 5 male-female and 7 female-female. The data did not provide information such as age of the individuals or any detailed relationship among them, except for the information regarding the breeding herd of the female-male dyad presented in previous sections. More informative data would have allowed a more extensive comparison among dyads.

We performed statistical analyses for the extracted dyads, and provide the percentage of dyads for which the results were statistically significant (below and above 0.25), for each distance interval, in Table 3. The detailed results for each data group are provided in our SOF. As shown in Table 3, the statistically significant results (above 0.25) are related to behaviours of type 3(A,B) and 3(B,A) (A approaches while B retreats and vice versa). In our SOF we also report the different percentage related to different sex-state pairs of the 20-min frequency, observing prevalent retreats for females and approaches for males in female-male dyads.

**Table 3:**
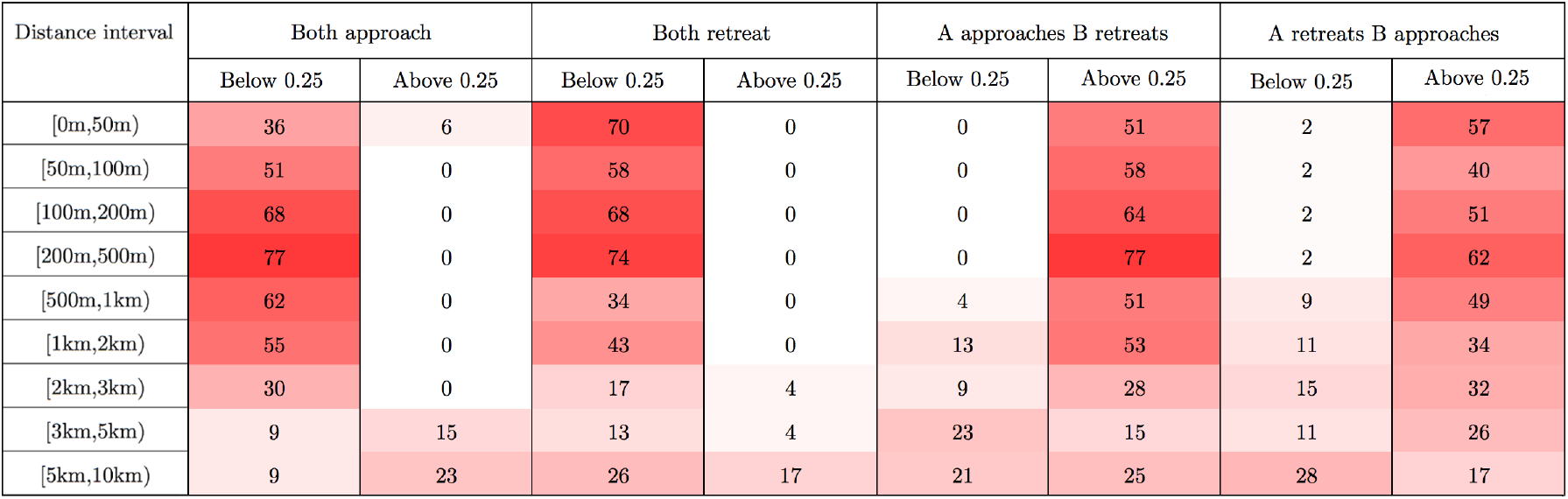
Analysis results for the pairs of interest. We observe that statistically significant results (above 0.25) are associated with behaviour of type 3 (one individual approaches while the other retreats). We colored the table cells according to the values: the darker cells contain higher values.

### 3.3. Extended analysis: female-male dyad

We performed an extended analysis that includes the absolute heading difference and relative speeds of the individuals for the focal female-male pair discussed in sections 3.2.1 and 3.2.2. We focus on the following eight behaviour modes, where A is the female and B the male:

a. A chasing B: A and B dyadic behaviour of type 3(A,B), speed of A greater than speed of B, similar heading
b. A following B: A and B dyadic behaviour of type 3(A,B), similar speed, similar heading
c. A escaping from B: A and B dyadic behaviour of type 3(B,A), speed of A greater than speed of B, similar heading
d. B chasing A: A and B dyadic behaviour of type 3(B,A), speed of B greater than speed of A, similar heading
e. B following A: A and B dyadic behaviour of type 3(B,A), similar speed, similar heading
f. B escaping from A: A and B dyadic behaviour of type 3(A,B), speed of B greater than speed of A, similar heading
g. A and B side by side: A and B dyadic behaviour of type 6, similar speed, similar heading
h. A and B approaching at a similar speed: A and B dyadic behaviour of type 1, similar speed, opposite headings

We categorize the relative speed of A (female) with respect to B (male) in proportional terms. Specifically, if the proportion exceeds 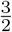, we categorize the speed of A as *greater* than B, while if below 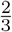 the speed of B is categorized as *greater* than A. Between these two proportions, we define that the two speeds as *similar*. In Fig. 13, we observe that the following behaviours (behaviours of type b and e) are the most common for the female-male pair of interest, especially for shorter distance between the individuals.

**Figure 13:**
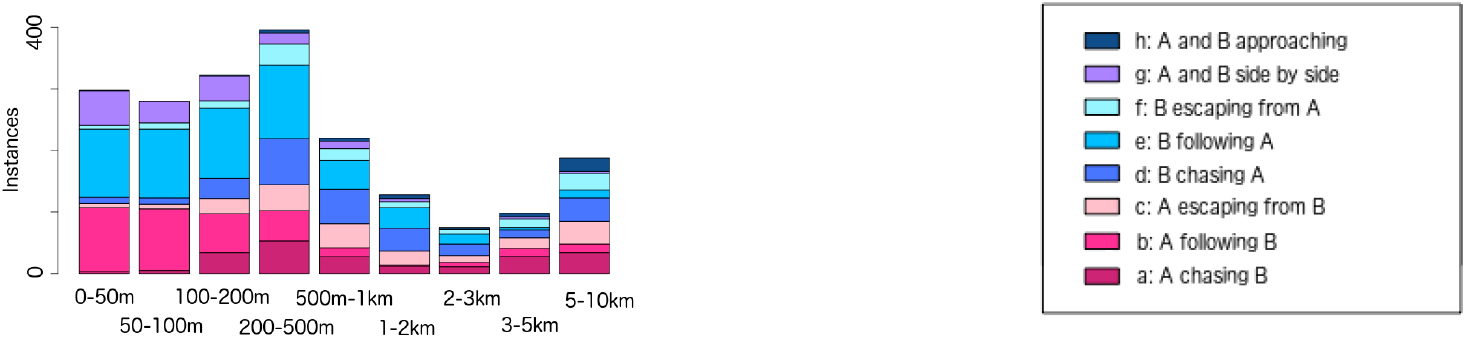
Results of the extended analysis. The legend is provided on the right, where A is the female and B the male. We observe that the following behaviours (type b and e) are the most frequent behaviours.

We also compared the frequencies of behaviours across diurnal and seasonal cycles, considering just data points for pair distances below 1 km. In Fig. 14 we depict the frequency of these behaviours at different times of the day (morning: 5-8am, evening: 5-8pm, night: midnight-3am) and for the different seasons. We observe that the frequency of interactions is relatively higher in the evening and substantially higher during the hot-dry season, but there are no outstanding differences among the different periods of the day and of the year in terms of the frequency of different behaviour types.

**Figure 14:**
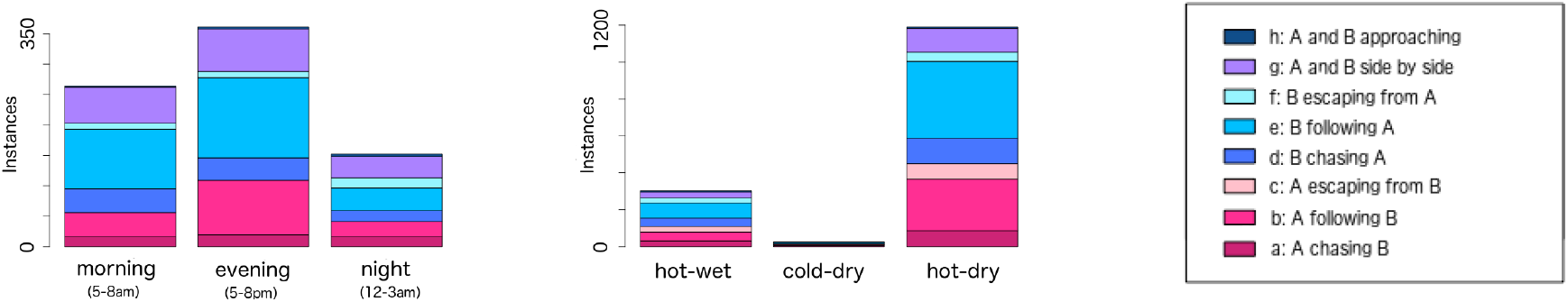
Extended analysis for dyadic behaviour types for different times of the day (left) and different seasons (center). Legend on the right. Also in this case, individual A corresponds to the female while individual B to the male.

## 4. Discussion

The literature contains several reviews on methods developed to study dyadic interactions. Miller (2012), for example, explores how spatially explicit simulated data can be used to analyse dynamic interactions between individuals. He uses five different techniques to quantify dynamic interactions based on GPS data of pairs of individuals, using both real (brown hyenas dyads) and simulated data. On the other hand, Long et al. (2014) evaluate the efficacy of eight different dynamic interaction indices, using both simulated and empirical data. They partition these indices into point-based measures, which typically study approach/retreat behaviour, and path-based measures, which look at movement behaviour. In particular, the index of proximity in space and the coefficient of association, example of point-based measures, look at the proportion of close simultaneous locations over the total number, while the correlation index and the dynamic interaction index, example of path-based measures, respectively evaluate the correlation and cohesiveness of movement segments that connect consecutive locations. Joo et al. (2018), following Long et al. (2014), assess the adequacy of 12 metrics introduced in previous works to assess specific aspects of joint-movement behaviour (two individuals move together for the total or a partial portion of their paths), focusing on proximity and coordination (synchrony) in direction and speed. The comparison is performed by building different scenarios with different levels of proximity and coordination to assess the ability of the metrics to capture various features. Some of these indices were used in Eriksen et al. (2009) to study both static and dynamic interactions among wolves and moose, and in Tosa et al. (2014) to examine dyads of female white-tailed deer within and between groups and to understand how those may related to pathogen transmission.

These existing techniques are based on the analysis of simultaneous locations, with or without a pair distance threshold and permutations of these, to perform different studies by comparing, for example, mean distance between simultaneous and permuted locations (coefficient of sociality), or using the overlap of home ranges as part of the analysis (Minta’s coefficient of interactions, half-weight association index). These dyadic approaches focus primarily on pairings of animals in space and time, with distance thresholds determining when dyads form and number of GPS fixes determining the length of time of the dyadic interaction. These techniques provide valuable information on which individuals form dyads and when they begin and end. Our methodology expands on this by introducing direction of movement and positions of individuals while interacting as a dyad. Our approach is also based on the analysis of simultaneous locations and can be defined as a point-based measure. However, it introduces the novelty of classifying angles related to the direction of movement and the position of the other individual in the dyad. After this classification, our methodology uses pair distance to group and analyse the results so that a deeper understanding of approach and retreat behaviours can be obtained. Additionally, through our extended dyadic behaviour analysis, we are able to evaluate frequencies of different dyadic behaviours that include differences in speed and absolute headings of the individuals.

A method for identifying various combinations of attraction, avoidance, and neutrality, using a framework of step-selection functions and both simulated and empirical data, is presented in Schlägel et al. (2019). In this work attraction and avoidance are considered in terms of an individual choosing locations that other individuals respectively use or do not use, while neutrality relates to ignoring the location use of others. In contrast to our approach, this method considers the distance to the home-range centers and not between pairs. Additionally, it is more suitable for evaluating short-range interactions (because the considered steps in the step-selection function might not reach far enough), while our method can apply to interactions that take place over any relevant distance. Other studies that include the relevance of sex in dyadic interactions is one on grizzly bear pairs where Doncaster’s measure (Doncaster, 1990) was used to understand sex-dependent and season-dependent interactions. Also, Spiegel et al. (2018) use a proximity-based social network method to analyse approach and retreat behaviours, with a specific focus on interaction rates between intrasex and intersex pairs.

Finally, a number of elephant interaction studies are reported in the literature. These include elephant-human interactions (Shaffer et al., 2019, Rossman et al., 2017), as well as zoo elephant tactile contact and proximity interactions (Bonaparte-Saller and Mench, 2018). Dyadic interactions among male African elephants (Goldenberg et al., 2014) and the social dynamics of female Asian elephants (de Silva et al., 2011) have also come under scrutiny, but these do not incorporate such movement elements as relative direction and speed that our methodology is specially designed to address.

## 5. Conclusions

In this paper we introduce a novel methodology to classify distance-dependent dyadic behaviour that allows us to extract approach and retreat behaviours by analysing animal GPS relocation data with matching frequencies and overlapping time periods. Our methodology was developed to classify distance-dependent behaviour, but it can be applied using other types of classification. We anticipate that our method will prove to be useful when applied to understanding behavioural persistence by analysing time series that capture the different behaviour types over time. In addition, it can be applied to investigating how different frequencies of data collection permit extraction of particular behaviours of interest at different spatio-temporal scales. It can also be embedded in studies that take various environmental factors, such as water sources or other types of resources, into account as covariates correlated with particular types of dyadic interactions.

## Acknowledgments and SOF File

The development of Nova, a precursor to Numerus Model Builder, was supported by NSF grant CNS-0939153 to Oberlin College and NSF-EEID grant 1617982 (PI: WMG). Wayne Getz is co-owner with two others of Numerus Inc., which provides a version of Numerus Modeler Builder online for free use. The collection of the GPS movement data were supported by grant NIH GM083863 (WMG, PI). In addition, LLV and JKB were funded by NIH 1R01GM117617-01 (PI: JKB). The supporting online file SOF, containing details of the model description and the analysis results is available at https://ludovicalv.github.io/PDFs/Elep_paper.pdf. The NMB models and the R code can be downloaded at https://ludovicalv.github.io/Dyadic_behaviour_method/.

## Data Accessibility Statement

We agree to archive the data associated with this manuscript should the manuscript be accepted. We intend to use a Zenodo repository.

## Authors’ contributions statement

LLV and WMG conceived the study and developed the methodology, LLV analysed the data, and LLV, JKB and WMG contributed to the writing of the manuscript.

