## Supplemental Material for "A relative-motion method for parsing spatio-temporal behaviour of dyads using GPS relocation data"

#### 1. Dyadic analysis

For the dyadic analysis, the procedure is similar to the individual analysis presented in the paper. The difference lies in the fact that we consider the simultaneous individual behaviour, and therefore we have a unique list to keep track of the dyadic behaviour types. This list, indicated by $M_D$, is composed of $n$ 5-dimensional vectors, initially populated with zeros. The procedure is the same as for the individual analysis up to the pair distance calculation. If this distance is below our maximum threshold for considering dyadic interactions, then the function $f_{cd}$ is used to categorize the dyadic behaviour in terms of the angles diffA and diffB. The update function will then update the list $M_D$, according to the dyadic behaviour type and the corresponding distance interval.

**Algorithm 1:** Methodology implementation

```
Input 𝒯_A, 𝒯_B, ℐ
M_D = list()
for i in {1, ..., n} do
    M_D[i] = v_0(5)
I_AB = T_A ∩ T_B
for t in I_AB do
    headA = f(A(t), A(t+1)), headB = f(B(t), B(t+1))
    dirAB = f(A(t), B(t)), dirBA = f(B(t), A(t))
    diffA = |headA - dirAB|, diffB = |headB - dirBA|
    d_AB = d((A(t), B(t))
    if ∃i : d_AB ∈ I_i then
        m_D = f_cd(diffA,diffB)
        M_D[i] = u(m_D, M_D[i])
return M_A, M_B
```

### 2. Extended analysis

For the extended analysis, we extract eight behaviour types, by considering also the heading difference and the relative speed. We set the speed proportion limits $p_l$ and $p_u$, and a list $M_F$ to keep track of the counts. Given two individuals A and B, the speed proportion limits $p_l$ and $p_u$ are defined such that if the speed proportion $s_p$ is above $p_u$, then speed of A is considered *greater* than speed of B. If $s_p$ lies between $p_l$ and $p_u$, then the speed is considered *similar*, while if the proportion is below $p_l$, then the speed of B is considered *greater* than the speed of A.

After the calculation of the usual angles, we evaluate also the heading difference diffH, the pair distance $d_{AB}$, the individual speed $s_A$ and $s_B$ (as ratio between the distance $\Delta l$ and time interval $\Delta t$) and the proportion of the two individual speeds $s_p$. If the pair distance $d_{AB}$ lies within a chosen distance interval $I_i$, then the function $f_{cf}$ classify the pair behaviour type considering all the necessary inputs and the function $u$ subsequently updates the vector count.

**Algorithm 2:** Methodology implementation

```
Input 𝒯_A, 𝒯_B, ℐ, d_M, p_l, p_u
M_F = list()
for i in {1,...,n} do
    M_F[i] = v_0(8)
I_AB = T_A ∩ T_B
for t in I_AB do
    headA = f(A(t), A(t+1)), headB = f(B(t), B(t+1))
    dirAB = f(A(t), B(t)), dirBA = f(B(t), A(t))
    diffA = |headA - dirAB|, diffB = |headB - dirBA|
    diffH = |headA - headB|
    d_AB = d(A(t), B(t))
    s_A = d(A(t),A(t+1)) / Δt_A, s_B = d(B(t),B(t+1)) / Δt_B
    s_p = s_A / s_B
    if ∃i : d_AB ∈ I_i then
        m_F = f_cf(diffA,diffB,diffH, s_p, p_l, p_u)
        M_F[i] = u(m_F, M_F[i])
return M_F
```

#### 2.1. *Extended analysis: dyadic movement modes*

The following table combines the information related to the classified dyadic movement modes, relative speed and individual heading analysis. We provide a description of behaviours of interest, used in Section 7.4 of the main paper.

Dyadic behaviour:

1. both individuals approach each other
2. both individuals retreat from each other
3. one individual approaches while the other individual retreats
4. one individual moves orthogonally, the other approaches
5. one individual moves orthogonally, the other retreats
6. both individuals move orthogonally

To each of these six dyadic modes, we assign an additional designator: a.) similar speed, b.) A faster than B, 3.) B faster than A. This yields a total of 18 dyadic movement modes: 1a, 1b, ..., 6c. Considering also the heading analysis allows us to extract various behaviours of interest, that we describe in the following table.

| Modes | Dyadic behaviour | Speed analysis | Description |
|---|---|---|---|
| 1a | Both individuals approach each other | Similar | With opposite individual headings, A and B approaching at a similar speed |
| 1b | Both individuals approach each other | A faster than B | |
| 1c | Both individuals approach each other | B faster than A | |
| 2a | Both individuals retreat from each other | Similar | |
| 2b | Both individuals retreat from each other | A faster than B | |
| 2c | Both individuals retreat from each other | B faster than A | |
| 3a | One individual (A) approaches while the other individual (B) retreats | Similar | With similar individual heading, following behaviour |
| 3b | One individual (A) approaches while the other individual (B) retreats | A faster than B | With similar individual heading, A chasing B |
| 3c | One individual approaches (A) while the other individual (B) retreats | B faster than A | With similar individual heading, B escaping from A |

| Modes | Behaviour | Speed | Description |
|---|---|---|---|
| 4a | One individual (A) moves orthogonally, the other (B) approaches | Similar | |
| 4b | One individual moves orthogonally, the other approaches | A faster than B | |
| 4c | One individual moves orthogonally, the other approaches | B faster than A | |
| 5a | One individual moves orthogonally, the other retreats | Similar | |
| 5b | One individual moves orthogonally, the other retreats | A faster than B | |
| 5c | One individual moves orthogonally, the other retreats | B faster than A | |
| 6a | Both individuals move orthogonally | Similar | With similar individual heading, side by side movement |
| 6b | Both individuals move orthogonally | A faster than B | |
| 6c | Both individuals move orthogonally | B faster than A | |

Note that we considered speed and heading difference in the analysis to extract meaningful behaviours (e.g. following, side by side). However, given all the possible combinations, some resulting behaviours might not be meaningful. For example, "both individuals approaching, similar speed, similar heading" is not a possible behaviour, since both individual cannot be moving towards each other and have similar absolute headings.

### 3. Simulated data

In this section, we describe the relative-motion, biased random-walk (RM-BRW) models implemented in Numerus Model Builder (NMB) and used to generate simulated data. We provide the description of the movement model for individual A of pair (A,B), since the behaviour of individual B is the same as the one of A, just with a different direction (B approaching/retreating from A instead of A approaching/retreating from B).

Given the initial position $(x_0, y_0)$ of individual A, the location coordinates are updated as follows:

$$
\begin{aligned}
x_{t+1} &= x_t + s_t \cos\theta_t \\
y_{t+1} &= y_t + s_t \sin\theta_t
\end{aligned}
$$

where $s_t$ is the step length and $\theta_t$ is the absolute heading. The step length is drawn from the uniform distribution:

$$s_t \sim \text{UNIFORM}(s_{\min}, s_{\max})$$

while the absolute heading is drawn from different distributions, which are described later.

#### *3.1. Distance-dependent behaviour (attraction and repulsion circumferences)*

In the first model, these distributions are distance-dependent with noise introduced using the coefficient $\rho \in [0, 1]$ and attracting and repulsing circles of radii $d_R$ and $d_I$ (Fig. 1):

$$\theta_{t+1} \sim \begin{cases} \text{UNIFORM}(\theta_{A\to B} - (1-\rho)\frac{\pi}{2}, \theta_{A\to B} + (1-\rho)\frac{\pi}{2}) & \text{if case 1} \\ \text{UNIFORM}(-\theta_{A\to B} - (1-\rho)\frac{\pi}{2}, -\theta_{A\to B} + (1-\rho)\frac{\pi}{2}) & \text{if case 2} \\ \text{UNIFORM}(-\pi, \pi) & \text{otherwise} \end{cases}$$

where $\theta_{A\to B}$ is the heading from $A(t)$ to $B(t)$, $\rho \in [0, 1]$ and:

- case 1: $d_R < d_{AB} < d_I$ and UNIFORM(0,1) $< p_{\text{eff}}$
- case 2: $d_{AB} < d_R$ and UNIFORM(0,1) $< p_{\text{eff}}$

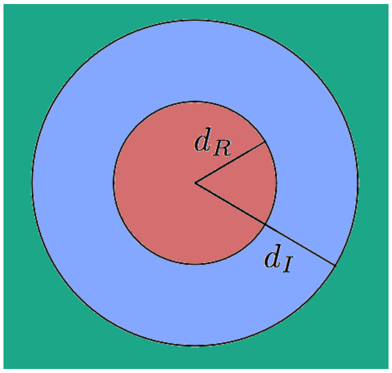


Figure 1: Area of repulsion (red), attraction (blue) and indifference (green), depending on $d_R$ and $d_I$, around one individual.

The first case represents approach: the individuals are at a distance between the repulsion distance $d_R$ and the indifference distance $d_I$, while the second case represents repulsion. These behaviours happen with probability $p_{\text{eff}}$, otherwise the movement is random.

#### *3.2. Time-dependent behaviour*

In the second model, the distributions used to drawn $\theta_{t+1}$ are a function of time. Given the period $\omega$ and the functions $f_1$ and $f_2$:

$$f_1(t) = \sin\left(\frac{2\pi t}{\omega}\right)$$

$$f_2(t) = \sin\left(\frac{2\pi t}{\frac{\omega}{2}}\right)$$

The value of the absolute heading is drawn as follow:

$$\theta_{t+1} \sim \begin{cases} \text{UNIFORM}(\theta_{A\to B} - (1-\rho)\frac{\pi}{2}, \theta_{A\to B} + (1-\rho)\frac{\pi}{2}) & \text{if case 1} \\ \text{UNIFORM}(-\theta_{A\to B} - (1-\rho)\frac{\pi}{2}, -\theta_{A\to B} + (1-\rho)\frac{\pi}{2}) & \text{if case 2} \\ \text{UNIFORM}(-\pi, \pi) & \text{otherwise} \end{cases}$$

where:

- case 1: $f_1(t) > 0$ and $f_2(t) > 0$
- case 2: $f_1(t) > 0$ and $f_2(t) < 0$

The first case represents A approaching B, the second case A retreating from B while the third case is a random walk, without a preferred direction.

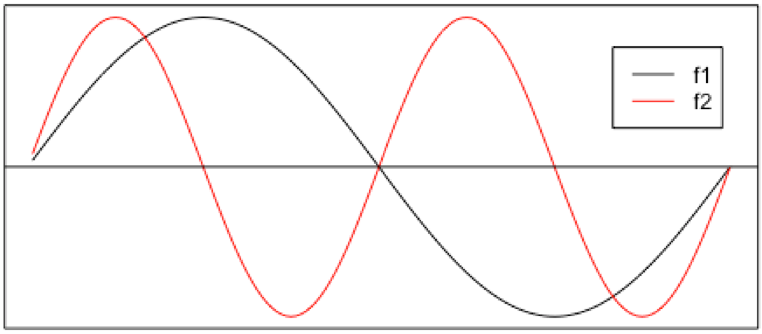


Figure 2: One period of $f_1$ (black) and 2 periods of $f_2$ (red).

Parameters used in the simulations:

| Name | Value |
|---|---|
| $s_{\text{min}}$ | 5 |
| $s_{\text{max}}$ | 6 |
| $\omega$ | 144 |
| $\rho$ | 0.5 |
| $d_I$ | 60 |
| $d_R$ | 30 |
| $p_{\text{eff}}$ | 0.5 |

Note that the choice of parameters was arbitrary. Other values can be selected, depending on what aspects of the model are being evaluated or tested.

### 4. Results

In the results reported here, we used the Euclidean distance to calculate the dyadic distance in units and we scaled the coordinates to be able to use the function `bearing` to calculate the various angles.

#### *4.1. Individual behaviour: individual B*

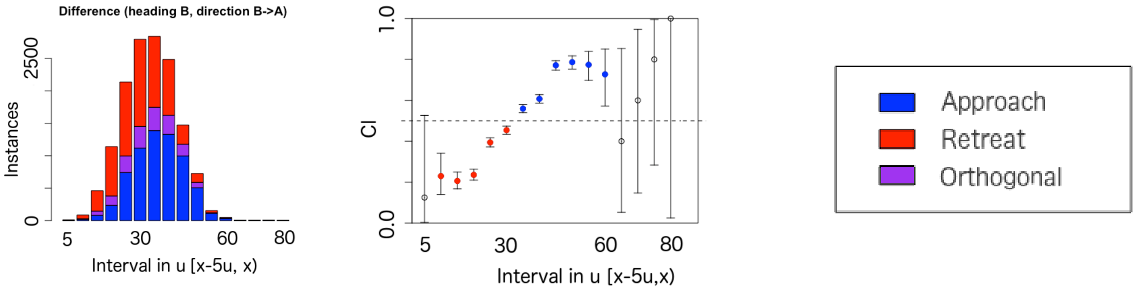


Figure 3: Barplot for individual B (left). Estimated confidence intervals (CI) for individual B, colored if statistically significant result (center) according to the legend (right).

#### *4.2. Dyadic behaviour: pair (A,B)*

We indicate behaviours of type 3(A,B) and 3(B,A) using the numbers 3 and 4 respectively.

| Pair | Distance interval (units) | Total | Count | Type | Lower CI | Upper CI | Retreat | Approach |
|---|---|---|---|---|---|---|---|---|
| A and B | [0,5) | 7 | 0 | 1 | 0 | 0.4096 | | |
| A and B | | 7 | 3 | 2 | 0.099 | 0.8159 | | |
| A and B | | 7 | 3 | 3 | 0.099 | 0.8159 | | |
| A and B | | 7 | 1 | 4 | 0.0036 | 0.5787 | | |
| A and B | [5,10) | 60 | 5 | 1 | 0.0276 | 0.1839 | ✓ | |
| A and B | | 60 | 30 | 2 | 0.3681 | 0.6319 | | ✓ |
| A and B | | 60 | 16 | 3 | 0.1607 | 0.3966 | | |
| A and B | | 60 | 9 | 4 | 0.071 | 0.2657 | | |
| A and B | [10,15) | 347 | 17 | 1 | 0.0288 | 0.0773 | ✓ | |
| A and B | | 347 | 224 | 2 | 0.5927 | 0.6959 | | ✓ |
| A and B | | 347 | 54 | 3 | 0.1191 | 0.1981 | ✓ | |
| A and B | | 347 | 52 | 4 | 0.114 | 0.1918 | ✓ | |
| A and B | [15,20) | 880 | 53 | 1 | 0.0454 | 0.078 | ✓ | |
| A and B | | 880 | 530 | 2 | 0.5691 | 0.6348 | | ✓ |
| A and B | | 880 | 142 | 3 | 0.1377 | 0.1874 | ✓ | |
| A and B | | 880 | 155 | 4 | 0.1515 | 0.2029 | ✓ | |
| A and B | [20,25) | 1628 | 385 | 1 | 0.216 | 0.2579 | | |
| A and B | | 1628 | 696 | 2 | 0.4033 | 0.452 | | ✓ |
| A and B | | 1628 | 278 | 3 | 0.1528 | 0.1899 | ✓ | |
| A and B | | 1628 | 269 | 4 | 0.1475 | 0.1842 | ✓ | |
| A and B | [25,30) | 2148 | 612 | 1 | 0.2659 | 0.3045 | | ✓ |
| A and B | | 2148 | 801 | 2 | 0.3524 | 0.3938 | | ✓ |
| A and B | | 2148 | 377 | 3 | 0.1596 | 0.1923 | ✓ | |
| A and B | | 2148 | 358 | 4 | 0.1511 | 0.1831 | ✓ | |
| A and B | [30,35) | 2143 | 861 | 1 | 0.3809 | 0.4229 | | ✓ |
| A and B | | 2143 | 608 | 2 | 0.2647 | 0.3033 | | ✓ |
| A and B | | 2143 | 334 | 3 | 0.1407 | 0.1719 | ✓ | |
| A and B | | 2143 | 340 | 4 | 0.1434 | 0.1748 | ✓ | |
| A and B | [35,40) | 1924 | 833 | 1 | 0.4107 | 0.4554 | | ✓ |
| A and B | | 1924 | 422 | 2 | 0.201 | 0.2385 | ✓ | |
| A and B | | 1924 | 329 | 3 | 0.1544 | 0.1886 | ✓ | |
| A and B | | 1924 | 340 | 4 | 0.1599 | 0.1945 | ✓ | |
| A and B | [40,45) | 1125 | 683 | 1 | 0.5779 | 0.6358 | | ✓ |
| A and B | | 1125 | 79 | 2 | 0.056 | 0.0868 | ✓ | |
| A and B | | 1125 | 175 | 3 | 0.1349 | 0.1781 | ✓ | |
| A and B | | 1125 | 188 | 4 | 0.1458 | 0.1902 | ✓ | |
| A and B | [45,50) | 558 | 337 | 1 | 0.562 | 0.6448 | | ✓ |
| A and B | | 558 | 20 | 2 | 0.022 | 0.0548 | ✓ | |
| A and B | | 558 | 92 | 3 | 0.135 | 0.1983 | ✓ | |
| A and B | | 558 | 109 | 4 | 0.1632 | 0.2307 | ✓ | |
| A and B | [50,55) | 132 | 85 | 1 | 0.5559 | 0.7253 | | ✓ |
| A and B | | 132 | 4 | 2 | 0.0083 | 0.0758 | ✓ | |
| A and B | | 132 | 26 | 3 | 0.1329 | 0.2751 | | |
| A and B | | 132 | 17 | 4 | 0.0768 | 0.1982 | ✓ | |
| A and B | [55,60) | 35 | 22 | 1 | 0.4492 | 0.7853 | | ✓ |
| A and B | | 35 | 1 | 2 | 7e-04 | 0.1492 | ✓ | |
| A and B | | 35 | 8 | 3 | 0.1042 | 0.4014 | | |
| A and B | | 35 | 4 | 4 | 0.032 | 0.2674 | | |
| A and B | [60,65) | 5 | 1 | 1 | 0.0051 | 0.7164 | | |
| A and B | | 5 | 1 | 2 | 0.0051 | 0.7164 | | |
| A and B | | 5 | 2 | 3 | 0.0527 | 0.8534 | | |
| A and B | | 5 | 1 | 4 | 0.0051 | 0.7164 | | |
| A and B | [65,70) | 5 | 1 | 1 | 0.0051 | 0.7164 | | |
| A and B | | 5 | 0 | 2 | 0 | 0.5218 | | |
| A and B | | 5 | 2 | 3 | 0.0527 | 0.8534 | | |
| A and B | | 5 | 2 | 4 | 0.0527 | 0.8534 | | |
| A and B | [70,75) | 5 | 0 | 1 | 0 | 0.5218 | | |
| A and B | | 5 | 0 | 2 | 0 | 0.5218 | | |
| A and B | | 5 | 1 | 3 | 0.0051 | 0.7164 | | |
| A and B | | 5 | 4 | 4 | 0.2836 | 0.9949 | | ✓ |
| A and B | [75,80) | 1 | 1 | 1 | 0.025 | 1 | | |
| A and B | | 1 | 0 | 2 | 0 | 0.975 | | |
| A and B | | 1 | 0 | 3 | 0 | 0.975 | | |
| A and B | | 1 | 0 | 4 | 0 | 0.975 | | |

#### *4.3. Results for time-dependent RM-BRW model*

In this section we show the results of the individual analysis obtained by subdividing our 10-day simulation data depending on the time of the day (first quarter, second quarter and second half of the day), considering dyadic distance below 100 units. In Figure 4 we show the results related to the first quarter of the day (first row), the results for the analysis of data from the second quarter of the day (second row) and the results corresponding to the second half of the day (third row). As expected, we observe that in the first quarter of the day the individuals mostly show approach behaviour (in blue) while in the second quarter of the day they show mostly retreat behaviour (in red). In the second half of the day, when the individuals move independently, we observe a balanced mixture of the different behaviour types, without a prevalent one. In Figure 5 we depict the results of the dyadic analysis, divided depending on the time of the day. Also in this case, the analysis captured the different dyadic behaviour (both approaching, both retreating) for the first 2 quarters of the day, while the second half presents the various dyadic behaviour types.

Note that we binned the results for distance below 100. However, in this case, the dyadic distance was not bounded above as before, so there are more data for distance above 100 (maximum 4002 units), which we show in Figures 6 and 7. These results show the same time-dependent patterns as in the previous analysis, which are captured by our methodology, using subdivided data.

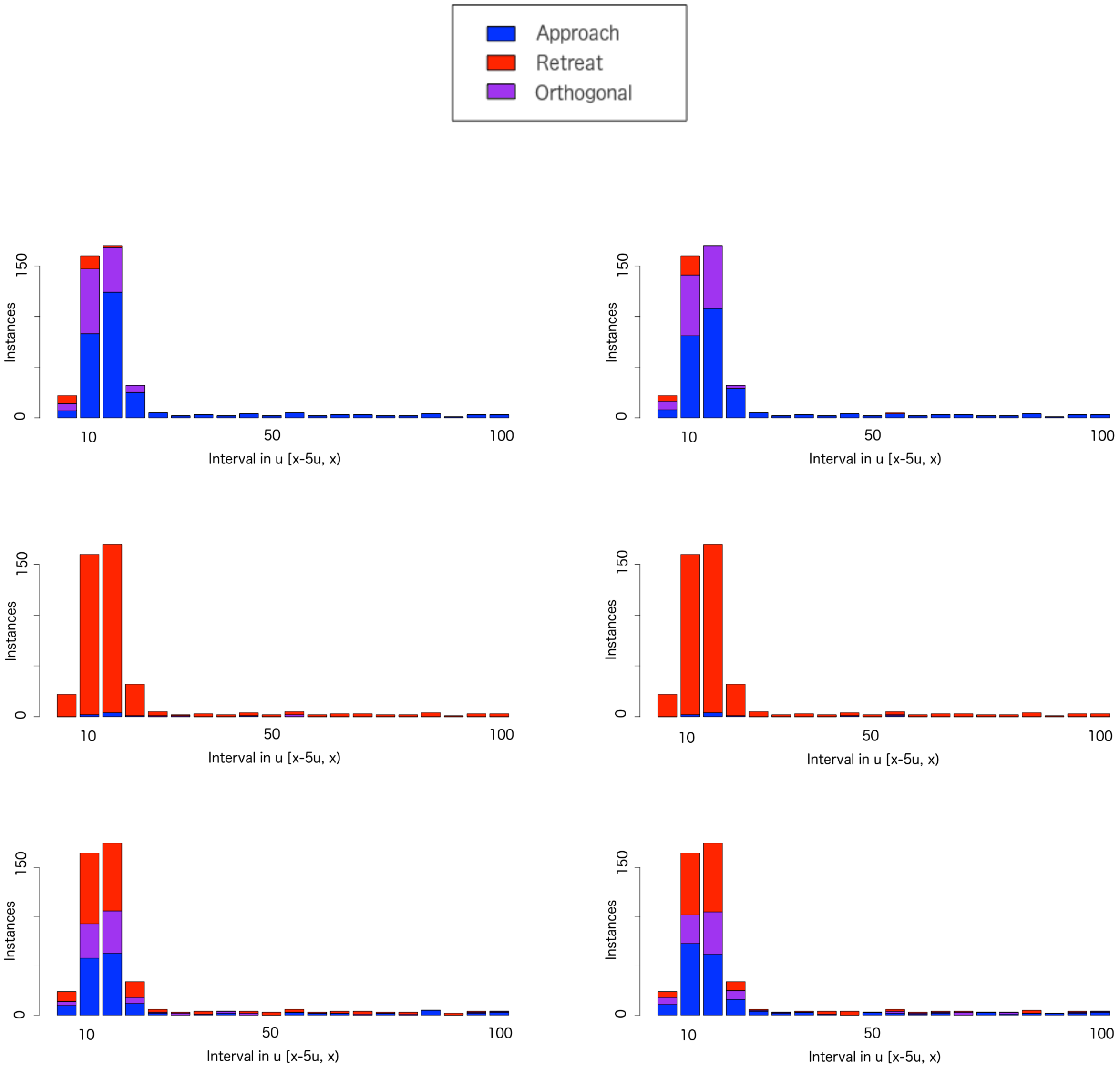


Figure 4: Results of individual analysis for individual A (left) and B (center), for the time-dependent RM-BRW model, for different times of the day. First row: first quarter of the day, second row: second quarter of the day, third row: second half of the day. Legend on the top.

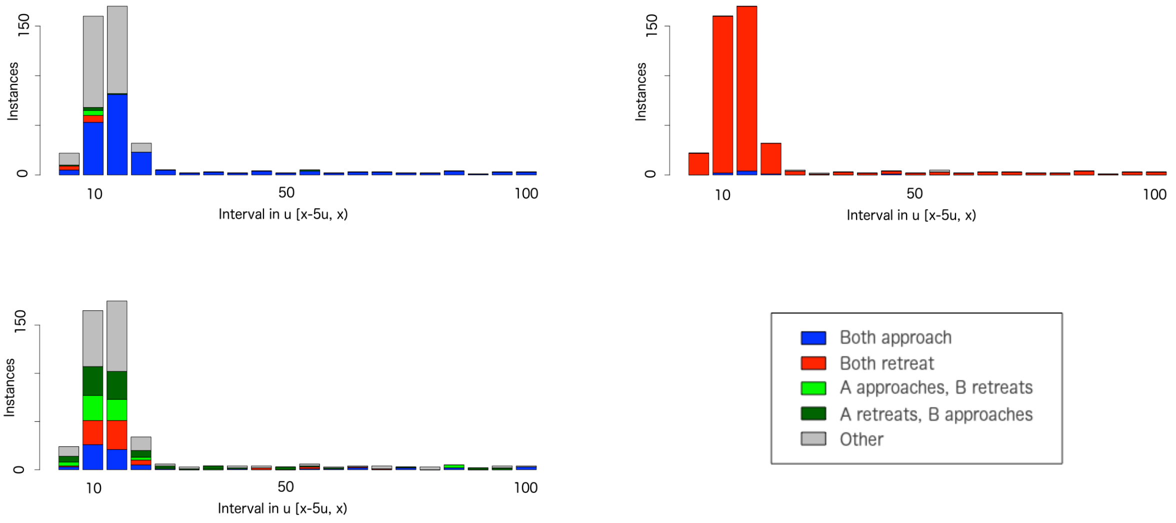


Figure 5: Results of dyadic analysis for individual A and B, for the time-dependent RM-BRW model, for different times of the day. From the top left: first quarter of the day, second quarter of the day, second half of the day, legend.

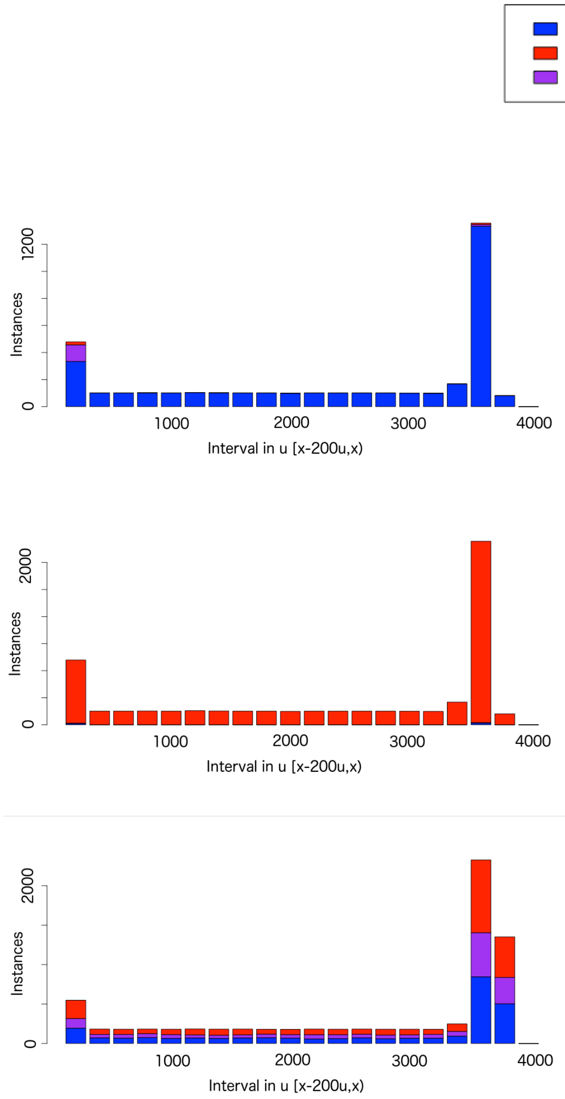


Figure 6: Results of individual analysis for individual A (left) and B (center), for the time-dependent RM-BRW model, for different times of the day. First row: first quarter of the day, second row: second quarter of the day, third row: second half of the day. Legend on the top.

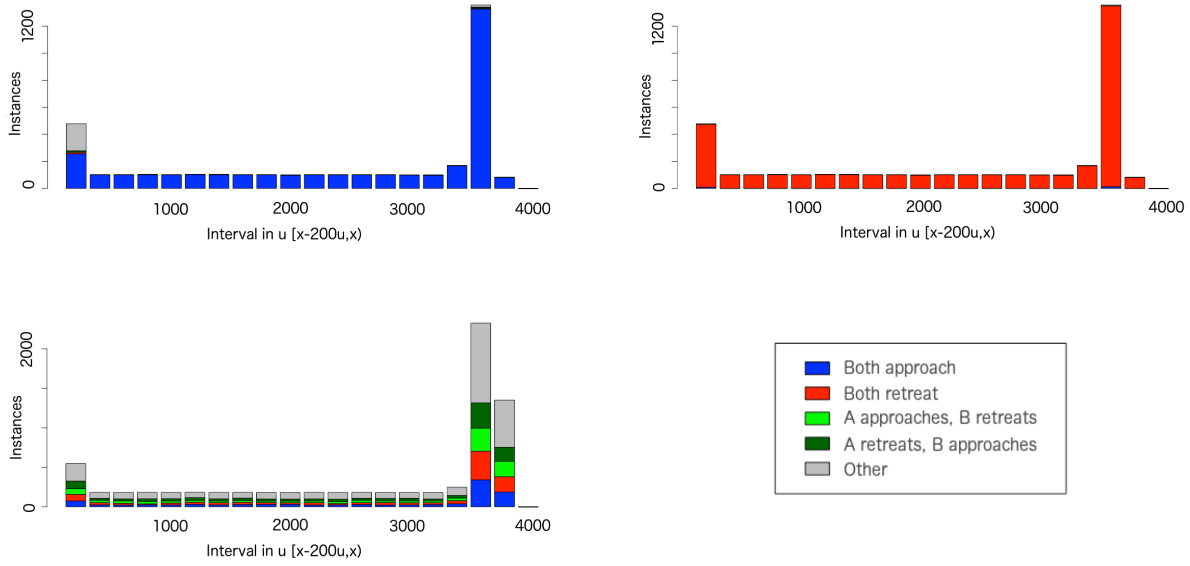


Figure 7: Results of dyadic analysis for individual A and B, for the time-dependent RM-BRW model, for different times of the day. From the top left: first quarter of the day, second quarter of the day, second half of the day, legend.

### 5. Empirical data

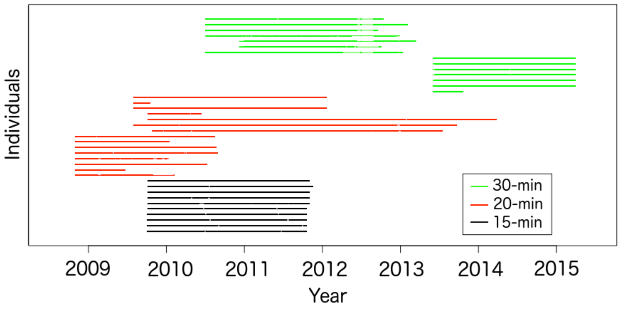


Figure 8: Time line of data collection for each individual and for each different time interval.

#### *5.1. Dyadic behaviour: pair (female,male)*

As before, we indicate behaviours of type 3(A,B) and 3(B,A) with numbers 3 and 4. Statistically significant results are marked using the symbol ✓. Letter A represents the female while letter B represents the male.

| Pair | Distance interval (m) | Total | Count | Type | Lower CI | Upper CI | Sign. diff. |
|---|---|---|---|---|---|---|---|
| A and B | [0,50) | 485 | 49 | 1 | 0.0757 | 0.1314 | ✓ |
| A and B | | 485 | 26 | 2 | 0.0353 | 0.0776 | ✓ |
| A and B | | 485 | 185 | 3 | 0.338 | 0.4263 | ✓ |
| A and B | | 485 | 225 | 4 | 0.4188 | 0.5094 | ✓ |
| A and B | [50,100) | 522 | 27 | 1 | 0.0344 | 0.0744 | ✓ |
| A and B | | 522 | 41 | 2 | 0.057 | 0.105 | ✓ |
| A and B | | 522 | 207 | 3 | 0.3543 | 0.44 | ✓ |
| A and B | | 522 | 247 | 4 | 0.4296 | 0.517 | ✓ |
| A and B | [100,200) | 857 | 80 | 1 | 0.0747 | 0.1148 | ✓ |
| A and B | | 857 | 78 | 2 | 0.0726 | 0.1123 | ✓ |
| A and B | | 857 | 328 | 3 | 0.3501 | 0.4162 | ✓ |
| A and B | | 857 | 371 | 4 | 0.3994 | 0.4668 | ✓ |
| A and B | [200,500) | 1700 | 230 | 1 | 0.1194 | 0.1525 | ✓ |
| A and B | | 1700 | 208 | 2 | 0.1071 | 0.1389 | ✓ |
| A and B | | 1700 | 527 | 3 | 0.2881 | 0.3326 | ✓ |
| A and B | | 1700 | 735 | 4 | 0.4086 | 0.4563 | ✓ |
| A and B | [500,1000) | 1369 | 222 | 1 | 0.143 | 0.1828 | ✓ |
| A and B | | 1369 | 214 | 2 | 0.1375 | 0.1767 | ✓ |
| A and B | | 1369 | 276 | 3 | 0.1806 | 0.2239 | ✓ |
| A and B | | 1369 | 657 | 4 | 0.4531 | 0.5068 | ✓ |
| A and B | [1000,2000) | 1064 | 211 | 1 | 0.1747 | 0.2236 | ✓ |
| A and B | | 1064 | 187 | 2 | 0.1533 | 0.2 | ✓ |
| A and B | | 1064 | 176 | 3 | 0.1436 | 0.1891 | ✓ |
| A and B | | 1064 | 490 | 4 | 0.4303 | 0.491 | ✓ |
| A and B | [2000,3000) | 742 | 143 | 1 | 0.1649 | 0.223 | ✓ |
| A and B | | 742 | 170 | 2 | 0.1993 | 0.2611 | |
| A and B | | 742 | 153 | 3 | 0.1776 | 0.2371 | ✓ |
| A and B | | 742 | 276 | 4 | 0.3371 | 0.4079 | ✓ |
| A and B | [3000,5000) | 997 | 216 | 1 | 0.1914 | 0.2435 | ✓ |
| A and B | | 997 | 232 | 2 | 0.2068 | 0.2602 | |
| A and B | | 997 | 280 | 3 | 0.2531 | 0.3099 | ✓ |
| A and B | | 997 | 269 | 4 | 0.2425 | 0.2985 | |
| A and B | [5000,10000) | 2144 | 559 | 1 | 0.2422 | 0.2799 | |
| A and B | | 2144 | 512 | 2 | 0.2209 | 0.2574 | |
| A and B | | 2144 | 517 | 3 | 0.2232 | 0.2598 | |
| A and B | | 2144 | 556 | 4 | 0.2409 | 0.2784 | |

#### 5.2. *Hot-wet season*

| Pair | Distance interval (m) | Total | Count | Type | Lower CI | Upper CI | Sign. diff. |
|---|---|---|---|---|---|---|---|
| A and B | [0,50) | 82 | 6 | 1 | 0.0273 | 0.1525 | ✓ |
| A and B | | | 7 | 2 | 0.035 | 0.168 | ✓ |
| A and B | | | 28 | 3 | 0.2403 | 0.4545 | |
| A and B | | | 41 | 4 | 0.3875 | 0.6125 | ✓ |
| A and B | [50,100) | 89 | 9 | 1 | 0.0473 | 0.1833 | ✓ |
| A and B | | | 8 | 2 | 0.0396 | 0.1695 | ✓ |
| A and B | | | 30 | 3 | 0.2403 | 0.4451 | |
| A and B | | | 42 | 4 | 0.3651 | 0.5806 | ✓ |
| A and B | [100,200) | 186 | 24 | 1 | 0.0845 | 0.1859 | ✓ |
| A and B | | | 25 | 2 | 0.0889 | 0.192 | ✓ |
| A and B | | | 54 | 3 | 0.2262 | 0.3612 | |
| A and B | | | 83 | 4 | 0.3735 | 0.5207 | ✓ |
| A and B | [200,500) | 469 | 67 | 1 | 0.1125 | 0.1778 | ✓ |
| A and B | | | 53 | 2 | 0.0858 | 0.1452 | ✓ |
| A and B | | | 156 | 3 | 0.2901 | 0.3773 | ✓ |
| A and B | | | 193 | 4 | 0.3666 | 0.4576 | ✓ |
| A and B | [500,1000) | 632 | 118 | 1 | 0.1571 | 0.2193 | ✓ |
| A and B | | | 102 | 2 | 0.1336 | 0.1924 | ✓ |
| A and B | | | 131 | 3 | 0.1763 | 0.241 | ✓ |
| A and B | | | 281 | 4 | 0.4054 | 0.4843 | ✓ |
| A and B | [1000,2000) | 576 | 118 | 1 | 0.1726 | 0.2402 | ✓ |
| A and B | | | 106 | 2 | 0.1532 | 0.2181 | ✓ |
| A and B | | | 85 | 3 | 0.1196 | 0.1792 | ✓ |
| A and B | | | 267 | 4 | 0.4222 | 0.5052 | ✓ |
| A and B | [2000,3000) | 377 | 84 | 1 | 0.1818 | 0.2682 | |
| A and B | | | 82 | 2 | 0.1769 | 0.2626 | |
| A and B | | | 59 | 3 | 0.1213 | 0.1972 | ✓ |
| A and B | | | 152 | 4 | 0.3533 | 0.4546 | ✓ |
| A and B | [3000,5000) | 529 | 117 | 1 | 0.1865 | 0.259 | |
| A and B | | | 111 | 2 | 0.1759 | 0.2471 | ✓ |
| A and B | | | 136 | 3 | 0.2203 | 0.2966 | |
| A and B | | | 165 | 4 | 0.2726 | 0.3533 | ✓ |
| A and B | [5000,10000) | 769 | 186 | 1 | 0.212 | 0.2737 | |
| A and B | | | 200 | 2 | 0.2294 | 0.2926 | |
| A and B | | | 181 | 3 | 0.2058 | 0.267 | |
| A and B | | | 202 | 4 | 0.2319 | 0.2953 | |

### 5.3. *Cold-dry season*

| Pair | Distance interval (m) | Total | Count | Type | Lower CI | Upper CI | Sign. diff. |
|---|---|---|---|---|---|---|---|
| A and B | [0,50) | 6 | 0 | 1 | 0 | 0.4593 | |
| A and B | | | 0 | 2 | 0 | 0.4593 | |
| A and B | | | 4 | 3 | 0.2228 | 0.9567 | |
| A and B | | | 2 | 4 | 0.0433 | 0.7772 | |
| A and B | [50,100) | 6 | 0 | 1 | 0 | 0.4593 | |
| A and B | | | 0 | 2 | 0 | 0.4593 | |
| A and B | | | 3 | 3 | 0.1181 | 0.8819 | |
| A and B | | | 3 | 4 | 0.1181 | 0.8819 | |
| A and B | [100,200) | 14 | 1 | 1 | 0.0018 | 0.3387 | |
| A and B | | | 2 | 2 | 0.0178 | 0.4281 | |
| A and B | | | 3 | 3 | 0.0466 | 0.508 | |
| A and B | | | 8 | 4 | 0.2886 | 0.8234 | ✓ |
| A and B | [200,500) | 36 | 5 | 1 | 0.0467 | 0.295 | |
| A and B | | | 2 | 2 | 0.0068 | 0.1866 | ✓ |
| A and B | | | 17 | 3 | 0.3041 | 0.6451 | ✓ |
| A and B | | | 12 | 4 | 0.1856 | 0.5097 | |
| A and B | [500,1000) | 30 | 8 | 1 | 0.1228 | 0.4589 | |
| A and B | | | 8 | 2 | 0.1228 | 0.4589 | |
| A and B | | | 4 | 3 | 0.0376 | 0.3072 | |
| A and B | | | 10 | 4 | 0.1729 | 0.5281 | |
| A and B | [1000,2000) | 62 | 16 | 1 | 0.1553 | 0.385 | |
| A and B | | | 17 | 2 | 0.1685 | 0.4023 | |
| A and B | | | 21 | 3 | 0.2233 | 0.4701 | |
| A and B | | | 8 | 4 | 0.0574 | 0.2385 | ✓ |
| A and B | [2000,3000) | 80 | 13 | 1 | 0.0895 | 0.2618 | |
| A and B | | | 21 | 2 | 0.1704 | 0.3729 | |
| A and B | | | 32 | 3 | 0.292 | 0.5156 | ✓ |
| A and B | | | 14 | 4 | 0.0991 | 0.2762 | |
| A and B | [3000,5000) | 197 | 38 | 1 | 0.1403 | 0.255 | |
| A and B | | | 66 | 2 | 0.2695 | 0.4056 | ✓ |
| A and B | | | 62 | 3 | 0.2506 | 0.3845 | ✓ |
| A and B | | | 31 | 4 | 0.1095 | 0.2159 | ✓ |
| A and B | [5000,10000) | 719 | 193 | 1 | 0.2363 | 0.3024 | |
| A and B | | | 176 | 2 | 0.2138 | 0.2779 | |
| A and B | | | 196 | 3 | 0.2403 | 0.3067 | |
| A and B | | | 154 | 4 | 0.1847 | 0.246 | ✓ |

#### *5.4. Hot-dry season*

| Pair | Distance interval (m) | Total | Count | Type | Lower CI | Upper CI | Sign. diff. |
|---|---|---|---|---|---|---|---|
| A and B | [0,50) | 397 | 43 | 1 | 0.0795 | 0.1431 | ✓ |
| A and B | | | 19 | 2 | 0.0291 | 0.0737 | ✓ |
| A and B | | | 153 | 3 | 0.3373 | 0.4352 | ✓ |
| A and B | | | 182 | 4 | 0.4086 | 0.5089 | ✓ |
| A and B | [50,100) | 427 | 18 | 1 | 0.0252 | 0.0658 | ✓ |
| A and B | | | 33 | 2 | 0.0538 | 0.1068 | ✓ |
| A and B | | | 174 | 3 | 0.3605 | 0.4558 | ✓ |
| A and B | | | 202 | 4 | 0.4249 | 0.5216 | ✓ |
| A and B | [100,200) | 657 | 55 | 1 | 0.0637 | 0.1076 | ✓ |
| A and B | | | 51 | 2 | 0.0583 | 0.1008 | ✓ |
| A and B | | | 271 | 3 | 0.3745 | 0.4512 | ✓ |
| A and B | | | 280 | 4 | 0.388 | 0.465 | ✓ |
| A and B | [200,500) | 1195 | 158 | 1 | 0.1135 | 0.1527 | ✓ |
| A and B | | | 153 | 2 | 0.1096 | 0.1483 | ✓ |
| A and B | | | 354 | 3 | 0.2705 | 0.323 | ✓ |
| A and B | | | 530 | 4 | 0.4151 | 0.4722 | ✓ |
| A and B | [500,1000) | 707 | 96 | 1 | 0.1114 | 0.1633 | ✓ |
| A and B | | | 104 | 2 | 0.1218 | 0.1754 | ✓ |
| A and B | | | 141 | 3 | 0.1706 | 0.2308 | ✓ |
| A and B | | | 366 | 4 | 0.4801 | 0.5551 | ✓ |
| A and B | [1000,2000) | 426 | 77 | 1 | 0.1454 | 0.2206 | ✓ |
| A and B | | | 64 | 2 | 0.1177 | 0.1878 | ✓ |
| A and B | | | 70 | 3 | 0.1304 | 0.203 | ✓ |
| A and B | | | 215 | 4 | 0.4562 | 0.5532 | ✓ |
| A and B | [2000,3000) | 285 | 46 | 1 | 0.1207 | 0.2094 | ✓ |
| A and B | | | 67 | 2 | 0.1871 | 0.2887 | |
| A and B | | | 62 | 3 | 0.171 | 0.27 | |
| A and B | | | 110 | 4 | 0.3291 | 0.4452 | ✓ |
| A and B | [3000,5000) | 271 | 61 | 1 | 0.1768 | 0.2795 | |
| A and B | | | 55 | 2 | 0.1567 | 0.2558 | |
| A and B | | | 82 | 3 | 0.2485 | 0.3611 | |
| A and B | | | 73 | 4 | 0.2175 | 0.3264 | |
| A and B | [5000,10000) | 656 | 180 | 1 | 0.2406 | 0.3103 | |
| A and B | | | 136 | 2 | 0.1769 | 0.2404 | ✓ |
| A and B | | | 140 | 3 | 0.1826 | 0.2468 | ✓ |
| A and B | | | 200 | 4 | 0.2698 | 0.3417 | ✓ |

In Fig. 9 we show the pair distance, coloring each point in time with the color corresponding to the dyadic behaviour type. Also in this case, we observe a decrease of the pair distance with approach behaviour (indicated in blue), as expected, as well as an increase in the pair distance with retreat behaviour (shown in red).

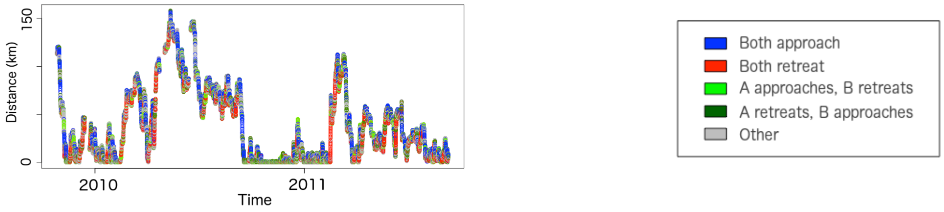


Figure 9: Distance between the pair, colored according to the dyadic behaviour type (legend on the right).

### 6. All pairs of interest

Also in this case, we indicate behaviours of type 3(A,B) and 3(B,A) with numbers 3 and 4.

Table 1: Results - 15-min pairs

| Distance interval | Type 1 (below) | Type 1 (above) | Type 2 (below) | Type 2 (above) | Type 3 (below) | Type 3 (above) | Type 4 (below) | Type 4 (above) |
|---|---|---|---|---|---|---|---|---|
| [0m,50m) | 29 | 18 | 88 | 0 | 0 | 59 | 0 | 65 |
| [50m,100m) | 53 | 0 | 65 | 0 | 0 | 53 | 0 | 41 |
| [100m,200m) | 59 | 0 | 71 | 0 | 0 | 53 | 0 | 41 |
| [200m,500m) | 59 | 0 | 71 | 0 | 0 | 65 | 0 | 59 |
| [500m,1km) | 53 | 0 | 12 | 0 | 0 | 47 | 6 | 35 |
| [1km,2km) | 53 | 0 | 18 | 0 | 12 | 53 | 18 | 24 |
| [2km,3km) | 41 | 0 | 12 | 12 | 6 | 41 | 18 | 29 |
| [3km,5km) | 0 | 18 | 12 | 12 | 18 | 24 | 24 | 24 |
| [5km,10km) | 6 | 24 | 35 | 18 | 29 | 29 | 35 | 18 |

Table 2: Results - 20-min pairs

| Distance interval | Type 1 (below) | Type 1 (above) | Type 2 (below) | Type 2 (above) | Type 3 (below) | Type 3 (above) | Type 4 (below) | Type 4 (above) |
|---|---|---|---|---|---|---|---|---|
| [0m,50m) | 52 | 0 | 83 | 0 | 0 | 61 | 4 | 70 |
| [50m,100m) | 65 | 0 | 65 | 0 | 0 | 74 | 4 | 48 |
| [100m,200m) | 87 | 0 | 78 | 0 | 0 | 83 | 4 | 61 |
| [200m,500m) | 87 | 0 | 78 | 0 | 0 | 87 | 4 | 61 |
| [500m,1km) | 61 | 0 | 39 | 0 | 9 | 39 | 13 | 57 |
| [1km,2km) | 61 | 0 | 43 | 0 | 13 | 52 | 13 | 30 |
| [2km,3km) | 30 | 0 | 17 | 0 | 17 | 26 | 13 | 35 |
| [3km,5km) | 22 | 9 | 13 | 0 | 26 | 17 | 9 | 30 |
| [5km,10km) | 17 | 13 | 4 | 26 | 17 | 17 | 30 | 13 |

Table 3: Results - 30-min pairs

| Distance interval | Type 1 (below) | Type 1 (above) | Type 2 (below) | Type 2 (above) | Type 3 (below) | Type 3 (above) | Type 4 (below) | Type 4 (above) |
|---|---|---|---|---|---|---|---|---|
| [0m,50m) | 15 | 0 | 23 | 0 | 0 | 23 | 0 | 15 |
| [50m,100m) | 23 | 0 | 38 | 0 | 0 | 38 | 0 | 23 |
| [100m,200m) | 46 | 0 | 46 | 0 | 0 | 46 | 0 | 46 |
| [200m,500m) | 85 | 0 | 69 | 0 | 0 | 77 | 0 | 69 |
| [500m,1km) | 77 | 0 | 54 | 0 | 0 | 77 | 8 | 54 |
| [1km,2km) | 46 | 0 | 77 | 0 | 15 | 54 | 0 | 54 |
| [2km,3km) | 15 | 0 | 23 | 0 | 0 | 15 | 15 | 31 |
| [3km,5km) | 0 | 23 | 15 | 0 | 23 | 0 | 0 | 23 |
| [5km,10km) | 0 | 38 | 54 | 0 | 15 | 31 | 15 | 23 |

Table 4: Results - 20-min pairs (male-male)

| Distance interval | Type 1 (below) | Type 1 (above) | Type 2 (below) | Type 2 (above) | Type 3 (below) | Type 3 (above) | Type 4 (below) | Type 4 (above) |
|---|---|---|---|---|---|---|---|---|
| [0m,50m) | 55 | 0 | 100 | 0 | 0 | 73 | 9 | 73 |
| [50m,100m) | 64 | 0 | 73 | 0 | 0 | 73 | 9 | 45 |
| [100m,200m) | 82 | 0 | 91 | 0 | 0 | 82 | 9 | 64 |
| [200m,500m) | 82 | 0 | 82 | 0 | 0 | 91 | 9 | 45 |
| [500m,1km) | 45 | 0 | 18 | 0 | 9 | 55 | 18 | 36 |
| [1km,2km) | 45 | 0 | 27 | 0 | 0 | 64 | 27 | 9 |
| [2km,3km) | 9 | 0 | 0 | 0 | 9 | 18 | 18 | 9 |
| [3km,5km) | 18 | 18 | 0 | 0 | 27 | 9 | 9 | 18 |
| [5km,10km) | 18 | 18 | 9 | 18 | 0 | 36 | 45 | 9 |

Table 5: Results - 20-min pairs (male-female)

| Distance interval | Type 1 (below) | Type 1 (above) | Type 2 (below) | Type 2 (above) | Type 3 (below) | Type 3 (above) | Type 4 (below) | Type 4 (above) |
|---|---|---|---|---|---|---|---|---|
| [0m,50m) | 80 | 0 | 100 | 0 | 0 | 80 | 0 | 100 |
| [50m,100m) | 100 | 0 | 100 | 0 | 0 | 100 | 0 | 80 |
| [100m,200m) | 100 | 0 | 100 | 0 | 0 | 100 | 0 | 80 |
| [200m,500m) | 80 | 0 | 100 | 0 | 0 | 100 | 0 | 60 |
| [500m,1km) | 80 | 0 | 80 | 0 | 0 | 100 | 40 | 0 |
| [1km,2km) | 80 | 0 | 80 | 0 | 0 | 100 | 60 | 0 |
| [2km,3km) | 40 | 0 | 40 | 0 | 0 | 100 | 60 | 0 |
| [3km,5km) | 40 | 0 | 0 | 0 | 0 | 40 | 60 | 40 |
| [5km,10km) | 0 | 0 | 0 | 20 | 20 | 40 | 20 | 0 |

Table 6: Results - 20-min pairs (female-female)

| Distance interval | Type 1 (below) | Type 1 (above) | Type 2 (below) | Type 2 (above) | Type 3 (below) | Type 3 (above) | Type 4 (below) | Type 4 (above) |
|---|---|---|---|---|---|---|---|---|
| [0m,50m) | 29 | 0 | 43 | 0 | 0 | 29 | 0 | 43 |
| [50m,100m) | 43 | 0 | 29 | 0 | 0 | 57 | 0 | 29 |
| [100m,200m) | 86 | 0 | 43 | 0 | 0 | 71 | 0 | 43 |
| [200m,500m) | 100 | 0 | 57 | 0 | 0 | 86 | 0 | 71 |
| [500m,1km) | 71 | 0 | 43 | 0 | 0 | 29 | 0 | 71 |
| [1km,2km) | 71 | 0 | 43 | 0 | 0 | 57 | 0 | 29 |
| [2km,3km) | 57 | 0 | 29 | 0 | 14 | 43 | 0 | 43 |
| [3km,5km) | 14 | 0 | 43 | 0 | 14 | 14 | 0 | 43 |
| [5km,10km) | 29 | 14 | 0 | 43 | 29 | 0 | 29 | 0 |

### *6.1. 15-min pairs*

| Pair | Distance interval (m) | Total | Count | Type | Lower CI | Upper CI | Sign. diff. |
|---|---|---|---|---|---|---|---|
| 1 and 2 | [0,50) | 77 | 19 | 1 | 0.1556 | 0.3582 | |
| 1 and 2 | | 77 | 7 | 2 | 0.0373 | 0.1784 | ✓ |
| 1 and 2 | | 77 | 20 | 3 | 0.1664 | 0.3723 | |
| 1 and 2 | | 77 | 31 | 4 | 0.2923 | 0.5206 | ✓ |
| 1 and 2 | [50,100) | 25 | 3 | 1 | 0.0255 | 0.3122 | |
| 1 and 2 | | 25 | 2 | 2 | 0.0098 | 0.2603 | |
| 1 and 2 | | 25 | 9 | 3 | 0.1797 | 0.5748 | |
| 1 and 2 | | 25 | 11 | 4 | 0.244 | 0.6507 | |
| 1 and 2 | [100,200) | 60 | 12 | 1 | 0.1078 | 0.3233 | |
| 1 and 2 | | 60 | 11 | 2 | 0.0952 | 0.3044 | |
| 1 and 2 | | 60 | 21 | 3 | 0.2313 | 0.484 | |
| 1 and 2 | | 60 | 16 | 4 | 0.1607 | 0.3966 | |
| 1 and 2 | [200,500) | 87 | 13 | 1 | 0.082 | 0.242 | ✓ |
| 1 and 2 | | 87 | 13 | 2 | 0.082 | 0.242 | ✓ |
| 1 and 2 | | 87 | 47 | 3 | 0.43 | 0.6477 | ✓ |
| 1 and 2 | | 87 | 14 | 4 | 0.0909 | 0.2552 | |
| 1 and 2 | [500,1000) | 125 | 24 | 1 | 0.1271 | 0.2721 | |
| 1 and 2 | | 125 | 33 | 2 | 0.1892 | 0.3503 | |
| 1 and 2 | | 125 | 23 | 3 | 0.1204 | 0.2632 | |
| 1 and 2 | | 125 | 45 | 4 | 0.2761 | 0.4507 | ✓ |
| 1 and 2 | [1000,2000) | 251 | 50 | 1 | 0.1516 | 0.254 | |
| 1 and 2 | | 251 | 74 | 2 | 0.2391 | 0.3554 | |
| 1 and 2 | | 251 | 55 | 3 | 0.1695 | 0.2755 | |
| 1 and 2 | | 251 | 72 | 4 | 0.2317 | 0.3471 | |
| 1 and 2 | [2000,3000) | 410 | 77 | 1 | 0.1512 | 0.229 | ✓ |
| 1 and 2 | | 410 | 98 | 2 | 0.1985 | 0.2833 | |
| 1 and 2 | | 410 | 107 | 3 | 0.2191 | 0.3063 | |
| 1 and 2 | | 410 | 128 | 4 | 0.2676 | 0.3595 | ✓ |
| 1 and 2 | [3000,5000) | 1095 | 246 | 1 | 0.2002 | 0.2506 | |
| 1 and 2 | | 1095 | 249 | 2 | 0.2029 | 0.2534 | |
| 1 and 2 | | 1095 | 266 | 3 | 0.2178 | 0.2695 | |
| 1 and 2 | | 1095 | 334 | 4 | 0.2779 | 0.3332 | ✓ |
| 1 and 2 | [5000,10000) | 3838 | 965 | 1 | 0.2378 | 0.2655 | |
| 1 and 2 | | 3838 | 970 | 2 | 0.239 | 0.2668 | |
| 1 and 2 | | 3838 | 939 | 3 | 0.2311 | 0.2586 | |
| 1 and 2 | | 3838 | 964 | 4 | 0.2375 | 0.2652 | |

| Pair | Distance interval (m) | Total | Count | Type | Lower CI | Upper CI | Sign. diff. |
|---|---|---|---|---|---|---|---|
| 1 and 3 | [0,50) | 393 | 102 | 1 | 0.2169 | 0.3059 | |
| 1 and 3 | | 393 | 28 | 2 | 0.0479 | 0.1013 | ✓ |
| 1 and 3 | | 393 | 126 | 3 | 0.2747 | 0.3692 | ✓ |
| 1 and 3 | | 393 | 137 | 4 | 0.3015 | 0.398 | ✓ |
| 1 and 3 | [50,100) | 132 | 19 | 1 | 0.0889 | 0.2156 | ✓ |
| 1 and 3 | | 132 | 10 | 2 | 0.0369 | 0.1349 | ✓ |
| 1 and 3 | | 132 | 45 | 3 | 0.2607 | 0.4284 | ✓ |
| 1 and 3 | | 132 | 58 | 4 | 0.3532 | 0.5284 | ✓ |
| 1 and 3 | [100,200) | 183 | 34 | 1 | 0.1322 | 0.2498 | ✓ |
| 1 and 3 | | 183 | 33 | 2 | 0.1275 | 0.2438 | ✓ |
| 1 and 3 | | 183 | 60 | 3 | 0.2604 | 0.401 | ✓ |
| 1 and 3 | | 183 | 56 | 4 | 0.2402 | 0.3783 | |
| 1 and 3 | [200,500) | 343 | 56 | 1 | 0.1257 | 0.2067 | ✓ |
| 1 and 3 | | 343 | 60 | 2 | 0.1362 | 0.2194 | ✓ |
| 1 and 3 | | 343 | 145 | 3 | 0.3699 | 0.477 | ✓ |
| 1 and 3 | | 343 | 82 | 4 | 0.1949 | 0.2878 | |
| 1 and 3 | [500,1000) | 576 | 99 | 1 | 0.1419 | 0.2052 | ✓ |
| 1 and 3 | | 576 | 127 | 2 | 0.1873 | 0.2566 | |
| 1 and 3 | | 576 | 203 | 3 | 0.3134 | 0.393 | ✓ |
| 1 and 3 | | 576 | 147 | 4 | 0.2201 | 0.2929 | |
| 1 and 3 | [1000,2000) | 1255 | 256 | 1 | 0.182 | 0.2274 | ✓ |
| 1 and 3 | | 1255 | 273 | 2 | 0.195 | 0.2414 | ✓ |
| 1 and 3 | | 1255 | 365 | 3 | 0.2658 | 0.3168 | ✓ |
| 1 and 3 | | 1255 | 361 | 4 | 0.2627 | 0.3136 | ✓ |
| 1 and 3 | [2000,3000) | 1579 | 375 | 1 | 0.2167 | 0.2593 | |
| 1 and 3 | | 1579 | 365 | 2 | 0.2106 | 0.2528 | |
| 1 and 3 | | 1579 | 430 | 3 | 0.2505 | 0.295 | ✓ |
| 1 and 3 | | 1579 | 409 | 4 | 0.2376 | 0.2814 | |
| 1 and 3 | [3000,5000) | 2833 | 719 | 1 | 0.2379 | 0.2702 | |
| 1 and 3 | | 2833 | 701 | 2 | 0.2316 | 0.2638 | |
| 1 and 3 | | 2833 | 769 | 3 | 0.2551 | 0.2882 | ✓ |
| 1 and 3 | | 2833 | 644 | 4 | 0.212 | 0.2432 | ✓ |
| 1 and 3 | [5000,10000) | 7976 | 1931 | 1 | 0.2327 | 0.2517 | |
| 1 and 3 | | 7976 | 1977 | 2 | 0.2384 | 0.2575 | |
| 1 and 3 | | 7976 | 2106 | 3 | 0.2544 | 0.2739 | ✓ |
| 1 and 3 | | 7976 | 1962 | 4 | 0.2366 | 0.2556 | |

| Pair | Distance interval (m) | Total | Count | Type | Lower CI | Upper CI | Sign. diff. |
|---|---|---|---|---|---|---|---|
| 1 and 4 | [0,50) | 164 | 58 | 1 | 0.2807 | 0.432 | ✓ |
| 1 and 4 | | 164 | 11 | 2 | 0.034 | 0.1168 | ✓ |
| 1 and 4 | | 164 | 44 | 3 | 0.2022 | 0.343 | |
| 1 and 4 | | 164 | 51 | 4 | 0.2411 | 0.3878 | |
| 1 and 4 | [50,100) | 61 | 11 | 1 | 0.0936 | 0.2998 | |
| 1 and 4 | | 61 | 8 | 2 | 0.0584 | 0.2422 | ✓ |
| 1 and 4 | | 61 | 26 | 3 | 0.3004 | 0.5594 | ✓ |
| 1 and 4 | | 61 | 16 | 4 | 0.158 | 0.3907 | |
| 1 and 4 | [100,200) | 116 | 17 | 1 | 0.0878 | 0.2242 | ✓ |
| 1 and 4 | | 116 | 22 | 2 | 0.1228 | 0.2729 | |
| 1 and 4 | | 116 | 35 | 3 | 0.22 | 0.3939 | |
| 1 and 4 | | 116 | 42 | 4 | 0.2749 | 0.4565 | ✓ |
| 1 and 4 | [200,500) | 147 | 31 | 1 | 0.148 | 0.2858 | |
| 1 and 4 | | 147 | 31 | 2 | 0.148 | 0.2858 | |
| 1 and 4 | | 147 | 35 | 3 | 0.1718 | 0.3153 | |
| 1 and 4 | | 147 | 50 | 4 | 0.2641 | 0.4228 | ✓ |
| 1 and 4 | [500,1000) | 136 | 26 | 1 | 0.1288 | 0.2674 | |
| 1 and 4 | | 136 | 26 | 2 | 0.1288 | 0.2674 | |
| 1 and 4 | | 136 | 44 | 3 | 0.2459 | 0.409 | |
| 1 and 4 | | 136 | 40 | 4 | 0.2191 | 0.3783 | |
| 1 and 4 | [1000,2000) | 283 | 79 | 1 | 0.2277 | 0.3353 | |
| 1 and 4 | | 283 | 67 | 2 | 0.1884 | 0.2907 | |
| 1 and 4 | | 283 | 55 | 3 | 0.1499 | 0.2453 | ✓ |
| 1 and 4 | | 283 | 82 | 4 | 0.2376 | 0.3464 | |
| 1 and 4 | [2000,3000) | 572 | 124 | 1 | 0.1837 | 0.2528 | |
| 1 and 4 | | 572 | 149 | 2 | 0.225 | 0.2985 | |
| 1 and 4 | | 572 | 179 | 3 | 0.2751 | 0.3527 | ✓ |
| 1 and 4 | | 572 | 120 | 4 | 0.1771 | 0.2455 | ✓ |
| 1 and 4 | [3000,5000) | 1585 | 367 | 1 | 0.211 | 0.2531 | |
| 1 and 4 | | 1585 | 428 | 2 | 0.2483 | 0.2926 | |
| 1 and 4 | | 1585 | 470 | 3 | 0.2741 | 0.3197 | ✓ |
| 1 and 4 | | 1585 | 320 | 4 | 0.1824 | 0.2225 | ✓ |
| 1 and 4 | [5000,10000) | 4880 | 1164 | 1 | 0.2266 | 0.2507 | |
| 1 and 4 | | 4880 | 1412 | 2 | 0.2766 | 0.3023 | ✓ |
| 1 and 4 | | 4880 | 1194 | 3 | 0.2327 | 0.257 | |
| 1 and 4 | | 4880 | 1110 | 4 | 0.2158 | 0.2395 | ✓ |

| Pair | Distance interval (m) | Total | Count | Type | Lower CI | Upper CI | Sign. diff. |
|---|---|---|---|---|---|---|---|
| 1 and 5 | [0,50) | 1047 | 165 | 1 | 0.136 | 0.1811 | ✓ |
| 1 and 5 | | 1047 | 57 | 2 | 0.0415 | 0.07 | ✓ |
| 1 and 5 | | 1047 | 463 | 3 | 0.4119 | 0.4729 | ✓ |
| 1 and 5 | | 1047 | 362 | 4 | 0.3169 | 0.3754 | ✓ |
| 1 and 5 | [50,100) | 410 | 40 | 1 | 0.0706 | 0.1305 | ✓ |
| 1 and 5 | | 410 | 27 | 2 | 0.0438 | 0.0944 | ✓ |
| 1 and 5 | | 410 | 211 | 3 | 0.4651 | 0.564 | ✓ |
| 1 and 5 | | 410 | 132 | 4 | 0.2769 | 0.3696 | ✓ |
| 1 and 5 | [100,200) | 517 | 66 | 1 | 0.1001 | 0.1595 | ✓ |
| 1 and 5 | | 517 | 37 | 2 | 0.0509 | 0.0973 | ✓ |
| 1 and 5 | | 517 | 265 | 3 | 0.4686 | 0.5564 | ✓ |
| 1 and 5 | | 517 | 149 | 4 | 0.2495 | 0.3293 | |
| 1 and 5 | [200,500) | 862 | 137 | 1 | 0.1351 | 0.1851 | ✓ |
| 1 and 5 | | 862 | 118 | 2 | 0.1146 | 0.1617 | ✓ |
| 1 and 5 | | 862 | 365 | 3 | 0.3902 | 0.4572 | ✓ |
| 1 and 5 | | 862 | 242 | 4 | 0.251 | 0.312 | ✓ |
| 1 and 5 | [500,1000) | 819 | 137 | 1 | 0.1423 | 0.1946 | ✓ |
| 1 and 5 | | 819 | 148 | 2 | 0.1549 | 0.2088 | ✓ |
| 1 and 5 | | 819 | 289 | 3 | 0.3201 | 0.3867 | ✓ |
| 1 and 5 | | 819 | 245 | 4 | 0.2679 | 0.3318 | ✓ |
| 1 and 5 | [1000,2000) | 1340 | 292 | 1 | 0.1961 | 0.241 | ✓ |
| 1 and 5 | | 1340 | 268 | 2 | 0.1789 | 0.2224 | ✓ |
| 1 and 5 | | 1340 | 382 | 3 | 0.261 | 0.3101 | ✓ |
| 1 and 5 | | 1340 | 398 | 4 | 0.2726 | 0.3223 | ✓ |
| 1 and 5 | [2000,3000) | 1282 | 246 | 1 | 0.1707 | 0.2145 | ✓ |
| 1 and 5 | | 1282 | 297 | 2 | 0.2088 | 0.2558 | |
| 1 and 5 | | 1282 | 363 | 3 | 0.2586 | 0.3087 | ✓ |
| 1 and 5 | | 1282 | 376 | 4 | 0.2685 | 0.3191 | ✓ |
| 1 and 5 | [3000,5000) | 2394 | 587 | 1 | 0.2281 | 0.2629 | |
| 1 and 5 | | 2394 | 599 | 2 | 0.233 | 0.2681 | |
| 1 and 5 | | 2394 | 623 | 3 | 0.2428 | 0.2783 | |
| 1 and 5 | | 2394 | 585 | 4 | 0.2273 | 0.2621 | |
| 1 and 5 | [5000,10000) | 5457 | 1396 | 1 | 0.2443 | 0.2676 | |
| 1 and 5 | | 5457 | 1174 | 2 | 0.2043 | 0.2263 | ✓ |
| 1 and 5 | | 5457 | 1496 | 3 | 0.2623 | 0.2862 | ✓ |
| 1 and 5 | | 5457 | 1391 | 4 | 0.2434 | 0.2667 | |

| Pair | Distance interval (m) | Total | Count | Type | Lower CI | Upper CI | Sign. diff. |
|---|---|---|---|---|---|---|---|
| 2 and 3 | [0,50) | 780 | 113 | 1 | 0.1209 | 0.1716 | ✓ |
| 2 and 3 | | 780 | 57 | 2 | 0.0558 | 0.0936 | ✓ |
| 2 and 3 | | 780 | 252 | 3 | 0.2903 | 0.3572 | ✓ |
| 2 and 3 | | 780 | 358 | 4 | 0.4236 | 0.4947 | ✓ |
| 2 and 3 | [50,100) | 532 | 61 | 1 | 0.0889 | 0.1448 | ✓ |
| 2 and 3 | | 532 | 41 | 2 | 0.0559 | 0.1031 | ✓ |
| 2 and 3 | | 532 | 193 | 3 | 0.3218 | 0.4053 | ✓ |
| 2 and 3 | | 532 | 237 | 4 | 0.4027 | 0.4889 | ✓ |
| 2 and 3 | [100,200) | 544 | 57 | 1 | 0.0803 | 0.1336 | ✓ |
| 2 and 3 | | 544 | 49 | 2 | 0.0674 | 0.1173 | ✓ |
| 2 and 3 | | 544 | 212 | 3 | 0.3485 | 0.4321 | ✓ |
| 2 and 3 | | 544 | 226 | 4 | 0.3737 | 0.4581 | ✓ |
| 2 and 3 | [200,500) | 516 | 95 | 1 | 0.1516 | 0.2203 | ✓ |
| 2 and 3 | | 516 | 85 | 2 | 0.1338 | 0.1996 | ✓ |
| 2 and 3 | | 516 | 207 | 3 | 0.3586 | 0.4449 | ✓ |
| 2 and 3 | | 516 | 129 | 4 | 0.2132 | 0.2897 | |
| 2 and 3 | [500,1000) | 384 | 82 | 1 | 0.1736 | 0.258 | |
| 2 and 3 | | 384 | 102 | 2 | 0.2221 | 0.3128 | |
| 2 and 3 | | 384 | 106 | 3 | 0.2319 | 0.3237 | |
| 2 and 3 | | 384 | 94 | 4 | 0.2026 | 0.291 | |
| 2 and 3 | [1000,2000) | 728 | 158 | 1 | 0.1876 | 0.2488 | ✓ |
| 2 and 3 | | 728 | 190 | 2 | 0.2294 | 0.2945 | |
| 2 and 3 | | 728 | 199 | 3 | 0.2413 | 0.3073 | |
| 2 and 3 | | 728 | 181 | 4 | 0.2176 | 0.2817 | |
| 2 and 3 | [2000,3000) | 642 | 148 | 1 | 0.1985 | 0.2651 | |
| 2 and 3 | | 642 | 173 | 2 | 0.2355 | 0.3056 | |
| 2 and 3 | | 642 | 162 | 3 | 0.2192 | 0.2878 | |
| 2 and 3 | | 642 | 159 | 4 | 0.2147 | 0.2829 | |
| 2 and 3 | [3000,5000) | 1450 | 371 | 1 | 0.2336 | 0.2791 | |
| 2 and 3 | | 1450 | 395 | 2 | 0.2496 | 0.2961 | |
| 2 and 3 | | 1450 | 285 | 3 | 0.1764 | 0.218 | ✓ |
| 2 and 3 | | 1450 | 399 | 4 | 0.2523 | 0.2989 | ✓ |
| 2 and 3 | [5000,10000) | 5017 | 1284 | 1 | 0.2439 | 0.2682 | |
| 2 and 3 | | 5017 | 1257 | 2 | 0.2386 | 0.2628 | |
| 2 and 3 | | 5017 | 1178 | 3 | 0.2231 | 0.2468 | ✓ |
| 2 and 3 | | 5017 | 1298 | 4 | 0.2466 | 0.2711 | |

| Pair | Distance interval (m) | Total | Count | Type | Lower CI | Upper CI | Sign. diff. |
|---|---|---|---|---|---|---|---|
| 2 and 4 | [0,50) | 348 | 32 | 1 | 0.0637 | 0.1273 | ✓ |
| 2 and 4 | | 348 | 33 | 2 | 0.0662 | 0.1306 | ✓ |
| 2 and 4 | | 348 | 112 | 3 | 0.273 | 0.3737 | ✓ |
| 2 and 4 | | 348 | 171 | 4 | 0.4377 | 0.5452 | ✓ |
| 2 and 4 | [50,100) | 252 | 19 | 1 | 0.046 | 0.1152 | ✓ |
| 2 and 4 | | 252 | 21 | 2 | 0.0523 | 0.1246 | ✓ |
| 2 and 4 | | 252 | 66 | 3 | 0.2087 | 0.3208 | |
| 2 and 4 | | 252 | 146 | 4 | 0.5158 | 0.641 | ✓ |
| 2 and 4 | [100,200) | 282 | 40 | 1 | 0.1033 | 0.1881 | ✓ |
| 2 and 4 | | 282 | 27 | 2 | 0.064 | 0.1362 | ✓ |
| 2 and 4 | | 282 | 81 | 3 | 0.2351 | 0.3439 | |
| 2 and 4 | | 282 | 134 | 4 | 0.4156 | 0.5352 | ✓ |
| 2 and 4 | [200,500) | 392 | 66 | 1 | 0.1327 | 0.2092 | ✓ |
| 2 and 4 | | 392 | 59 | 2 | 0.1166 | 0.1898 | ✓ |
| 2 and 4 | | 392 | 95 | 3 | 0.2007 | 0.2879 | |
| 2 and 4 | | 392 | 172 | 4 | 0.389 | 0.4895 | ✓ |
| 2 and 4 | [500,1000) | 288 | 65 | 1 | 0.1787 | 0.2784 | |
| 2 and 4 | | 288 | 61 | 2 | 0.1661 | 0.2636 | |
| 2 and 4 | | 288 | 78 | 3 | 0.2204 | 0.3261 | |
| 2 and 4 | | 288 | 84 | 4 | 0.2398 | 0.3479 | |
| 2 and 4 | [1000,2000) | 363 | 84 | 1 | 0.189 | 0.2783 | |
| 2 and 4 | | 363 | 98 | 2 | 0.225 | 0.3188 | |
| 2 and 4 | | 363 | 125 | 3 | 0.2955 | 0.3957 | ✓ |
| 2 and 4 | | 363 | 56 | 4 | 0.1187 | 0.1956 | ✓ |
| 2 and 4 | [2000,3000) | 427 | 100 | 1 | 0.1948 | 0.2773 | |
| 2 and 4 | | 427 | 102 | 2 | 0.1992 | 0.2822 | |
| 2 and 4 | | 427 | 127 | 3 | 0.2544 | 0.3433 | ✓ |
| 2 and 4 | | 427 | 98 | 4 | 0.1904 | 0.2724 | |
| 2 and 4 | [3000,5000) | 1125 | 258 | 1 | 0.2051 | 0.255 | |
| 2 and 4 | | 1125 | 298 | 2 | 0.2393 | 0.2917 | |
| 2 and 4 | | 1125 | 256 | 3 | 0.2034 | 0.2532 | |
| 2 and 4 | | 1125 | 313 | 4 | 0.2522 | 0.3054 | ✓ |
| 2 and 4 | [5000,10000) | 4291 | 1080 | 1 | 0.2388 | 0.265 | |
| 2 and 4 | | 4291 | 1080 | 2 | 0.2388 | 0.265 | |
| 2 and 4 | | 4291 | 1013 | 3 | 0.2234 | 0.2491 | ✓ |
| 2 and 4 | | 4291 | 1118 | 4 | 0.2475 | 0.274 | |

| Pair | Distance interval (m) | Total | Count | Type | Lower CI | Upper CI | Sign. diff. |
|---|---|---|---|---|---|---|---|
| 3 and 4 | [0,50) | 155 | 51 | 1 | 0.2558 | 0.409 | ✓ |
| 3 and 4 | | 155 | 6 | 2 | 0.0143 | 0.0823 | ✓ |
| 3 and 4 | | 155 | 48 | 3 | 0.238 | 0.3888 | |
| 3 and 4 | | 155 | 50 | 4 | 0.2498 | 0.4023 | |
| 3 and 4 | [50,100) | 46 | 5 | 1 | 0.0362 | 0.2357 | ✓ |
| 3 and 4 | | 46 | 6 | 2 | 0.0494 | 0.2626 | |
| 3 and 4 | | 46 | 19 | 3 | 0.27 | 0.5677 | ✓ |
| 3 and 4 | | 46 | 16 | 4 | 0.2135 | 0.5025 | |
| 3 and 4 | [100,200) | 85 | 13 | 1 | 0.084 | 0.2473 | ✓ |
| 3 and 4 | | 85 | 7 | 2 | 0.0338 | 0.1623 | ✓ |
| 3 and 4 | | 85 | 30 | 3 | 0.2523 | 0.4641 | ✓ |
| 3 and 4 | | 85 | 35 | 4 | 0.3061 | 0.5238 | ✓ |
| 3 and 4 | [200,500) | 128 | 24 | 1 | 0.124 | 0.266 | |
| 3 and 4 | | 128 | 24 | 2 | 0.124 | 0.266 | |
| 3 and 4 | | 128 | 32 | 3 | 0.1777 | 0.3342 | |
| 3 and 4 | | 128 | 48 | 4 | 0.291 | 0.4649 | ✓ |
| 3 and 4 | [500,1000) | 248 | 44 | 1 | 0.132 | 0.2308 | ✓ |
| 3 and 4 | | 248 | 64 | 2 | 0.2048 | 0.3172 | |
| 3 and 4 | | 248 | 49 | 3 | 0.1499 | 0.2527 | |
| 3 and 4 | | 248 | 91 | 4 | 0.3068 | 0.4302 | ✓ |
| 3 and 4 | [1000,2000) | 600 | 122 | 1 | 0.1718 | 0.2378 | ✓ |
| 3 and 4 | | 600 | 144 | 2 | 0.2063 | 0.2762 | |
| 3 and 4 | | 600 | 149 | 3 | 0.2142 | 0.2849 | |
| 3 and 4 | | 600 | 185 | 4 | 0.2716 | 0.347 | ✓ |
| 3 and 4 | [2000,3000) | 710 | 166 | 1 | 0.2031 | 0.2667 | |
| 3 and 4 | | 710 | 196 | 2 | 0.2435 | 0.3105 | |
| 3 and 4 | | 710 | 144 | 3 | 0.1738 | 0.2343 | ✓ |
| 3 and 4 | | 710 | 204 | 4 | 0.2543 | 0.3221 | ✓ |
| 3 and 4 | [3000,5000) | 1741 | 438 | 1 | 0.2313 | 0.2727 | |
| 3 and 4 | | 1741 | 436 | 2 | 0.2302 | 0.2715 | |
| 3 and 4 | | 1741 | 434 | 3 | 0.2291 | 0.2703 | |
| 3 and 4 | | 1741 | 433 | 4 | 0.2286 | 0.2697 | |
| 3 and 4 | [5000,10000) | 6053 | 1589 | 1 | 0.2515 | 0.2738 | ✓ |
| 3 and 4 | | 6053 | 1515 | 2 | 0.2394 | 0.2614 | |
| 3 and 4 | | 6053 | 1549 | 3 | 0.245 | 0.2671 | |
| 3 and 4 | | 6053 | 1400 | 4 | 0.2207 | 0.2421 | ✓ |

| Pair | Distance interval (m) | Total | Count | Type | Lower CI | Upper CI | Sign. diff. |
|---|---|---|---|---|---|---|---|
| 3 and 5 | [0,50) | 194 | 73 | 1 | 0.3079 | 0.4485 | ✓ |
| 3 and 5 | | 194 | 17 | 2 | 0.0519 | 0.1366 | ✓ |
| 3 and 5 | | 194 | 41 | 3 | 0.1561 | 0.2756 | |
| 3 and 5 | | 194 | 63 | 4 | 0.2594 | 0.3955 | ✓ |
| 3 and 5 | [50,100) | 47 | 15 | 1 | 0.1909 | 0.4712 | |
| 3 and 5 | | 47 | 3 | 2 | 0.0134 | 0.1754 | ✓ |
| 3 and 5 | | 47 | 12 | 3 | 0.1394 | 0.4035 | |
| 3 and 5 | | 47 | 17 | 4 | 0.2267 | 0.5148 | |
| 3 and 5 | [100,200) | 75 | 19 | 1 | 0.1599 | 0.367 | |
| 3 and 5 | | 75 | 11 | 2 | 0.0756 | 0.2473 | ✓ |
| 3 and 5 | | 75 | 17 | 3 | 0.1379 | 0.3379 | |
| 3 and 5 | | 75 | 28 | 4 | 0.2643 | 0.4927 | ✓ |
| 3 and 5 | [200,500) | 119 | 14 | 1 | 0.0658 | 0.1895 | ✓ |
| 3 and 5 | | 119 | 19 | 2 | 0.099 | 0.2381 | ✓ |
| 3 and 5 | | 119 | 45 | 3 | 0.2909 | 0.4716 | ✓ |
| 3 and 5 | | 119 | 41 | 4 | 0.2598 | 0.4372 | ✓ |
| 3 and 5 | [500,1000) | 258 | 49 | 1 | 0.1439 | 0.2432 | ✓ |
| 3 and 5 | | 258 | 54 | 2 | 0.1613 | 0.2641 | |
| 3 and 5 | | 258 | 75 | 3 | 0.236 | 0.3502 | |
| 3 and 5 | | 258 | 80 | 4 | 0.2542 | 0.3704 | ✓ |
| 3 and 5 | [1000,2000) | 598 | 116 | 1 | 0.163 | 0.228 | ✓ |
| 3 and 5 | | 598 | 134 | 2 | 0.1913 | 0.2597 | |
| 3 and 5 | | 598 | 125 | 3 | 0.1771 | 0.2439 | ✓ |
| 3 and 5 | | 598 | 223 | 4 | 0.334 | 0.4131 | ✓ |
| 3 and 5 | [2000,3000) | 811 | 169 | 1 | 0.1809 | 0.238 | ✓ |
| 3 and 5 | | 811 | 213 | 2 | 0.2326 | 0.2944 | |
| 3 and 5 | | 811 | 223 | 3 | 0.2445 | 0.3071 | |
| 3 and 5 | | 811 | 206 | 4 | 0.2244 | 0.2854 | |
| 3 and 5 | [3000,5000) | 1921 | 492 | 1 | 0.2367 | 0.2763 | |
| 3 and 5 | | 1921 | 516 | 2 | 0.2489 | 0.289 | |
| 3 and 5 | | 1921 | 412 | 3 | 0.1963 | 0.2335 | ✓ |
| 3 and 5 | | 1921 | 501 | 4 | 0.2413 | 0.2811 | |
| 3 and 5 | [5000,10000) | 4552 | 1161 | 1 | 0.2424 | 0.268 | |
| 3 and 5 | | 4552 | 1241 | 2 | 0.2597 | 0.2858 | ✓ |
| 3 and 5 | | 4552 | 1075 | 3 | 0.2239 | 0.2488 | ✓ |
| 3 and 5 | | 4552 | 1075 | 4 | 0.2239 | 0.2488 | ✓ |

| Pair | Distance interval (m) | Total | Count | Type | Lower CI | Upper CI | Sign. diff. |
|---|---|---|---|---|---|---|---|
| 6 and 7 | [0,50) | 712 | 93 | 1 | 0.1067 | 0.1576 | ✓ |
| 6 and 7 | | 712 | 38 | 2 | 0.038 | 0.0725 | ✓ |
| 6 and 7 | | 712 | 301 | 3 | 0.3862 | 0.46 | ✓ |
| 6 and 7 | | 712 | 280 | 4 | 0.3572 | 0.4302 | ✓ |
| 6 and 7 | [50,100) | 488 | 59 | 1 | 0.0933 | 0.1532 | ✓ |
| 6 and 7 | | 488 | 34 | 2 | 0.0487 | 0.096 | ✓ |
| 6 and 7 | | 488 | 165 | 3 | 0.2962 | 0.382 | ✓ |
| 6 and 7 | | 488 | 230 | 4 | 0.4263 | 0.5167 | ✓ |
| 6 and 7 | [100,200) | 489 | 67 | 1 | 0.1078 | 0.1707 | ✓ |
| 6 and 7 | | 489 | 67 | 2 | 0.1078 | 0.1707 | ✓ |
| 6 and 7 | | 489 | 154 | 3 | 0.274 | 0.3581 | ✓ |
| 6 and 7 | | 489 | 201 | 4 | 0.3671 | 0.4561 | ✓ |
| 6 and 7 | [200,500) | 567 | 102 | 1 | 0.1491 | 0.214 | ✓ |
| 6 and 7 | | 567 | 81 | 2 | 0.1151 | 0.1744 | ✓ |
| 6 and 7 | | 567 | 163 | 3 | 0.2505 | 0.3267 | ✓ |
| 6 and 7 | | 567 | 221 | 4 | 0.3494 | 0.4313 | ✓ |
| 6 and 7 | [500,1000) | 479 | 103 | 1 | 0.1791 | 0.2546 | |
| 6 and 7 | | 479 | 120 | 2 | 0.2123 | 0.2918 | |
| 6 and 7 | | 479 | 161 | 3 | 0.2939 | 0.3804 | ✓ |
| 6 and 7 | | 479 | 95 | 4 | 0.1635 | 0.2369 | ✓ |
| 6 and 7 | [1000,2000) | 1374 | 314 | 1 | 0.2066 | 0.2517 | |
| 6 and 7 | | 1374 | 312 | 2 | 0.2052 | 0.2502 | |
| 6 and 7 | | 1374 | 391 | 3 | 0.2608 | 0.3092 | ✓ |
| 6 and 7 | | 1374 | 357 | 4 | 0.2368 | 0.2839 | |
| 6 and 7 | [2000,3000) | 1726 | 390 | 1 | 0.2064 | 0.2464 | ✓ |
| 6 and 7 | | 1726 | 410 | 2 | 0.2176 | 0.2583 | |
| 6 and 7 | | 1726 | 434 | 3 | 0.2311 | 0.2726 | |
| 6 and 7 | | 1726 | 492 | 4 | 0.2638 | 0.307 | ✓ |
| 6 and 7 | [3000,5000) | 3092 | 833 | 1 | 0.2538 | 0.2854 | ✓ |
| 6 and 7 | | 3092 | 751 | 2 | 0.2279 | 0.2584 | |
| 6 and 7 | | 3092 | 746 | 3 | 0.2263 | 0.2568 | |
| 6 and 7 | | 3092 | 762 | 4 | 0.2313 | 0.262 | |
| 6 and 7 | [5000,10000) | 6021 | 1558 | 1 | 0.2477 | 0.27 | |
| 6 and 7 | | 6021 | 1405 | 2 | 0.2227 | 0.2442 | ✓ |
| 6 and 7 | | 6021 | 1444 | 3 | 0.2291 | 0.2508 | |
| 6 and 7 | | 6021 | 1614 | 4 | 0.2569 | 0.2794 | ✓ |

| Pair | Distance interval (m) | Total | Count | Type | Lower CI | Upper CI | Sign. diff. |
|---|---|---|---|---|---|---|---|
| 6 and 8 | [0,50) | 32 | 8 | 1 | 0.1146 | 0.434 | |
| 6 and 8 | | 32 | 5 | 2 | 0.0528 | 0.3279 | |
| 6 and 8 | | 32 | 14 | 3 | 0.2636 | 0.6234 | ✓ |
| 6 and 8 | | 32 | 5 | 4 | 0.0528 | 0.3279 | |
| 6 and 8 | [50,100) | 26 | 9 | 1 | 0.1721 | 0.5567 | |
| 6 and 8 | | 26 | 3 | 2 | 0.0245 | 0.3015 | |
| 6 and 8 | | 26 | 11 | 3 | 0.2335 | 0.6308 | |
| 6 and 8 | | 26 | 3 | 4 | 0.0245 | 0.3015 | |
| 6 and 8 | [100,200) | 35 | 9 | 1 | 0.1249 | 0.4326 | |
| 6 and 8 | | 35 | 8 | 2 | 0.1042 | 0.4014 | |
| 6 and 8 | | 35 | 6 | 3 | 0.0656 | 0.3365 | |
| 6 and 8 | | 35 | 12 | 4 | 0.1913 | 0.5221 | |
| 6 and 8 | [200,500) | 98 | 26 | 1 | 0.1812 | 0.3641 | |
| 6 and 8 | | 98 | 19 | 2 | 0.121 | 0.2861 | |
| 6 and 8 | | 98 | 27 | 3 | 0.1901 | 0.375 | |
| 6 and 8 | | 98 | 26 | 4 | 0.1812 | 0.3641 | |
| 6 and 8 | [500,1000) | 215 | 46 | 1 | 0.1611 | 0.2749 | |
| 6 and 8 | | 215 | 48 | 2 | 0.1694 | 0.2849 | |
| 6 and 8 | | 215 | 66 | 3 | 0.246 | 0.3733 | |
| 6 and 8 | | 215 | 55 | 4 | 0.1989 | 0.3196 | |
| 6 and 8 | [1000,2000) | 460 | 96 | 1 | 0.1724 | 0.2487 | ✓ |
| 6 and 8 | | 460 | 131 | 2 | 0.244 | 0.3284 | |
| 6 and 8 | | 460 | 138 | 3 | 0.2584 | 0.3442 | ✓ |
| 6 and 8 | | 460 | 95 | 4 | 0.1704 | 0.2464 | ✓ |
| 6 and 8 | [2000,3000) | 785 | 185 | 1 | 0.2064 | 0.267 | |
| 6 and 8 | | 785 | 198 | 2 | 0.2222 | 0.2841 | |
| 6 and 8 | | 785 | 228 | 3 | 0.2589 | 0.3236 | ✓ |
| 6 and 8 | | 785 | 174 | 4 | 0.1931 | 0.2524 | |
| 6 and 8 | [3000,5000) | 1804 | 449 | 1 | 0.2291 | 0.2695 | |
| 6 and 8 | | 1804 | 441 | 2 | 0.2248 | 0.265 | |
| 6 and 8 | | 1804 | 421 | 3 | 0.214 | 0.2536 | |
| 6 and 8 | | 1804 | 493 | 4 | 0.2528 | 0.2945 | ✓ |
| 6 and 8 | [5000,10000) | 6333 | 1513 | 1 | 0.2284 | 0.2496 | ✓ |
| 6 and 8 | | 6333 | 1592 | 2 | 0.2407 | 0.2623 | |
| 6 and 8 | | 6333 | 1576 | 3 | 0.2382 | 0.2597 | |
| 6 and 8 | | 6333 | 1652 | 4 | 0.2501 | 0.2719 | ✓ |

| Pair | Distance interval (m) | Total | Count | Type | Lower CI | Upper CI | Sign. diff. |
|---|---|---|---|---|---|---|---|
| 6 and 9 | [0,50) | 51 | 12 | 1 | 0.1279 | 0.3749 | |
| 6 and 9 | | 51 | 2 | 2 | 0.0048 | 0.1346 | ✓ |
| 6 and 9 | | 51 | 12 | 3 | 0.1279 | 0.3749 | |
| 6 and 9 | | 51 | 25 | 4 | 0.3475 | 0.634 | ✓ |
| 6 and 9 | [50,100) | 17 | 3 | 1 | 0.038 | 0.4343 | |
| 6 and 9 | | 17 | 2 | 2 | 0.0146 | 0.3644 | |
| 6 and 9 | | 17 | 5 | 3 | 0.1031 | 0.5596 | |
| 6 and 9 | | 17 | 7 | 4 | 0.1844 | 0.6708 | |
| 6 and 9 | [100,200) | 65 | 9 | 1 | 0.0653 | 0.2466 | ✓ |
| 6 and 9 | | 65 | 11 | 2 | 0.0876 | 0.2827 | |
| 6 and 9 | | 65 | 22 | 3 | 0.2257 | 0.4665 | |
| 6 and 9 | | 65 | 23 | 4 | 0.2392 | 0.4823 | |
| 6 and 9 | [200,500) | 182 | 27 | 1 | 0.1001 | 0.2085 | ✓ |
| 6 and 9 | | 182 | 38 | 2 | 0.1522 | 0.2751 | |
| 6 and 9 | | 182 | 55 | 3 | 0.2365 | 0.3745 | |
| 6 and 9 | | 182 | 62 | 4 | 0.2722 | 0.4144 | ✓ |
| 6 and 9 | [500,1000) | 378 | 67 | 1 | 0.1401 | 0.2196 | ✓ |
| 6 and 9 | | 378 | 95 | 2 | 0.2084 | 0.2982 | |
| 6 and 9 | | 378 | 116 | 3 | 0.2607 | 0.3561 | ✓ |
| 6 and 9 | | 378 | 100 | 4 | 0.2208 | 0.3121 | |
| 6 and 9 | [1000,2000) | 1024 | 237 | 1 | 0.2059 | 0.2585 | |
| 6 and 9 | | 1024 | 243 | 2 | 0.2115 | 0.2646 | |
| 6 and 9 | | 1024 | 266 | 3 | 0.2331 | 0.2878 | |
| 6 and 9 | | 1024 | 278 | 4 | 0.2444 | 0.2998 | |
| 6 and 9 | [2000,3000) | 1256 | 276 | 1 | 0.1971 | 0.2437 | ✓ |
| 6 and 9 | | 1256 | 283 | 2 | 0.2025 | 0.2495 | ✓ |
| 6 and 9 | | 1256 | 342 | 3 | 0.2478 | 0.2978 | |
| 6 and 9 | | 1256 | 355 | 4 | 0.2579 | 0.3084 | ✓ |
| 6 and 9 | [3000,5000) | 2940 | 785 | 1 | 0.2511 | 0.2834 | ✓ |
| 6 and 9 | | 2940 | 730 | 2 | 0.2328 | 0.2643 | |
| 6 and 9 | | 2940 | 745 | 3 | 0.2378 | 0.2695 | |
| 6 and 9 | | 2940 | 680 | 4 | 0.2162 | 0.247 | ✓ |
| 6 and 9 | [5000,10000) | 8171 | 2236 | 1 | 0.264 | 0.2835 | ✓ |
| 6 and 9 | | 8171 | 1884 | 2 | 0.2215 | 0.2399 | ✓ |
| 6 and 9 | | 8171 | 1940 | 3 | 0.2282 | 0.2468 | ✓ |
| 6 and 9 | | 8171 | 2111 | 4 | 0.2489 | 0.268 | |

| Pair | Distance interval (m) | Total | Count | Type | Lower CI | Upper CI | Sign. diff. |
|---|---|---|---|---|---|---|---|
| 6 and 10 | [0,50) | 113 | 22 | 1 | 0.1262 | 0.2798 | |
| 6 and 10 | | 113 | 18 | 2 | 0.0972 | 0.24 | ✓ |
| 6 and 10 | | 113 | 52 | 3 | 0.366 | 0.5565 | ✓ |
| 6 and 10 | | 113 | 21 | 4 | 0.1189 | 0.2699 | |
| 6 and 10 | [50,100) | 91 | 13 | 1 | 0.0783 | 0.2319 | ✓ |
| 6 and 10 | | 91 | 12 | 2 | 0.07 | 0.219 | ✓ |
| 6 and 10 | | 91 | 45 | 3 | 0.388 | 0.6014 | ✓ |
| 6 and 10 | | 91 | 21 | 4 | 0.1489 | 0.3309 | |
| 6 and 10 | [100,200) | 96 | 18 | 1 | 0.1151 | 0.28 | |
| 6 and 10 | | 96 | 14 | 2 | 0.0821 | 0.2326 | ✓ |
| 6 and 10 | | 96 | 43 | 3 | 0.3463 | 0.5529 | ✓ |
| 6 and 10 | | 96 | 21 | 4 | 0.1408 | 0.3147 | |
| 6 and 10 | [200,500) | 296 | 61 | 1 | 0.1615 | 0.2567 | |
| 6 and 10 | | 296 | 46 | 2 | 0.1161 | 0.2018 | ✓ |
| 6 and 10 | | 296 | 125 | 3 | 0.3654 | 0.4808 | ✓ |
| 6 and 10 | | 296 | 64 | 4 | 0.1707 | 0.2675 | |
| 6 and 10 | [500,1000) | 451 | 97 | 1 | 0.178 | 0.2559 | |
| 6 and 10 | | 451 | 95 | 2 | 0.1739 | 0.2512 | |
| 6 and 10 | | 451 | 157 | 3 | 0.3042 | 0.3941 | ✓ |
| 6 and 10 | | 451 | 102 | 4 | 0.1883 | 0.2676 | |
| 6 and 10 | [1000,2000) | 1126 | 282 | 1 | 0.2254 | 0.2768 | |
| 6 and 10 | | 1126 | 292 | 2 | 0.2339 | 0.286 | |
| 6 and 10 | | 1126 | 290 | 3 | 0.2322 | 0.2841 | |
| 6 and 10 | | 1126 | 262 | 4 | 0.2083 | 0.2585 | |
| 6 and 10 | [2000,3000) | 1415 | 360 | 1 | 0.2319 | 0.278 | |
| 6 and 10 | | 1415 | 389 | 2 | 0.2518 | 0.299 | ✓ |
| 6 and 10 | | 1415 | 335 | 3 | 0.2148 | 0.2598 | |
| 6 and 10 | | 1415 | 331 | 4 | 0.2121 | 0.2569 | |
| 6 and 10 | [3000,5000) | 3649 | 975 | 1 | 0.2529 | 0.2819 | ✓ |
| 6 and 10 | | 3649 | 833 | 2 | 0.2147 | 0.2423 | ✓ |
| 6 and 10 | | 3649 | 916 | 3 | 0.237 | 0.2654 | |
| 6 and 10 | | 3649 | 925 | 4 | 0.2394 | 0.2679 | |
| 6 and 10 | [5000,10000) | 8987 | 2412 | 1 | 0.2592 | 0.2777 | ✓ |
| 6 and 10 | | 8987 | 2128 | 2 | 0.228 | 0.2457 | ✓ |
| 6 and 10 | | 8987 | 2369 | 3 | 0.2545 | 0.2728 | ✓ |
| 6 and 10 | | 8987 | 2078 | 4 | 0.2225 | 0.2401 | ✓ |

| Pair | Distance interval (m) | Total | Count | Type | Lower CI | Upper CI | Sign. diff. |
|---|---|---|---|---|---|---|---|
| 7 and 8 | [0,50) | 512 | 64 | 1 | 0.0976 | 0.1568 | ✓ |
| 7 and 8 | | 512 | 44 | 2 | 0.0631 | 0.1137 | ✓ |
| 7 and 8 | | 512 | 210 | 3 | 0.3672 | 0.4542 | ✓ |
| 7 and 8 | | 512 | 194 | 4 | 0.3367 | 0.4225 | ✓ |
| 7 and 8 | [50,100) | 395 | 61 | 1 | 0.1202 | 0.1939 | ✓ |
| 7 and 8 | | 395 | 39 | 2 | 0.0712 | 0.1325 | ✓ |
| 7 and 8 | | 395 | 180 | 3 | 0.4058 | 0.5063 | ✓ |
| 7 and 8 | | 395 | 115 | 4 | 0.2468 | 0.3387 | |
| 7 and 8 | [100,200) | 409 | 68 | 1 | 0.1315 | 0.206 | ✓ |
| 7 and 8 | | 409 | 53 | 2 | 0.0986 | 0.1661 | ✓ |
| 7 and 8 | | 409 | 168 | 3 | 0.3627 | 0.4602 | ✓ |
| 7 and 8 | | 409 | 120 | 4 | 0.2497 | 0.3401 | |
| 7 and 8 | [200,500) | 643 | 125 | 1 | 0.1645 | 0.2271 | ✓ |
| 7 and 8 | | 643 | 101 | 2 | 0.1298 | 0.1875 | ✓ |
| 7 and 8 | | 643 | 222 | 3 | 0.3085 | 0.3834 | ✓ |
| 7 and 8 | | 643 | 195 | 4 | 0.2679 | 0.3404 | ✓ |
| 7 and 8 | [500,1000) | 916 | 187 | 1 | 0.1785 | 0.2317 | ✓ |
| 7 and 8 | | 916 | 181 | 2 | 0.1723 | 0.2249 | ✓ |
| 7 and 8 | | 916 | 289 | 3 | 0.2855 | 0.3467 | ✓ |
| 7 and 8 | | 916 | 259 | 4 | 0.2538 | 0.3131 | ✓ |
| 7 and 8 | [1000,2000) | 1752 | 413 | 1 | 0.216 | 0.2563 | |
| 7 and 8 | | 1752 | 404 | 2 | 0.211 | 0.251 | |
| 7 and 8 | | 1752 | 479 | 3 | 0.2526 | 0.2949 | ✓ |
| 7 and 8 | | 1752 | 456 | 4 | 0.2399 | 0.2815 | |
| 7 and 8 | [2000,3000) | 1767 | 436 | 1 | 0.2268 | 0.2675 | |
| 7 and 8 | | 1767 | 453 | 2 | 0.2361 | 0.2774 | |
| 7 and 8 | | 1767 | 420 | 3 | 0.218 | 0.2582 | |
| 7 and 8 | | 1767 | 458 | 4 | 0.2389 | 0.2803 | |
| 7 and 8 | [3000,5000) | 4009 | 974 | 1 | 0.2297 | 0.2565 | |
| 7 and 8 | | 4009 | 1069 | 2 | 0.253 | 0.2806 | ✓ |
| 7 and 8 | | 4009 | 938 | 3 | 0.2209 | 0.2474 | ✓ |
| 7 and 8 | | 4009 | 1028 | 4 | 0.243 | 0.2702 | |
| 7 and 8 | [5000,10000) | 9591 | 2466 | 1 | 0.2484 | 0.266 | |
| 7 and 8 | | 9591 | 2389 | 2 | 0.2405 | 0.2579 | |
| 7 and 8 | | 9591 | 2089 | 3 | 0.2096 | 0.2262 | ✓ |
| 7 and 8 | | 9591 | 2647 | 4 | 0.2671 | 0.2851 | ✓ |

| Pair | Distance interval (m) | Total | Count | Type | Lower CI | Upper CI | Sign. diff. |
|---|---|---|---|---|---|---|---|
| 7 and 9 | [0,50) | 102 | 20 | 1 | 0.1241 | 0.2865 | |
| 7 and 9 | | 102 | 5 | 2 | 0.0161 | 0.1107 | ✓ |
| 7 and 9 | | 102 | 35 | 3 | 0.2519 | 0.4437 | ✓ |
| 7 and 9 | | 102 | 42 | 4 | 0.3152 | 0.5136 | ✓ |
| 7 and 9 | [50,100) | 75 | 10 | 1 | 0.0658 | 0.2316 | ✓ |
| 7 and 9 | | 75 | 9 | 2 | 0.0564 | 0.2156 | ✓ |
| 7 and 9 | | 75 | 19 | 3 | 0.1599 | 0.367 | |
| 7 and 9 | | 75 | 37 | 4 | 0.3758 | 0.6114 | ✓ |
| 7 and 9 | [100,200) | 73 | 9 | 1 | 0.058 | 0.2212 | ✓ |
| 7 and 9 | | 73 | 7 | 2 | 0.0394 | 0.1876 | ✓ |
| 7 and 9 | | 73 | 22 | 3 | 0.1994 | 0.42 | |
| 7 and 9 | | 73 | 35 | 4 | 0.361 | 0.5996 | ✓ |
| 7 and 9 | [200,500) | 200 | 27 | 1 | 0.0909 | 0.1903 | ✓ |
| 7 and 9 | | 200 | 27 | 2 | 0.0909 | 0.1903 | ✓ |
| 7 and 9 | | 200 | 74 | 3 | 0.303 | 0.4409 | ✓ |
| 7 and 9 | | 200 | 72 | 4 | 0.2935 | 0.4307 | ✓ |
| 7 and 9 | [500,1000) | 373 | 75 | 1 | 0.1616 | 0.2454 | ✓ |
| 7 and 9 | | 373 | 92 | 2 | 0.2037 | 0.2936 | |
| 7 and 9 | | 373 | 98 | 3 | 0.2188 | 0.3105 | |
| 7 and 9 | | 373 | 108 | 4 | 0.244 | 0.3385 | |
| 7 and 9 | [1000,2000) | 823 | 204 | 1 | 0.2187 | 0.2789 | |
| 7 and 9 | | 823 | 176 | 2 | 0.1863 | 0.2435 | ✓ |
| 7 and 9 | | 823 | 221 | 3 | 0.2385 | 0.3002 | |
| 7 and 9 | | 823 | 222 | 4 | 0.2397 | 0.3015 | |
| 7 and 9 | [2000,3000) | 1198 | 283 | 1 | 0.2124 | 0.2613 | |
| 7 and 9 | | 1198 | 271 | 2 | 0.2028 | 0.251 | |
| 7 and 9 | | 1198 | 317 | 3 | 0.2398 | 0.2906 | |
| 7 and 9 | | 1198 | 327 | 4 | 0.2479 | 0.2991 | |
| 7 and 9 | [3000,5000) | 2800 | 693 | 1 | 0.2316 | 0.2639 | |
| 7 and 9 | | 2800 | 702 | 2 | 0.2347 | 0.2672 | |
| 7 and 9 | | 2800 | 748 | 3 | 0.2508 | 0.2839 | ✓ |
| 7 and 9 | | 2800 | 657 | 4 | 0.2191 | 0.2508 | |
| 7 and 9 | [5000,10000) | 7627 | 1896 | 1 | 0.2389 | 0.2585 | |
| 7 and 9 | | 7627 | 1824 | 2 | 0.2296 | 0.2489 | ✓ |
| 7 and 9 | | 7627 | 1944 | 3 | 0.2451 | 0.2648 | |
| 7 and 9 | | 7627 | 1963 | 4 | 0.2476 | 0.2673 | |

| Pair | Distance interval (m) | Total | Count | Type | Lower CI | Upper CI | Sign. diff. |
|---|---|---|---|---|---|---|---|
| 7 and 10 | [0,50) | 30 | 7 | 1 | 0.0993 | 0.4228 | |
| 7 and 10 | | 30 | 3 | 2 | 0.0211 | 0.2653 | |
| 7 and 10 | | 30 | 6 | 3 | 0.0771 | 0.3857 | |
| 7 and 10 | | 30 | 14 | 4 | 0.2834 | 0.6567 | ✓ |
| 7 and 10 | [50,100) | 28 | 6 | 1 | 0.083 | 0.4095 | |
| 7 and 10 | | 28 | 3 | 2 | 0.0227 | 0.2823 | |
| 7 and 10 | | 28 | 9 | 3 | 0.1588 | 0.5235 | |
| 7 and 10 | | 28 | 10 | 4 | 0.1864 | 0.5593 | |
| 7 and 10 | [100,200) | 22 | 6 | 1 | 0.1073 | 0.5022 | |
| 7 and 10 | | 22 | 3 | 2 | 0.0291 | 0.3491 | |
| 7 and 10 | | 22 | 10 | 3 | 0.2439 | 0.6779 | |
| 7 and 10 | | 22 | 3 | 4 | 0.0291 | 0.3491 | |
| 7 and 10 | [200,500) | 106 | 20 | 1 | 0.1192 | 0.2762 | |
| 7 and 10 | | 106 | 21 | 2 | 0.127 | 0.2868 | |
| 7 and 10 | | 106 | 47 | 3 | 0.3469 | 0.5431 | ✓ |
| 7 and 10 | | 106 | 18 | 4 | 0.1039 | 0.255 | |
| 7 and 10 | [500,1000) | 240 | 50 | 1 | 0.1588 | 0.2653 | |
| 7 and 10 | | 240 | 67 | 2 | 0.2234 | 0.3405 | |
| 7 and 10 | | 240 | 74 | 3 | 0.2505 | 0.371 | ✓ |
| 7 and 10 | | 240 | 49 | 4 | 0.155 | 0.2608 | |
| 7 and 10 | [1000,2000) | 515 | 91 | 1 | 0.1447 | 0.2124 | ✓ |
| 7 and 10 | | 515 | 127 | 2 | 0.2099 | 0.2862 | |
| 7 and 10 | | 515 | 164 | 3 | 0.2784 | 0.3606 | ✓ |
| 7 and 10 | | 515 | 133 | 4 | 0.221 | 0.2983 | |
| 7 and 10 | [2000,3000) | 937 | 203 | 1 | 0.1907 | 0.2444 | ✓ |
| 7 and 10 | | 937 | 223 | 2 | 0.2111 | 0.2666 | |
| 7 and 10 | | 937 | 307 | 3 | 0.2976 | 0.3587 | ✓ |
| 7 and 10 | | 937 | 204 | 4 | 0.1917 | 0.2455 | ✓ |
| 7 and 10 | [3000,5000) | 2464 | 579 | 1 | 0.2184 | 0.2522 | |
| 7 and 10 | | 2464 | 583 | 2 | 0.2199 | 0.2539 | |
| 7 and 10 | | 2464 | 723 | 3 | 0.2755 | 0.3118 | ✓ |
| 7 and 10 | | 2464 | 579 | 4 | 0.2184 | 0.2522 | |
| 7 and 10 | [5000,10000) | 8433 | 2060 | 1 | 0.2351 | 0.2536 | |
| 7 and 10 | | 8433 | 2121 | 2 | 0.2423 | 0.2609 | |
| 7 and 10 | | 8433 | 2205 | 3 | 0.2521 | 0.271 | ✓ |
| 7 and 10 | | 8433 | 2047 | 4 | 0.2336 | 0.252 | |

| Pair | Distance interval (m) | Total | Count | Type | Lower CI | Upper CI | Sign. diff. |
|---|---|---|---|---|---|---|---|
| 8 and 9 | [0,50) | 26 | 7 | 1 | 0.1157 | 0.4779 | |
| 8 and 9 | | 26 | 1 | 2 | 0.001 | 0.1964 | ✓ |
| 8 and 9 | | 26 | 9 | 3 | 0.1721 | 0.5567 | |
| 8 and 9 | | 26 | 9 | 4 | 0.1721 | 0.5567 | |
| 8 and 9 | [50,100) | 22 | 3 | 1 | 0.0291 | 0.3491 | |
| 8 and 9 | | 22 | 1 | 2 | 0.0012 | 0.2284 | ✓ |
| 8 and 9 | | 22 | 7 | 3 | 0.1386 | 0.5487 | |
| 8 and 9 | | 22 | 11 | 4 | 0.2822 | 0.7178 | ✓ |
| 8 and 9 | [100,200) | 26 | 4 | 1 | 0.0436 | 0.3487 | |
| 8 and 9 | | 26 | 1 | 2 | 0.001 | 0.1964 | ✓ |
| 8 and 9 | | 26 | 12 | 3 | 0.2659 | 0.6663 | ✓ |
| 8 and 9 | | 26 | 9 | 4 | 0.1721 | 0.5567 | |
| 8 and 9 | [200,500) | 88 | 15 | 1 | 0.0987 | 0.2655 | |
| 8 and 9 | | 88 | 13 | 2 | 0.0811 | 0.2394 | ✓ |
| 8 and 9 | | 88 | 21 | 3 | 0.1542 | 0.3414 | |
| 8 and 9 | | 88 | 39 | 4 | 0.3373 | 0.553 | ✓ |
| 8 and 9 | [500,1000) | 236 | 40 | 1 | 0.1239 | 0.2236 | ✓ |
| 8 and 9 | | 236 | 51 | 2 | 0.1653 | 0.2741 | |
| 8 and 9 | | 236 | 59 | 3 | 0.1961 | 0.3103 | |
| 8 and 9 | | 236 | 86 | 4 | 0.3029 | 0.4293 | ✓ |
| 8 and 9 | [1000,2000) | 479 | 94 | 1 | 0.1616 | 0.2347 | ✓ |
| 8 and 9 | | 479 | 114 | 2 | 0.2005 | 0.2787 | |
| 8 and 9 | | 479 | 152 | 3 | 0.2758 | 0.3611 | ✓ |
| 8 and 9 | | 479 | 119 | 4 | 0.2103 | 0.2897 | |
| 8 and 9 | [2000,3000) | 595 | 115 | 1 | 0.1623 | 0.2273 | ✓ |
| 8 and 9 | | 595 | 184 | 2 | 0.2723 | 0.3481 | ✓ |
| 8 and 9 | | 595 | 173 | 3 | 0.2545 | 0.3291 | ✓ |
| 8 and 9 | | 595 | 123 | 4 | 0.1749 | 0.2415 | ✓ |
| 8 and 9 | [3000,5000) | 1666 | 400 | 1 | 0.2198 | 0.2614 | |
| 8 and 9 | | 1666 | 488 | 2 | 0.2711 | 0.3154 | ✓ |
| 8 and 9 | | 1666 | 405 | 3 | 0.2227 | 0.2644 | |
| 8 and 9 | | 1666 | 373 | 4 | 0.2041 | 0.2447 | ✓ |
| 8 and 9 | [5000,10000) | 7888 | 1944 | 1 | 0.237 | 0.2561 | |
| 8 and 9 | | 7888 | 2065 | 2 | 0.2521 | 0.2716 | ✓ |
| 8 and 9 | | 7888 | 2026 | 3 | 0.2472 | 0.2666 | |
| 8 and 9 | | 7888 | 1853 | 4 | 0.2256 | 0.2444 | ✓ |

| Pair | Distance interval (m) | Total | Count | Type | Lower CI | Upper CI | Sign. diff. |
|---|---|---|---|---|---|---|---|
| 9 and 10 | [0,50) | 69 | 10 | 1 | 0.0717 | 0.2504 | |
| 9 and 10 | | 69 | 9 | 2 | 0.0614 | 0.2332 | ✓ |
| 9 and 10 | | 69 | 39 | 3 | 0.4404 | 0.6842 | ✓ |
| 9 and 10 | | 69 | 11 | 4 | 0.0824 | 0.2674 | |
| 9 and 10 | [50,100) | 52 | 12 | 1 | 0.1253 | 0.3684 | |
| 9 and 10 | | 52 | 9 | 2 | 0.0823 | 0.3033 | |
| 9 and 10 | | 52 | 20 | 3 | 0.253 | 0.5298 | ✓ |
| 9 and 10 | | 52 | 11 | 4 | 0.1106 | 0.347 | |
| 9 and 10 | [100,200) | 92 | 20 | 1 | 0.1381 | 0.3156 | |
| 9 and 10 | | 92 | 10 | 2 | 0.0534 | 0.1908 | ✓ |
| 9 and 10 | | 92 | 44 | 3 | 0.373 | 0.585 | ✓ |
| 9 and 10 | | 92 | 18 | 4 | 0.1203 | 0.2915 | |
| 9 and 10 | [200,500) | 272 | 59 | 1 | 0.1694 | 0.2707 | |
| 9 and 10 | | 272 | 52 | 2 | 0.1462 | 0.243 | ✓ |
| 9 and 10 | | 272 | 85 | 3 | 0.2579 | 0.3713 | ✓ |
| 9 and 10 | | 272 | 76 | 4 | 0.2269 | 0.3368 | |
| 9 and 10 | [500,1000) | 676 | 144 | 1 | 0.1827 | 0.2458 | ✓ |
| 9 and 10 | | 676 | 156 | 2 | 0.1995 | 0.2644 | |
| 9 and 10 | | 676 | 222 | 3 | 0.2931 | 0.3652 | ✓ |
| 9 and 10 | | 676 | 154 | 4 | 0.1967 | 0.2613 | |
| 9 and 10 | [1000,2000) | 1694 | 381 | 1 | 0.2052 | 0.2456 | ✓ |
| 9 and 10 | | 1694 | 406 | 2 | 0.2195 | 0.2607 | |
| 9 and 10 | | 1694 | 544 | 3 | 0.2989 | 0.344 | ✓ |
| 9 and 10 | | 1694 | 363 | 4 | 0.195 | 0.2346 | ✓ |
| 9 and 10 | [2000,3000) | 2233 | 591 | 1 | 0.2465 | 0.2835 | |
| 9 and 10 | | 2233 | 502 | 2 | 0.2076 | 0.2427 | ✓ |
| 9 and 10 | | 2233 | 595 | 3 | 0.2482 | 0.2853 | |
| 9 and 10 | | 2233 | 545 | 4 | 0.2264 | 0.2624 | |
| 9 and 10 | [3000,5000) | 4775 | 1227 | 1 | 0.2446 | 0.2696 | |
| 9 and 10 | | 4775 | 1123 | 2 | 0.2232 | 0.2475 | ✓ |
| 9 and 10 | | 4775 | 1253 | 3 | 0.25 | 0.2751 | |
| 9 and 10 | | 4775 | 1172 | 4 | 0.2333 | 0.2579 | |
| 9 and 10 | [5000,10000) | 11216 | 2911 | 1 | 0.2514 | 0.2678 | ✓ |
| 9 and 10 | | 11216 | 2666 | 2 | 0.2298 | 0.2457 | ✓ |
| 9 and 10 | | 11216 | 2995 | 3 | 0.2589 | 0.2753 | ✓ |
| 9 and 10 | | 11216 | 2644 | 4 | 0.2279 | 0.2437 | ✓ |

### *6.2. 20-min pairs*

| Pair | Distance interval (m) | Total | Count | Type | Lower CI | Upper CI | Sign. diff. |
|---|---|---|---|---|---|---|---|
| 1 and 3 | [0,50) | 324 | 51 | 1 | 0.1195 | 0.2017 | ✓ |
| 1 and 3 | | 324 | 23 | 2 | 0.0455 | 0.1046 | ✓ |
| 1 and 3 | | 324 | 136 | 3 | 0.3654 | 0.4756 | ✓ |
| 1 and 3 | | 324 | 114 | 4 | 0.2999 | 0.4066 | ✓ |
| 1 and 3 | [50,100) | 121 | 19 | 1 | 0.0973 | 0.2343 | ✓ |
| 1 and 3 | | 121 | 8 | 2 | 0.029 | 0.1261 | ✓ |
| 1 and 3 | | 121 | 58 | 3 | 0.3877 | 0.572 | ✓ |
| 1 and 3 | | 121 | 36 | 4 | 0.2179 | 0.3874 | |
| 1 and 3 | [100,200) | 96 | 11 | 1 | 0.0586 | 0.1958 | ✓ |
| 1 and 3 | | 96 | 8 | 2 | 0.0367 | 0.1576 | ✓ |
| 1 and 3 | | 96 | 44 | 3 | 0.3562 | 0.5631 | ✓ |
| 1 and 3 | | 96 | 33 | 4 | 0.2498 | 0.4477 | |
| 1 and 3 | [200,500) | 172 | 28 | 1 | 0.111 | 0.2266 | ✓ |
| 1 and 3 | | 172 | 25 | 2 | 0.0963 | 0.207 | ✓ |
| 1 and 3 | | 172 | 62 | 3 | 0.2888 | 0.437 | ✓ |
| 1 and 3 | | 172 | 57 | 4 | 0.2616 | 0.4071 | ✓ |
| 1 and 3 | [500,1000) | 261 | 60 | 1 | 0.1803 | 0.2858 | |
| 1 and 3 | | 261 | 46 | 2 | 0.132 | 0.228 | ✓ |
| 1 and 3 | | 261 | 65 | 3 | 0.1978 | 0.3061 | |
| 1 and 3 | | 261 | 90 | 4 | 0.2873 | 0.4059 | ✓ |
| 1 and 3 | [1000,2000) | 576 | 122 | 1 | 0.1791 | 0.2475 | ✓ |
| 1 and 3 | | 576 | 117 | 2 | 0.171 | 0.2383 | ✓ |
| 1 and 3 | | 576 | 167 | 3 | 0.2532 | 0.3289 | ✓ |
| 1 and 3 | | 576 | 170 | 4 | 0.2582 | 0.3342 | ✓ |
| 1 and 3 | [2000,3000) | 677 | 176 | 1 | 0.2273 | 0.2948 | |
| 1 and 3 | | 677 | 189 | 2 | 0.2457 | 0.3146 | |
| 1 and 3 | | 677 | 157 | 3 | 0.2006 | 0.2656 | |
| 1 and 3 | | 677 | 155 | 4 | 0.1978 | 0.2625 | |
| 1 and 3 | [3000,5000) | 1657 | 378 | 1 | 0.2081 | 0.2491 | ✓ |
| 1 and 3 | | 1657 | 423 | 2 | 0.2344 | 0.277 | |
| 1 and 3 | | 1657 | 444 | 3 | 0.2468 | 0.29 | |
| 1 and 3 | | 1657 | 412 | 4 | 0.228 | 0.2702 | |
| 1 and 3 | [5000,10000) | 2650 | 600 | 1 | 0.2106 | 0.2428 | ✓ |
| 1 and 3 | | 2650 | 717 | 2 | 0.2537 | 0.2879 | ✓ |
| 1 and 3 | | 2650 | 738 | 3 | 0.2615 | 0.296 | ✓ |
| 1 and 3 | | 2650 | 595 | 4 | 0.2088 | 0.2409 | ✓ |

| Pair | Distance interval (m) | Total | Count | Type | Lower CI | Upper CI | Sign. diff. |
|---|---|---|---|---|---|---|---|
| 1 and 4 | [0,50) | 171 | 35 | 1 | 0.1469 | 0.273 | |
| 1 and 4 | | 171 | 12 | 2 | 0.0368 | 0.1194 | ✓ |
| 1 and 4 | | 171 | 58 | 3 | 0.2687 | 0.4154 | ✓ |
| 1 and 4 | | 171 | 66 | 4 | 0.3126 | 0.4633 | ✓ |
| 1 and 4 | [50,100) | 123 | 21 | 1 | 0.1089 | 0.2491 | ✓ |
| 1 and 4 | | 123 | 7 | 2 | 0.0232 | 0.1137 | ✓ |
| 1 and 4 | | 123 | 38 | 3 | 0.2288 | 0.3986 | |
| 1 and 4 | | 123 | 57 | 4 | 0.3731 | 0.5556 | ✓ |
| 1 and 4 | [100,200) | 113 | 14 | 1 | 0.0694 | 0.1991 | ✓ |
| 1 and 4 | | 113 | 17 | 2 | 0.0901 | 0.2299 | ✓ |
| 1 and 4 | | 113 | 30 | 3 | 0.1868 | 0.3568 | |
| 1 and 4 | | 113 | 52 | 4 | 0.366 | 0.5565 | ✓ |
| 1 and 4 | [200,500) | 133 | 19 | 1 | 0.0883 | 0.2141 | ✓ |
| 1 and 4 | | 133 | 27 | 2 | 0.1383 | 0.2814 | |
| 1 and 4 | | 133 | 40 | 3 | 0.2243 | 0.3863 | |
| 1 and 4 | | 133 | 47 | 4 | 0.2725 | 0.4409 | ✓ |
| 1 and 4 | [500,1000) | 228 | 44 | 1 | 0.1439 | 0.2503 | |
| 1 and 4 | | 228 | 45 | 2 | 0.1478 | 0.255 | |
| 1 and 4 | | 228 | 50 | 3 | 0.1674 | 0.2787 | |
| 1 and 4 | | 228 | 89 | 4 | 0.3266 | 0.457 | ✓ |
| 1 and 4 | [1000,2000) | 433 | 111 | 1 | 0.2159 | 0.3002 | |
| 1 and 4 | | 433 | 86 | 2 | 0.1621 | 0.2394 | ✓ |
| 1 and 4 | | 433 | 124 | 3 | 0.2442 | 0.3315 | |
| 1 and 4 | | 433 | 112 | 4 | 0.218 | 0.3026 | |
| 1 and 4 | [2000,3000) | 224 | 49 | 1 | 0.1664 | 0.2787 | |
| 1 and 4 | | 224 | 63 | 2 | 0.2234 | 0.345 | |
| 1 and 4 | | 224 | 64 | 3 | 0.2275 | 0.3497 | |
| 1 and 4 | | 224 | 48 | 4 | 0.1624 | 0.2739 | |
| 1 and 4 | [3000,5000) | 773 | 206 | 1 | 0.2356 | 0.2992 | |
| 1 and 4 | | 773 | 188 | 2 | 0.2133 | 0.275 | |
| 1 and 4 | | 773 | 206 | 3 | 0.2356 | 0.2992 | |
| 1 and 4 | | 773 | 173 | 4 | 0.1949 | 0.2549 | |
| 1 and 4 | [5000,10000) | 1255 | 336 | 1 | 0.2434 | 0.2931 | |
| 1 and 4 | | 1255 | 317 | 2 | 0.2288 | 0.2776 | |
| 1 and 4 | | 1255 | 322 | 3 | 0.2326 | 0.2817 | |
| 1 and 4 | | 1255 | 280 | 4 | 0.2003 | 0.2472 | ✓ |

| Pair | Distance interval (m) | Total | Count | Type | Lower CI | Upper CI | Sign. diff. |
|---|---|---|---|---|---|---|---|
| 1 and 5 | [0,50) | 155 | 23 | 1 | 0.0964 | 0.2143 | ✓ |
| 1 and 5 | | 155 | 17 | 2 | 0.0652 | 0.1698 | ✓ |
| 1 and 5 | | 155 | 43 | 3 | 0.2086 | 0.355 | |
| 1 and 5 | | 155 | 72 | 4 | 0.3841 | 0.5463 | ✓ |
| 1 and 5 | [50,100) | 105 | 24 | 1 | 0.1523 | 0.3207 | |
| 1 and 5 | | 105 | 10 | 2 | 0.0466 | 0.1682 | ✓ |
| 1 and 5 | | 105 | 37 | 3 | 0.2616 | 0.4517 | ✓ |
| 1 and 5 | | 105 | 34 | 4 | 0.2357 | 0.4221 | |
| 1 and 5 | [100,200) | 107 | 18 | 1 | 0.1029 | 0.2528 | |
| 1 and 5 | | 107 | 15 | 2 | 0.0806 | 0.2207 | ✓ |
| 1 and 5 | | 107 | 30 | 3 | 0.1978 | 0.3755 | |
| 1 and 5 | | 107 | 44 | 4 | 0.317 | 0.5105 | ✓ |
| 1 and 5 | [200,500) | 170 | 22 | 1 | 0.0829 | 0.1894 | ✓ |
| 1 and 5 | | 170 | 28 | 2 | 0.1123 | 0.2292 | ✓ |
| 1 and 5 | | 170 | 55 | 3 | 0.2539 | 0.3994 | ✓ |
| 1 and 5 | | 170 | 65 | 4 | 0.309 | 0.4599 | ✓ |
| 1 and 5 | [500,1000) | 211 | 41 | 1 | 0.1432 | 0.2542 | |
| 1 and 5 | | 211 | 46 | 2 | 0.1643 | 0.2799 | |
| 1 and 5 | | 211 | 37 | 3 | 0.1266 | 0.2335 | ✓ |
| 1 and 5 | | 211 | 87 | 4 | 0.3452 | 0.482 | ✓ |
| 1 and 5 | [1000,2000) | 424 | 90 | 1 | 0.1743 | 0.2543 | |
| 1 and 5 | | 424 | 106 | 2 | 0.2095 | 0.2941 | |
| 1 and 5 | | 424 | 122 | 3 | 0.2451 | 0.3334 | |
| 1 and 5 | | 424 | 106 | 4 | 0.2095 | 0.2941 | |
| 1 and 5 | [2000,3000) | 375 | 98 | 1 | 0.2176 | 0.3089 | |
| 1 and 5 | | 375 | 88 | 2 | 0.1927 | 0.2809 | |
| 1 and 5 | | 375 | 86 | 3 | 0.1877 | 0.2753 | |
| 1 and 5 | | 375 | 103 | 4 | 0.2301 | 0.3228 | |
| 1 and 5 | [3000,5000) | 924 | 259 | 1 | 0.2515 | 0.3105 | ✓ |
| 1 and 5 | | 924 | 210 | 2 | 0.2006 | 0.2557 | |
| 1 and 5 | | 924 | 247 | 3 | 0.239 | 0.2971 | |
| 1 and 5 | | 924 | 208 | 4 | 0.1985 | 0.2534 | |
| 1 and 5 | [5000,10000) | 1544 | 427 | 1 | 0.2544 | 0.2996 | ✓ |
| 1 and 5 | | 1544 | 371 | 2 | 0.2192 | 0.2624 | |
| 1 and 5 | | 1544 | 402 | 3 | 0.2386 | 0.283 | |
| 1 and 5 | | 1544 | 344 | 4 | 0.2023 | 0.2444 | ✓ |

| Pair | Distance interval (m) | Total | Count | Type | Lower CI | Upper CI | Sign. diff. |
|---|---|---|---|---|---|---|---|
| 1 and 14 | [0,50) | 176 | 38 | 1 | 0.1576 | 0.2841 | |
| 1 and 14 | | 176 | 12 | 2 | 0.0357 | 0.1161 | ✓ |
| 1 and 14 | | 176 | 59 | 3 | 0.266 | 0.4101 | ✓ |
| 1 and 14 | | 176 | 67 | 4 | 0.3087 | 0.4568 | ✓ |
| 1 and 14 | [50,100) | 131 | 10 | 1 | 0.0372 | 0.1359 | ✓ |
| 1 and 14 | | 131 | 5 | 2 | 0.0125 | 0.0868 | ✓ |
| 1 and 14 | | 131 | 66 | 3 | 0.4152 | 0.5923 | ✓ |
| 1 and 14 | | 131 | 50 | 4 | 0.2982 | 0.4706 | ✓ |
| 1 and 14 | [100,200) | 101 | 10 | 1 | 0.0485 | 0.1746 | ✓ |
| 1 and 14 | | 101 | 6 | 2 | 0.0221 | 0.1248 | ✓ |
| 1 and 14 | | 101 | 36 | 3 | 0.2636 | 0.4579 | ✓ |
| 1 and 14 | | 101 | 49 | 4 | 0.3845 | 0.5867 | ✓ |
| 1 and 14 | [200,500) | 170 | 23 | 1 | 0.0877 | 0.1961 | ✓ |
| 1 and 14 | | 170 | 24 | 2 | 0.0926 | 0.2027 | ✓ |
| 1 and 14 | | 170 | 67 | 3 | 0.3202 | 0.4718 | ✓ |
| 1 and 14 | | 170 | 56 | 4 | 0.2594 | 0.4055 | ✓ |
| 1 and 14 | [500,1000) | 124 | 16 | 1 | 0.0756 | 0.2011 | ✓ |
| 1 and 14 | | 124 | 22 | 2 | 0.1147 | 0.2562 | |
| 1 and 14 | | 124 | 54 | 3 | 0.3467 | 0.5274 | ✓ |
| 1 and 14 | | 124 | 32 | 4 | 0.1837 | 0.3443 | |
| 1 and 14 | [1000,2000) | 278 | 51 | 1 | 0.1398 | 0.234 | ✓ |
| 1 and 14 | | 278 | 66 | 2 | 0.1886 | 0.2919 | |
| 1 and 14 | | 278 | 108 | 3 | 0.3309 | 0.4485 | ✓ |
| 1 and 14 | | 278 | 53 | 4 | 0.1462 | 0.2418 | ✓ |
| 1 and 14 | [2000,3000) | 272 | 65 | 1 | 0.1895 | 0.2942 | |
| 1 and 14 | | 272 | 63 | 2 | 0.1828 | 0.2864 | |
| 1 and 14 | | 272 | 84 | 3 | 0.2544 | 0.3674 | ✓ |
| 1 and 14 | | 272 | 60 | 4 | 0.1728 | 0.2746 | |
| 1 and 14 | [3000,5000) | 366 | 88 | 1 | 0.1975 | 0.2876 | |
| 1 and 14 | | 366 | 87 | 2 | 0.195 | 0.2847 | |
| 1 and 14 | | 366 | 97 | 3 | 0.2205 | 0.3134 | |
| 1 and 14 | | 366 | 94 | 4 | 0.2128 | 0.3048 | |
| 1 and 14 | [5000,10000) | 900 | 191 | 1 | 0.1859 | 0.2404 | ✓ |
| 1 and 14 | | 900 | 234 | 2 | 0.2316 | 0.29 | |
| 1 and 14 | | 900 | 264 | 3 | 0.2637 | 0.3243 | ✓ |
| 1 and 14 | | 900 | 211 | 4 | 0.2071 | 0.2635 | |

| Pair | Distance interval (m) | Total | Count | Type | Lower CI | Upper CI | Sign. diff. |
|---|---|---|---|---|---|---|---|
| 3 and 15 | [0,50) | 61 | 9 | 1 | 0.0698 | 0.2617 | |
| 3 and 15 | | 61 | 5 | 2 | 0.0272 | 0.181 | ✓ |
| 3 and 15 | | 61 | 21 | 3 | 0.2273 | 0.4769 | |
| 3 and 15 | | 61 | 26 | 4 | 0.3004 | 0.5594 | ✓ |
| 3 and 15 | [50,100) | 37 | 3 | 1 | 0.017 | 0.2191 | ✓ |
| 3 and 15 | | 37 | 5 | 2 | 0.0454 | 0.2877 | |
| 3 and 15 | | 37 | 14 | 3 | 0.2246 | 0.5524 | |
| 3 and 15 | | 37 | 15 | 4 | 0.2475 | 0.579 | |
| 3 and 15 | [100,200) | 49 | 6 | 1 | 0.0463 | 0.2477 | ✓ |
| 3 and 15 | | 49 | 1 | 2 | 5e-04 | 0.1085 | ✓ |
| 3 and 15 | | 49 | 23 | 3 | 0.3253 | 0.6173 | ✓ |
| 3 and 15 | | 49 | 19 | 4 | 0.252 | 0.5376 | ✓ |
| 3 and 15 | [200,500) | 132 | 25 | 1 | 0.1265 | 0.2668 | |
| 3 and 15 | | 132 | 23 | 2 | 0.1138 | 0.2499 | ✓ |
| 3 and 15 | | 132 | 46 | 3 | 0.2677 | 0.4363 | ✓ |
| 3 and 15 | | 132 | 38 | 4 | 0.2124 | 0.3731 | |
| 3 and 15 | [500,1000) | 240 | 37 | 1 | 0.1109 | 0.2062 | ✓ |
| 3 and 15 | | 240 | 49 | 2 | 0.155 | 0.2608 | |
| 3 and 15 | | 240 | 72 | 3 | 0.2427 | 0.3623 | |
| 3 and 15 | | 240 | 82 | 4 | 0.2819 | 0.4054 | ✓ |
| 3 and 15 | [1000,2000) | 539 | 123 | 1 | 0.1934 | 0.266 | |
| 3 and 15 | | 539 | 105 | 2 | 0.1622 | 0.2308 | ✓ |
| 3 and 15 | | 539 | 166 | 3 | 0.2692 | 0.3489 | ✓ |
| 3 and 15 | | 539 | 145 | 4 | 0.232 | 0.3086 | |
| 3 and 15 | [2000,3000) | 542 | 145 | 1 | 0.2307 | 0.3069 | |
| 3 and 15 | | 542 | 152 | 2 | 0.243 | 0.3203 | |
| 3 and 15 | | 542 | 143 | 3 | 0.2272 | 0.3031 | |
| 3 and 15 | | 542 | 102 | 4 | 0.1561 | 0.2237 | ✓ |
| 3 and 15 | [3000,5000) | 1356 | 332 | 1 | 0.2222 | 0.2686 | |
| 3 and 15 | | 1356 | 349 | 2 | 0.2343 | 0.2815 | |
| 3 and 15 | | 1356 | 361 | 3 | 0.2429 | 0.2906 | |
| 3 and 15 | | 1356 | 314 | 4 | 0.2093 | 0.255 | |
| 3 and 15 | [5000,10000) | 3446 | 862 | 1 | 0.2358 | 0.265 | |
| 3 and 15 | | 3446 | 834 | 2 | 0.2278 | 0.2567 | |
| 3 and 15 | | 3446 | 959 | 3 | 0.2634 | 0.2936 | ✓ |
| 3 and 15 | | 3446 | 791 | 4 | 0.2156 | 0.244 | ✓ |

| Pair | Distance interval (m) | Total | Count | Type | Lower CI | Upper CI | Sign. diff. |
|---|---|---|---|---|---|---|---|
| 4 and 5 | [0,50) | 2398 | 258 | 1 | 0.0955 | 0.1207 | ✓ |
| 4 and 5 | | 2398 | 163 | 2 | 0.0582 | 0.0788 | ✓ |
| 4 and 5 | | 2398 | 659 | 3 | 0.257 | 0.2932 | ✓ |
| 4 and 5 | | 2398 | 1318 | 4 | 0.5295 | 0.5697 | ✓ |
| 4 and 5 | [50,100) | 1414 | 141 | 1 | 0.0846 | 0.1165 | ✓ |
| 4 and 5 | | 1414 | 98 | 2 | 0.0566 | 0.0838 | ✓ |
| 4 and 5 | | 1414 | 456 | 3 | 0.2982 | 0.3476 | ✓ |
| 4 and 5 | | 1414 | 719 | 4 | 0.4821 | 0.5349 | ✓ |
| 4 and 5 | [100,200) | 1000 | 120 | 1 | 0.1005 | 0.1418 | ✓ |
| 4 and 5 | | 1000 | 105 | 2 | 0.0867 | 0.1257 | ✓ |
| 4 and 5 | | 1000 | 282 | 3 | 0.2543 | 0.311 | ✓ |
| 4 and 5 | | 1000 | 493 | 4 | 0.4616 | 0.5245 | ✓ |
| 4 and 5 | [200,500) | 634 | 114 | 1 | 0.1507 | 0.212 | ✓ |
| 4 and 5 | | 634 | 69 | 2 | 0.0857 | 0.1357 | ✓ |
| 4 and 5 | | 634 | 200 | 3 | 0.2794 | 0.3532 | ✓ |
| 4 and 5 | | 634 | 251 | 4 | 0.3576 | 0.4352 | ✓ |
| 4 and 5 | [500,1000) | 294 | 60 | 1 | 0.1595 | 0.2547 | |
| 4 and 5 | | 294 | 47 | 2 | 0.1199 | 0.2069 | ✓ |
| 4 and 5 | | 294 | 99 | 3 | 0.2829 | 0.3939 | ✓ |
| 4 and 5 | | 294 | 88 | 4 | 0.2475 | 0.3552 | |
| 4 and 5 | [1000,2000) | 495 | 139 | 1 | 0.2416 | 0.3226 | |
| 4 and 5 | | 495 | 111 | 2 | 0.1882 | 0.2636 | |
| 4 and 5 | | 495 | 115 | 3 | 0.1958 | 0.2721 | |
| 4 and 5 | | 495 | 130 | 4 | 0.2244 | 0.3037 | |
| 4 and 5 | [2000,3000) | 313 | 93 | 1 | 0.247 | 0.3511 | |
| 4 and 5 | | 313 | 74 | 2 | 0.1904 | 0.2875 | |
| 4 and 5 | | 313 | 60 | 3 | 0.1496 | 0.2397 | ✓ |
| 4 and 5 | | 313 | 86 | 4 | 0.226 | 0.3278 | |
| 4 and 5 | [3000,5000) | 665 | 167 | 1 | 0.2186 | 0.2859 | |
| 4 and 5 | | 665 | 184 | 2 | 0.243 | 0.3124 | |
| 4 and 5 | | 665 | 134 | 3 | 0.1716 | 0.234 | ✓ |
| 4 and 5 | | 665 | 180 | 4 | 0.2372 | 0.3062 | |
| 4 and 5 | [5000,10000) | 1503 | 381 | 1 | 0.2317 | 0.2763 | |
| 4 and 5 | | 1503 | 348 | 2 | 0.2104 | 0.2537 | |
| 4 and 5 | | 1503 | 371 | 3 | 0.2252 | 0.2695 | |
| 4 and 5 | | 1503 | 403 | 4 | 0.2459 | 0.2913 | |

| Pair | Distance interval (m) | Total | Count | Type | Lower CI | Upper CI | Sign. diff. |
|---|---|---|---|---|---|---|---|
| 5 and 8 | [0,50) | 36 | 4 | 1 | 0.0311 | 0.2606 | |
| 5 and 8 | | 36 | 0 | 2 | 0 | 0.0974 | ✓ |
| 5 and 8 | | 36 | 13 | 3 | 0.2082 | 0.5378 | |
| 5 and 8 | | 36 | 19 | 4 | 0.3549 | 0.6959 | ✓ |
| 5 and 8 | [50,100) | 35 | 3 | 1 | 0.018 | 0.2306 | ✓ |
| 5 and 8 | | 35 | 2 | 2 | 0.007 | 0.1916 | ✓ |
| 5 and 8 | | 35 | 23 | 3 | 0.4779 | 0.8087 | ✓ |
| 5 and 8 | | 35 | 7 | 4 | 0.0844 | 0.3694 | |
| 5 and 8 | [100,200) | 75 | 6 | 1 | 0.0299 | 0.166 | ✓ |
| 5 and 8 | | 75 | 5 | 2 | 0.022 | 0.1488 | ✓ |
| 5 and 8 | | 75 | 50 | 3 | 0.5483 | 0.7714 | ✓ |
| 5 and 8 | | 75 | 14 | 4 | 0.106 | 0.2933 | |
| 5 and 8 | [200,500) | 106 | 19 | 1 | 0.1115 | 0.2657 | |
| 5 and 8 | | 106 | 13 | 2 | 0.0669 | 0.2006 | ✓ |
| 5 and 8 | | 106 | 55 | 3 | 0.4197 | 0.617 | ✓ |
| 5 and 8 | | 106 | 19 | 4 | 0.1115 | 0.2657 | |
| 5 and 8 | [500,1000) | 124 | 25 | 1 | 0.1349 | 0.2831 | |
| 5 and 8 | | 124 | 22 | 2 | 0.1147 | 0.2562 | |
| 5 and 8 | | 124 | 56 | 3 | 0.3621 | 0.5435 | ✓ |
| 5 and 8 | | 124 | 21 | 4 | 0.108 | 0.2472 | ✓ |
| 5 and 8 | [1000,2000) | 258 | 61 | 1 | 0.1859 | 0.2931 | |
| 5 and 8 | | 258 | 54 | 2 | 0.1613 | 0.2641 | |
| 5 and 8 | | 258 | 85 | 3 | 0.2724 | 0.3905 | ✓ |
| 5 and 8 | | 258 | 58 | 4 | 0.1754 | 0.2807 | |
| 5 and 8 | [2000,3000) | 265 | 61 | 1 | 0.1809 | 0.2856 | |
| 5 and 8 | | 265 | 56 | 2 | 0.1638 | 0.2655 | |
| 5 and 8 | | 265 | 106 | 3 | 0.3405 | 0.4617 | ✓ |
| 5 and 8 | | 265 | 42 | 4 | 0.1167 | 0.2081 | ✓ |
| 5 and 8 | [3000,5000) | 599 | 156 | 1 | 0.2257 | 0.2975 | |
| 5 and 8 | | 599 | 171 | 2 | 0.2496 | 0.3235 | |
| 5 and 8 | | 599 | 156 | 3 | 0.2257 | 0.2975 | |
| 5 and 8 | | 599 | 116 | 4 | 0.1628 | 0.2276 | ✓ |
| 5 and 8 | [5000,10000) | 2344 | 565 | 1 | 0.2238 | 0.2589 | |
| 5 and 8 | | 2344 | 630 | 2 | 0.2509 | 0.2872 | ✓ |
| 5 and 8 | | 2344 | 540 | 3 | 0.2135 | 0.248 | ✓ |
| 5 and 8 | | 2344 | 609 | 4 | 0.2422 | 0.2781 | |

| Pair | Distance interval (m) | Total | Count | Type | Lower CI | Upper CI | Sign. diff. |
|---|---|---|---|---|---|---|---|
| 5 and 13 | [0,50) | 48 | 5 | 1 | 0.0347 | 0.2266 | ✓ |
| 5 and 13 | | 48 | 3 | 2 | 0.0131 | 0.172 | ✓ |
| 5 and 13 | | 48 | 28 | 3 | 0.4321 | 0.7239 | ✓ |
| 5 and 13 | | 48 | 12 | 4 | 0.1364 | 0.396 | |
| 5 and 13 | [50,100) | 53 | 10 | 1 | 0.0944 | 0.3197 | |
| 5 and 13 | | 53 | 8 | 2 | 0.0675 | 0.2759 | |
| 5 and 13 | | 53 | 23 | 3 | 0.2984 | 0.5772 | ✓ |
| 5 and 13 | | 53 | 12 | 4 | 0.1228 | 0.3621 | |
| 5 and 13 | [100,200) | 41 | 11 | 1 | 0.1422 | 0.4294 | |
| 5 and 13 | | 41 | 6 | 2 | 0.0557 | 0.2917 | |
| 5 and 13 | | 41 | 19 | 3 | 0.3066 | 0.6258 | ✓ |
| 5 and 13 | | 41 | 5 | 4 | 0.0408 | 0.262 | |
| 5 and 13 | [200,500) | 103 | 13 | 1 | 0.0689 | 0.2062 | ✓ |
| 5 and 13 | | 103 | 21 | 2 | 0.1309 | 0.2946 | |
| 5 and 13 | | 103 | 48 | 3 | 0.3671 | 0.567 | ✓ |
| 5 and 13 | | 103 | 21 | 4 | 0.1309 | 0.2946 | |
| 5 and 13 | [500,1000) | 124 | 31 | 1 | 0.1766 | 0.3357 | |
| 5 and 13 | | 124 | 26 | 2 | 0.1418 | 0.2919 | |
| 5 and 13 | | 124 | 52 | 3 | 0.3314 | 0.5113 | ✓ |
| 5 and 13 | | 124 | 15 | 4 | 0.0693 | 0.1917 | ✓ |
| 5 and 13 | [1000,2000) | 221 | 55 | 1 | 0.1933 | 0.3113 | |
| 5 and 13 | | 221 | 47 | 2 | 0.1606 | 0.2726 | |
| 5 and 13 | | 221 | 65 | 3 | 0.2349 | 0.3589 | |
| 5 and 13 | | 221 | 54 | 4 | 0.1892 | 0.3065 | |
| 5 and 13 | [2000,3000) | 209 | 53 | 1 | 0.1961 | 0.3182 | |
| 5 and 13 | | 209 | 56 | 2 | 0.2092 | 0.3334 | |
| 5 and 13 | | 209 | 41 | 3 | 0.1446 | 0.2566 | |
| 5 and 13 | | 209 | 59 | 4 | 0.2224 | 0.3485 | |
| 5 and 13 | [3000,5000) | 466 | 139 | 1 | 0.2571 | 0.3421 | ✓ |
| 5 and 13 | | 466 | 101 | 2 | 0.1802 | 0.257 | |
| 5 and 13 | | 466 | 112 | 3 | 0.2022 | 0.2818 | |
| 5 and 13 | | 466 | 114 | 4 | 0.2063 | 0.2863 | |
| 5 and 13 | [5000,10000) | 1442 | 343 | 1 | 0.2161 | 0.2607 | |
| 5 and 13 | | 1442 | 408 | 2 | 0.2598 | 0.307 | ✓ |
| 5 and 13 | | 1442 | 363 | 3 | 0.2295 | 0.275 | |
| 5 and 13 | | 1442 | 328 | 4 | 0.2061 | 0.25 | |

| Pair | Distance interval (m) | Total | Count | Type | Lower CI | Upper CI | Sign. diff. |
|---|---|---|---|---|---|---|---|
| 5 and 14 | [0,50) | 87 | 17 | 1 | 0.1181 | 0.2943 | |
| 5 and 14 | | 87 | 8 | 2 | 0.0405 | 0.1732 | ✓ |
| 5 and 14 | | 87 | 42 | 3 | 0.3742 | 0.5925 | ✓ |
| 5 and 14 | | 87 | 20 | 4 | 0.1464 | 0.3325 | |
| 5 and 14 | [50,100) | 36 | 7 | 1 | 0.0819 | 0.3602 | |
| 5 and 14 | | 36 | 3 | 2 | 0.0175 | 0.2247 | ✓ |
| 5 and 14 | | 36 | 17 | 3 | 0.3041 | 0.6451 | ✓ |
| 5 and 14 | | 36 | 9 | 4 | 0.1212 | 0.422 | |
| 5 and 14 | [100,200) | 49 | 4 | 1 | 0.0227 | 0.196 | ✓ |
| 5 and 14 | | 49 | 2 | 2 | 0.005 | 0.1398 | ✓ |
| 5 and 14 | | 49 | 34 | 3 | 0.5458 | 0.8175 | ✓ |
| 5 and 14 | | 49 | 9 | 4 | 0.0876 | 0.3202 | |
| 5 and 14 | [200,500) | 90 | 18 | 1 | 0.1231 | 0.2975 | |
| 5 and 14 | | 90 | 13 | 2 | 0.0792 | 0.2343 | ✓ |
| 5 and 14 | | 90 | 43 | 3 | 0.3713 | 0.5857 | ✓ |
| 5 and 14 | | 90 | 16 | 4 | 0.1052 | 0.2726 | |
| 5 and 14 | [500,1000) | 99 | 19 | 1 | 0.1197 | 0.2834 | |
| 5 and 14 | | 99 | 21 | 2 | 0.1364 | 0.3058 | |
| 5 and 14 | | 99 | 49 | 3 | 0.3929 | 0.5973 | ✓ |
| 5 and 14 | | 99 | 10 | 4 | 0.0495 | 0.1779 | ✓ |
| 5 and 14 | [1000,2000) | 278 | 58 | 1 | 0.1624 | 0.2612 | |
| 5 and 14 | | 278 | 59 | 2 | 0.1657 | 0.265 | |
| 5 and 14 | | 278 | 110 | 3 | 0.3378 | 0.4558 | ✓ |
| 5 and 14 | | 278 | 51 | 4 | 0.1398 | 0.234 | ✓ |
| 5 and 14 | [2000,3000) | 308 | 56 | 1 | 0.1404 | 0.2295 | ✓ |
| 5 and 14 | | 308 | 68 | 2 | 0.1757 | 0.2713 | |
| 5 and 14 | | 308 | 80 | 3 | 0.2117 | 0.3125 | |
| 5 and 14 | | 308 | 104 | 4 | 0.285 | 0.3935 | ✓ |
| 5 and 14 | [3000,5000) | 571 | 131 | 1 | 0.1955 | 0.2661 | |
| 5 and 14 | | 571 | 152 | 2 | 0.2304 | 0.3045 | |
| 5 and 14 | | 571 | 115 | 3 | 0.1692 | 0.2367 | ✓ |
| 5 and 14 | | 571 | 173 | 4 | 0.2655 | 0.3425 | ✓ |
| 5 and 14 | [5000,10000) | 1231 | 334 | 1 | 0.2466 | 0.2971 | |
| 5 and 14 | | 1231 | 311 | 2 | 0.2286 | 0.2779 | |
| 5 and 14 | | 1231 | 288 | 3 | 0.2106 | 0.2586 | |
| 5 and 14 | | 1231 | 298 | 4 | 0.2184 | 0.267 | |

| Pair | Distance interval (m) | Total | Count | Type | Lower CI | Upper CI | Sign. diff. |
|---|---|---|---|---|---|---|---|
| 5 and 15 | [0,50) | 95 | 10 | 1 | 0.0516 | 0.1851 | ✓ |
| 5 and 15 | | 95 | 8 | 2 | 0.0371 | 0.1592 | ✓ |
| 5 and 15 | | 95 | 63 | 3 | 0.5589 | 0.7569 | ✓ |
| 5 and 15 | | 95 | 14 | 4 | 0.083 | 0.2349 | ✓ |
| 5 and 15 | [50,100) | 63 | 4 | 1 | 0.0176 | 0.1547 | ✓ |
| 5 and 15 | | 63 | 11 | 2 | 0.0905 | 0.291 | |
| 5 and 15 | | 63 | 40 | 3 | 0.504 | 0.7527 | ✓ |
| 5 and 15 | | 63 | 8 | 4 | 0.0565 | 0.235 | ✓ |
| 5 and 15 | [100,200) | 94 | 4 | 1 | 0.0117 | 0.1054 | ✓ |
| 5 and 15 | | 94 | 11 | 2 | 0.0599 | 0.1997 | ✓ |
| 5 and 15 | | 94 | 64 | 3 | 0.5767 | 0.7733 | ✓ |
| 5 and 15 | | 94 | 15 | 4 | 0.0922 | 0.2495 | ✓ |
| 5 and 15 | [200,500) | 124 | 11 | 1 | 0.0451 | 0.1532 | ✓ |
| 5 and 15 | | 124 | 13 | 2 | 0.057 | 0.1726 | ✓ |
| 5 and 15 | | 124 | 87 | 3 | 0.6129 | 0.7804 | ✓ |
| 5 and 15 | | 124 | 13 | 4 | 0.057 | 0.1726 | ✓ |
| 5 and 15 | [500,1000) | 150 | 23 | 1 | 0.0998 | 0.2211 | ✓ |
| 5 and 15 | | 150 | 30 | 2 | 0.1392 | 0.273 | |
| 5 and 15 | | 150 | 70 | 3 | 0.3849 | 0.5498 | ✓ |
| 5 and 15 | | 150 | 27 | 4 | 0.1221 | 0.251 | |
| 5 and 15 | [1000,2000) | 178 | 27 | 1 | 0.1024 | 0.213 | ✓ |
| 5 and 15 | | 178 | 53 | 2 | 0.2317 | 0.3707 | |
| 5 and 15 | | 178 | 70 | 3 | 0.321 | 0.4691 | ✓ |
| 5 and 15 | | 178 | 28 | 4 | 0.1071 | 0.2193 | ✓ |
| 5 and 15 | [2000,3000) | 187 | 38 | 1 | 0.148 | 0.2681 | |
| 5 and 15 | | 187 | 44 | 2 | 0.1765 | 0.3027 | |
| 5 and 15 | | 187 | 51 | 3 | 0.2103 | 0.3425 | |
| 5 and 15 | | 187 | 54 | 4 | 0.225 | 0.3594 | |
| 5 and 15 | [3000,5000) | 386 | 97 | 1 | 0.2088 | 0.2977 | |
| 5 and 15 | | 386 | 80 | 2 | 0.1679 | 0.2512 | |
| 5 and 15 | | 386 | 116 | 3 | 0.2552 | 0.349 | ✓ |
| 5 and 15 | | 386 | 93 | 4 | 0.1991 | 0.2868 | |
| 5 and 15 | [5000,10000) | 1507 | 358 | 1 | 0.2163 | 0.2599 | |
| 5 and 15 | | 1507 | 358 | 2 | 0.2163 | 0.2599 | |
| 5 and 15 | | 1507 | 370 | 3 | 0.224 | 0.2681 | |
| 5 and 15 | | 1507 | 421 | 4 | 0.2568 | 0.3028 | ✓ |

| Pair | Distance interval (m) | Total | Count | Type | Lower CI | Upper CI | Sign. diff. |
|---|---|---|---|---|---|---|---|
| 6 and 8 | [0,50) | 316 | 52 | 1 | 0.1254 | 0.2101 | ✓ |
| 6 and 8 | | 316 | 35 | 2 | 0.0784 | 0.1507 | ✓ |
| 6 and 8 | | 316 | 111 | 3 | 0.2987 | 0.4067 | ✓ |
| 6 and 8 | | 316 | 118 | 4 | 0.3199 | 0.4293 | ✓ |
| 6 and 8 | [50,100) | 330 | 26 | 1 | 0.0521 | 0.1133 | ✓ |
| 6 and 8 | | 330 | 27 | 2 | 0.0546 | 0.1168 | ✓ |
| 6 and 8 | | 330 | 131 | 3 | 0.3438 | 0.452 | ✓ |
| 6 and 8 | | 330 | 146 | 4 | 0.3881 | 0.4978 | ✓ |
| 6 and 8 | [100,200) | 821 | 79 | 1 | 0.0769 | 0.1185 | ✓ |
| 6 and 8 | | 821 | 101 | 2 | 0.1013 | 0.1475 | ✓ |
| 6 and 8 | | 821 | 291 | 3 | 0.3217 | 0.3883 | ✓ |
| 6 and 8 | | 821 | 350 | 4 | 0.3922 | 0.461 | ✓ |
| 6 and 8 | [200,500) | 2326 | 291 | 1 | 0.1119 | 0.1392 | ✓ |
| 6 and 8 | | 2326 | 317 | 2 | 0.1226 | 0.1509 | ✓ |
| 6 and 8 | | 2326 | 769 | 3 | 0.3115 | 0.3501 | ✓ |
| 6 and 8 | | 2326 | 949 | 4 | 0.3879 | 0.4283 | ✓ |
| 6 and 8 | [500,1000) | 2508 | 417 | 1 | 0.1519 | 0.1814 | ✓ |
| 6 and 8 | | 2508 | 381 | 2 | 0.1381 | 0.1666 | ✓ |
| 6 and 8 | | 2508 | 831 | 3 | 0.3129 | 0.3501 | ✓ |
| 6 and 8 | | 2508 | 879 | 4 | 0.3318 | 0.3695 | ✓ |
| 6 and 8 | [1000,2000) | 2200 | 433 | 1 | 0.1804 | 0.2141 | ✓ |
| 6 and 8 | | 2200 | 439 | 2 | 0.183 | 0.2169 | ✓ |
| 6 and 8 | | 2200 | 722 | 3 | 0.3086 | 0.3482 | ✓ |
| 6 and 8 | | 2200 | 606 | 4 | 0.2569 | 0.2946 | ✓ |
| 6 and 8 | [2000,3000) | 1288 | 271 | 1 | 0.1884 | 0.2337 | ✓ |
| 6 and 8 | | 1288 | 275 | 2 | 0.1914 | 0.2369 | ✓ |
| 6 and 8 | | 1288 | 379 | 3 | 0.2695 | 0.32 | ✓ |
| 6 and 8 | | 1288 | 363 | 4 | 0.2574 | 0.3073 | ✓ |
| 6 and 8 | [3000,5000) | 1663 | 399 | 1 | 0.2196 | 0.2612 | |
| 6 and 8 | | 1663 | 378 | 2 | 0.2074 | 0.2482 | ✓ |
| 6 and 8 | | 1663 | 467 | 3 | 0.2593 | 0.3031 | ✓ |
| 6 and 8 | | 1663 | 419 | 4 | 0.2312 | 0.2735 | |
| 6 and 8 | [5000,10000) | 2706 | 687 | 1 | 0.2376 | 0.2707 | |
| 6 and 8 | | 2706 | 634 | 2 | 0.2184 | 0.2507 | |
| 6 and 8 | | 2706 | 705 | 3 | 0.2441 | 0.2775 | |
| 6 and 8 | | 2706 | 680 | 4 | 0.235 | 0.2681 | |

| Pair | Distance interval (m) | Total | Count | Type | Lower CI | Upper CI | Sign. diff. |
|---|---|---|---|---|---|---|---|
| 6 and 13 | [0,50) | 364 | 38 | 1 | 0.0749 | 0.1405 | ✓ |
| 6 and 13 | | 364 | 15 | 2 | 0.0232 | 0.0671 | ✓ |
| 6 and 13 | | 364 | 144 | 3 | 0.345 | 0.4479 | ✓ |
| 6 and 13 | | 364 | 167 | 4 | 0.4067 | 0.5115 | ✓ |
| 6 and 13 | [50,100) | 299 | 33 | 1 | 0.0772 | 0.1515 | ✓ |
| 6 and 13 | | 299 | 19 | 2 | 0.0387 | 0.0975 | ✓ |
| 6 and 13 | | 299 | 101 | 3 | 0.2844 | 0.3945 | ✓ |
| 6 and 13 | | 299 | 146 | 4 | 0.4303 | 0.5465 | ✓ |
| 6 and 13 | [100,200) | 557 | 71 | 1 | 0.1009 | 0.158 | ✓ |
| 6 and 13 | | 557 | 62 | 2 | 0.0864 | 0.1404 | ✓ |
| 6 and 13 | | 557 | 180 | 3 | 0.2844 | 0.3638 | ✓ |
| 6 and 13 | | 557 | 244 | 4 | 0.3964 | 0.4804 | ✓ |
| 6 and 13 | [200,500) | 1079 | 161 | 1 | 0.1285 | 0.1719 | ✓ |
| 6 and 13 | | 1079 | 121 | 2 | 0.0939 | 0.1325 | ✓ |
| 6 and 13 | | 1079 | 268 | 3 | 0.2228 | 0.2753 | |
| 6 and 13 | | 1079 | 529 | 4 | 0.46 | 0.5206 | ✓ |
| 6 and 13 | [500,1000) | 724 | 112 | 1 | 0.1291 | 0.1831 | ✓ |
| 6 and 13 | | 724 | 102 | 2 | 0.1164 | 0.1684 | ✓ |
| 6 and 13 | | 724 | 182 | 3 | 0.2202 | 0.2846 | |
| 6 and 13 | | 724 | 328 | 4 | 0.4163 | 0.4901 | ✓ |
| 6 and 13 | [1000,2000) | 629 | 122 | 1 | 0.1638 | 0.2271 | ✓ |
| 6 and 13 | | 629 | 114 | 2 | 0.1519 | 0.2136 | ✓ |
| 6 and 13 | | 629 | 159 | 3 | 0.2193 | 0.2887 | |
| 6 and 13 | | 629 | 234 | 4 | 0.3341 | 0.4111 | ✓ |
| 6 and 13 | [2000,3000) | 416 | 90 | 1 | 0.1777 | 0.2591 | |
| 6 and 13 | | 416 | 84 | 2 | 0.1644 | 0.2438 | ✓ |
| 6 and 13 | | 416 | 90 | 3 | 0.1777 | 0.2591 | |
| 6 and 13 | | 416 | 152 | 4 | 0.319 | 0.4137 | ✓ |
| 6 and 13 | [3000,5000) | 539 | 128 | 1 | 0.2021 | 0.2757 | |
| 6 and 13 | | 539 | 123 | 2 | 0.1934 | 0.266 | |
| 6 and 13 | | 539 | 108 | 3 | 0.1674 | 0.2367 | ✓ |
| 6 and 13 | | 539 | 180 | 4 | 0.2942 | 0.3755 | ✓ |
| 6 and 13 | [5000,10000) | 1179 | 279 | 1 | 0.2126 | 0.262 | |
| 6 and 13 | | 1179 | 315 | 2 | 0.2421 | 0.2934 | |
| 6 and 13 | | 1179 | 238 | 3 | 0.1793 | 0.2259 | ✓ |
| 6 and 13 | | 1179 | 347 | 4 | 0.2684 | 0.3212 | ✓ |

| Pair | Distance interval (m) | Total | Count | Type | Lower CI | Upper CI | Sign. diff. |
|---|---|---|---|---|---|---|---|
| 8 and 9 | [0,50) | 3 | 0 | 1 | 0 | 0.7076 | |
| 8 and 9 | | 3 | 1 | 2 | 0.0084 | 0.9057 | |
| 8 and 9 | | 3 | 2 | 3 | 0.0943 | 0.9916 | |
| 8 and 9 | | 3 | 0 | 4 | 0 | 0.7076 | |
| 8 and 9 | [50,100) | 5 | 1 | 1 | 0.0051 | 0.7164 | |
| 8 and 9 | | 5 | 1 | 2 | 0.0051 | 0.7164 | |
| 8 and 9 | | 5 | 1 | 3 | 0.0051 | 0.7164 | |
| 8 and 9 | | 5 | 2 | 4 | 0.0527 | 0.8534 | |
| 8 and 9 | [100,200) | 21 | 0 | 1 | 0 | 0.1611 | ✓ |
| 8 and 9 | | 21 | 6 | 2 | 0.1128 | 0.5218 | |
| 8 and 9 | | 21 | 5 | 3 | 0.0822 | 0.4717 | |
| 8 and 9 | | 21 | 10 | 4 | 0.2571 | 0.7022 | ✓ |
| 8 and 9 | [200,500) | 161 | 22 | 1 | 0.0877 | 0.1995 | ✓ |
| 8 and 9 | | 161 | 32 | 2 | 0.1401 | 0.2688 | |
| 8 and 9 | | 161 | 52 | 3 | 0.2515 | 0.4011 | ✓ |
| 8 and 9 | | 161 | 55 | 4 | 0.2688 | 0.4204 | ✓ |
| 8 and 9 | [500,1000) | 222 | 47 | 1 | 0.1599 | 0.2714 | |
| 8 and 9 | | 222 | 46 | 2 | 0.1559 | 0.2665 | |
| 8 and 9 | | 222 | 56 | 3 | 0.1965 | 0.3147 | |
| 8 and 9 | | 222 | 73 | 4 | 0.2674 | 0.3949 | ✓ |
| 8 and 9 | [1000,2000) | 278 | 59 | 1 | 0.1657 | 0.265 | |
| 8 and 9 | | 278 | 73 | 2 | 0.2118 | 0.3185 | |
| 8 and 9 | | 278 | 74 | 3 | 0.2152 | 0.3223 | |
| 8 and 9 | | 278 | 72 | 4 | 0.2085 | 0.3147 | |
| 8 and 9 | [2000,3000) | 187 | 35 | 1 | 0.134 | 0.2506 | |
| 8 and 9 | | 187 | 44 | 2 | 0.1765 | 0.3027 | |
| 8 and 9 | | 187 | 49 | 3 | 0.2006 | 0.3312 | |
| 8 and 9 | | 187 | 59 | 4 | 0.2496 | 0.3873 | |
| 8 and 9 | [3000,5000) | 288 | 76 | 1 | 0.2139 | 0.3188 | |
| 8 and 9 | | 288 | 44 | 2 | 0.1133 | 0.1996 | ✓ |
| 8 and 9 | | 288 | 59 | 3 | 0.1598 | 0.2561 | |
| 8 and 9 | | 288 | 109 | 4 | 0.3222 | 0.4372 | ✓ |
| 8 and 9 | [5000,10000) | 299 | 83 | 1 | 0.2276 | 0.3321 | |
| 8 and 9 | | 299 | 87 | 2 | 0.2401 | 0.346 | |
| 8 and 9 | | 299 | 60 | 3 | 0.1568 | 0.2506 | |
| 8 and 9 | | 299 | 69 | 4 | 0.1842 | 0.2827 | |

| Pair | Distance interval (m) | Total | Count | Type | Lower CI | Upper CI | Sign. diff. |
|---|---|---|---|---|---|---|---|
| 8 and 10 | [0,50) | 14 | 1 | 1 | 0.0018 | 0.3387 | |
| 8 and 10 | | 14 | 2 | 2 | 0.0178 | 0.4281 | |
| 8 and 10 | | 14 | 1 | 3 | 0.0018 | 0.3387 | |
| 8 and 10 | | 14 | 10 | 4 | 0.419 | 0.9161 | ✓ |
| 8 and 10 | [50,100) | 10 | 1 | 1 | 0.0025 | 0.445 | |
| 8 and 10 | | 10 | 0 | 2 | 0 | 0.3085 | |
| 8 and 10 | | 10 | 5 | 3 | 0.1871 | 0.8129 | |
| 8 and 10 | | 10 | 4 | 4 | 0.1216 | 0.7376 | |
| 8 and 10 | [100,200) | 40 | 6 | 1 | 0.0571 | 0.2984 | |
| 8 and 10 | | 40 | 7 | 2 | 0.0734 | 0.3278 | |
| 8 and 10 | | 40 | 11 | 3 | 0.146 | 0.4389 | |
| 8 and 10 | | 40 | 16 | 4 | 0.2486 | 0.5667 | |
| 8 and 10 | [200,500) | 188 | 35 | 1 | 0.1332 | 0.2493 | ✓ |
| 8 and 10 | | 188 | 30 | 2 | 0.1103 | 0.2199 | ✓ |
| 8 and 10 | | 188 | 73 | 3 | 0.3182 | 0.4619 | ✓ |
| 8 and 10 | | 188 | 50 | 4 | 0.2043 | 0.3352 | |
| 8 and 10 | [500,1000) | 254 | 49 | 1 | 0.1463 | 0.2469 | ✓ |
| 8 and 10 | | 254 | 49 | 2 | 0.1463 | 0.2469 | ✓ |
| 8 and 10 | | 254 | 74 | 3 | 0.2362 | 0.3514 | |
| 8 and 10 | | 254 | 82 | 4 | 0.2657 | 0.3841 | ✓ |
| 8 and 10 | [1000,2000) | 295 | 57 | 1 | 0.1498 | 0.243 | ✓ |
| 8 and 10 | | 295 | 57 | 2 | 0.1498 | 0.243 | ✓ |
| 8 and 10 | | 295 | 96 | 3 | 0.2723 | 0.3821 | ✓ |
| 8 and 10 | | 295 | 85 | 4 | 0.2371 | 0.3435 | |
| 8 and 10 | [2000,3000) | 233 | 55 | 1 | 0.1831 | 0.2959 | |
| 8 and 10 | | 233 | 48 | 2 | 0.156 | 0.2637 | |
| 8 and 10 | | 233 | 64 | 3 | 0.2184 | 0.3368 | |
| 8 and 10 | | 233 | 66 | 4 | 0.2264 | 0.3458 | |
| 8 and 10 | [3000,5000) | 327 | 85 | 1 | 0.2132 | 0.311 | |
| 8 and 10 | | 327 | 77 | 2 | 0.1905 | 0.2853 | |
| 8 and 10 | | 327 | 57 | 3 | 0.1348 | 0.2199 | ✓ |
| 8 and 10 | | 327 | 108 | 4 | 0.2795 | 0.3841 | ✓ |
| 8 and 10 | [5000,10000) | 469 | 118 | 1 | 0.2129 | 0.2934 | |
| 8 and 10 | | 469 | 149 | 2 | 0.2758 | 0.362 | ✓ |
| 8 and 10 | | 469 | 111 | 3 | 0.1989 | 0.2778 | |
| 8 and 10 | | 469 | 91 | 4 | 0.1592 | 0.2328 | ✓ |

| Pair | Distance interval (m) | Total | Count | Type | Lower CI | Upper CI | Sign. diff. |
|---|---|---|---|---|---|---|---|
| 8 and 13 | [0,50) | 235 | 26 | 1 | 0.0736 | 0.1579 | ✓ |
| 8 and 13 | | 235 | 9 | 2 | 0.0177 | 0.0715 | ✓ |
| 8 and 13 | | 235 | 95 | 3 | 0.3409 | 0.47 | ✓ |
| 8 and 13 | | 235 | 105 | 4 | 0.3821 | 0.5128 | ✓ |
| 8 and 13 | [50,100) | 200 | 23 | 1 | 0.0743 | 0.1675 | ✓ |
| 8 and 13 | | 200 | 7 | 2 | 0.0142 | 0.0708 | ✓ |
| 8 and 13 | | 200 | 77 | 3 | 0.3172 | 0.4562 | ✓ |
| 8 and 13 | | 200 | 93 | 4 | 0.3944 | 0.5367 | ✓ |
| 8 and 13 | [100,200) | 317 | 28 | 1 | 0.0595 | 0.1251 | ✓ |
| 8 and 13 | | 317 | 26 | 2 | 0.0543 | 0.1179 | ✓ |
| 8 and 13 | | 317 | 105 | 3 | 0.2796 | 0.386 | ✓ |
| 8 and 13 | | 317 | 158 | 4 | 0.442 | 0.5548 | ✓ |
| 8 and 13 | [200,500) | 673 | 88 | 1 | 0.1062 | 0.1586 | ✓ |
| 8 and 13 | | 673 | 71 | 2 | 0.0833 | 0.1312 | ✓ |
| 8 and 13 | | 673 | 220 | 3 | 0.2915 | 0.3638 | ✓ |
| 8 and 13 | | 673 | 294 | 4 | 0.399 | 0.4753 | ✓ |
| 8 and 13 | [500,1000) | 689 | 106 | 1 | 0.1277 | 0.183 | ✓ |
| 8 and 13 | | 689 | 111 | 2 | 0.1344 | 0.1907 | ✓ |
| 8 and 13 | | 689 | 186 | 3 | 0.2371 | 0.3048 | |
| 8 and 13 | | 689 | 286 | 4 | 0.378 | 0.4529 | ✓ |
| 8 and 13 | [1000,2000) | 514 | 95 | 1 | 0.1522 | 0.2211 | ✓ |
| 8 and 13 | | 514 | 96 | 2 | 0.154 | 0.2232 | ✓ |
| 8 and 13 | | 514 | 104 | 3 | 0.1684 | 0.2397 | ✓ |
| 8 and 13 | | 514 | 219 | 4 | 0.3829 | 0.4701 | ✓ |
| 8 and 13 | [2000,3000) | 327 | 81 | 1 | 0.2019 | 0.2982 | |
| 8 and 13 | | 327 | 72 | 2 | 0.1765 | 0.2691 | |
| 8 and 13 | | 327 | 53 | 3 | 0.1238 | 0.2066 | ✓ |
| 8 and 13 | | 327 | 121 | 4 | 0.3176 | 0.4249 | ✓ |
| 8 and 13 | [3000,5000) | 409 | 91 | 1 | 0.1831 | 0.266 | |
| 8 and 13 | | 409 | 96 | 2 | 0.1945 | 0.2789 | |
| 8 and 13 | | 409 | 84 | 3 | 0.1673 | 0.2478 | ✓ |
| 8 and 13 | | 409 | 138 | 4 | 0.2917 | 0.3855 | ✓ |
| 8 and 13 | [5000,10000) | 827 | 184 | 1 | 0.1946 | 0.2524 | |
| 8 and 13 | | 827 | 191 | 2 | 0.2026 | 0.2612 | |
| 8 and 13 | | 827 | 204 | 3 | 0.2176 | 0.2775 | |
| 8 and 13 | | 827 | 248 | 4 | 0.2688 | 0.3324 | ✓ |

| Pair | Distance interval (m) | Total | Count | Type | Lower CI | Upper CI | Sign. diff. |
|---|---|---|---|---|---|---|---|
| 9 and 10 | [0,50) | 4671 | 261 | 1 | 0.0495 | 0.0629 | ✓ |
| 9 and 10 | | 4671 | 187 | 2 | 0.0346 | 0.0461 | ✓ |
| 9 and 10 | | 4671 | 2077 | 3 | 0.4303 | 0.459 | ✓ |
| 9 and 10 | | 4671 | 2146 | 4 | 0.4451 | 0.4739 | ✓ |
| 9 and 10 | [50,100) | 5707 | 264 | 1 | 0.041 | 0.052 | ✓ |
| 9 and 10 | | 5707 | 282 | 2 | 0.0439 | 0.0554 | ✓ |
| 9 and 10 | | 5707 | 2741 | 3 | 0.4673 | 0.4933 | ✓ |
| 9 and 10 | | 5707 | 2420 | 4 | 0.4112 | 0.437 | ✓ |
| 9 and 10 | [100,200) | 9126 | 609 | 1 | 0.0617 | 0.072 | ✓ |
| 9 and 10 | | 9126 | 674 | 2 | 0.0686 | 0.0794 | ✓ |
| 9 and 10 | | 9126 | 4394 | 3 | 0.4712 | 0.4918 | ✓ |
| 9 and 10 | | 9126 | 3449 | 4 | 0.368 | 0.388 | ✓ |
| 9 and 10 | [200,500) | 14708 | 1744 | 1 | 0.1134 | 0.1239 | ✓ |
| 9 and 10 | | 14708 | 1771 | 2 | 0.1152 | 0.1258 | ✓ |
| 9 and 10 | | 14708 | 6407 | 3 | 0.4276 | 0.4437 | ✓ |
| 9 and 10 | | 14708 | 4786 | 4 | 0.3178 | 0.333 | ✓ |
| 9 and 10 | [500,1000) | 8737 | 1490 | 1 | 0.1627 | 0.1786 | ✓ |
| 9 and 10 | | 8737 | 1407 | 2 | 0.1534 | 0.1689 | ✓ |
| 9 and 10 | | 8737 | 3270 | 3 | 0.3641 | 0.3845 | ✓ |
| 9 and 10 | | 8737 | 2570 | 4 | 0.2846 | 0.3038 | ✓ |
| 9 and 10 | [1000,2000) | 4234 | 835 | 1 | 0.1853 | 0.2095 | ✓ |
| 9 and 10 | | 4234 | 872 | 2 | 0.1939 | 0.2185 | ✓ |
| 9 and 10 | | 4234 | 1396 | 3 | 0.3156 | 0.3441 | ✓ |
| 9 and 10 | | 4234 | 1131 | 4 | 0.2538 | 0.2807 | ✓ |
| 9 and 10 | [2000,3000) | 1342 | 294 | 1 | 0.1972 | 0.2422 | ✓ |
| 9 and 10 | | 1342 | 284 | 2 | 0.19 | 0.2345 | ✓ |
| 9 and 10 | | 1342 | 398 | 3 | 0.2722 | 0.3218 | ✓ |
| 9 and 10 | | 1342 | 366 | 4 | 0.249 | 0.2974 | |
| 9 and 10 | [3000,5000) | 1602 | 380 | 1 | 0.2166 | 0.2588 | |
| 9 and 10 | | 1602 | 351 | 2 | 0.1991 | 0.2402 | ✓ |
| 9 and 10 | | 1602 | 415 | 3 | 0.2377 | 0.2812 | |
| 9 and 10 | | 1602 | 456 | 4 | 0.2626 | 0.3074 | ✓ |
| 9 and 10 | [5000,10000) | 2258 | 612 | 1 | 0.2528 | 0.2899 | ✓ |
| 9 and 10 | | 2258 | 564 | 2 | 0.232 | 0.2682 | |
| 9 and 10 | | 2258 | 540 | 3 | 0.2217 | 0.2573 | |
| 9 and 10 | | 2258 | 542 | 4 | 0.2225 | 0.2582 | |

| Pair | Distance interval (m) | Total | Count | Type | Lower CI | Upper CI | Sign. diff. |
|---|---|---|---|---|---|---|---|
| 9 and 11 | [0,50) | 10 | 0 | 1 | 0 | 0.3085 | |
| 9 and 11 | | 10 | 2 | 2 | 0.0252 | 0.5561 | |
| 9 and 11 | | 10 | 5 | 3 | 0.1871 | 0.8129 | |
| 9 and 11 | | 10 | 3 | 4 | 0.0667 | 0.6525 | |
| 9 and 11 | [50,100) | 12 | 0 | 1 | 0 | 0.2646 | |
| 9 and 11 | | 12 | 3 | 2 | 0.0549 | 0.5719 | |
| 9 and 11 | | 12 | 4 | 3 | 0.0992 | 0.6511 | |
| 9 and 11 | | 12 | 5 | 4 | 0.1517 | 0.7233 | |
| 9 and 11 | [100,200) | 41 | 3 | 1 | 0.0154 | 0.1992 | ✓ |
| 9 and 11 | | 41 | 5 | 2 | 0.0408 | 0.262 | |
| 9 and 11 | | 41 | 19 | 3 | 0.3066 | 0.6258 | ✓ |
| 9 and 11 | | 41 | 14 | 4 | 0.2008 | 0.5059 | |
| 9 and 11 | [200,500) | 158 | 27 | 1 | 0.1157 | 0.2388 | ✓ |
| 9 and 11 | | 158 | 34 | 2 | 0.1539 | 0.2875 | |
| 9 and 11 | | 158 | 54 | 3 | 0.2683 | 0.4213 | ✓ |
| 9 and 11 | | 158 | 43 | 4 | 0.2045 | 0.3486 | |
| 9 and 11 | [500,1000) | 218 | 46 | 1 | 0.1588 | 0.2712 | |
| 9 and 11 | | 218 | 47 | 2 | 0.1629 | 0.2762 | |
| 9 and 11 | | 218 | 59 | 3 | 0.2129 | 0.3348 | |
| 9 and 11 | | 218 | 66 | 4 | 0.2425 | 0.3684 | |
| 9 and 11 | [1000,2000) | 422 | 84 | 1 | 0.162 | 0.2404 | ✓ |
| 9 and 11 | | 422 | 107 | 2 | 0.2127 | 0.2979 | |
| 9 and 11 | | 422 | 122 | 3 | 0.2463 | 0.3349 | |
| 9 and 11 | | 422 | 109 | 4 | 0.2172 | 0.3028 | |
| 9 and 11 | [2000,3000) | 366 | 90 | 1 | 0.2026 | 0.2933 | |
| 9 and 11 | | 366 | 93 | 2 | 0.2103 | 0.3019 | |
| 9 and 11 | | 366 | 71 | 3 | 0.1547 | 0.2383 | ✓ |
| 9 and 11 | | 366 | 112 | 4 | 0.2592 | 0.356 | ✓ |
| 9 and 11 | [3000,5000) | 780 | 187 | 1 | 0.2102 | 0.2713 | |
| 9 and 11 | | 780 | 207 | 2 | 0.2347 | 0.2979 | |
| 9 and 11 | | 780 | 185 | 3 | 0.2077 | 0.2686 | |
| 9 and 11 | | 780 | 201 | 4 | 0.2273 | 0.2899 | |
| 9 and 11 | [5000,10000) | 2520 | 561 | 1 | 0.2065 | 0.2394 | ✓ |
| 9 and 11 | | 2520 | 710 | 2 | 0.2642 | 0.2998 | ✓ |
| 9 and 11 | | 2520 | 581 | 3 | 0.2142 | 0.2475 | ✓ |
| 9 and 11 | | 2520 | 668 | 4 | 0.2479 | 0.2828 | |

| Pair | Distance interval (m) | Total | Count | Type | Lower CI | Upper CI | Sign. diff. |
|---|---|---|---|---|---|---|---|
| 9 and 13 | [0,50) | 485 | 49 | 1 | 0.0757 | 0.1314 | ✓ |
| 9 and 13 | | 485 | 26 | 2 | 0.0353 | 0.0776 | ✓ |
| 9 and 13 | | 485 | 185 | 3 | 0.338 | 0.4263 | ✓ |
| 9 and 13 | | 485 | 225 | 4 | 0.4188 | 0.5094 | ✓ |
| 9 and 13 | [50,100) | 522 | 27 | 1 | 0.0344 | 0.0744 | ✓ |
| 9 and 13 | | 522 | 41 | 2 | 0.057 | 0.105 | ✓ |
| 9 and 13 | | 522 | 207 | 3 | 0.3543 | 0.44 | ✓ |
| 9 and 13 | | 522 | 247 | 4 | 0.4296 | 0.517 | ✓ |
| 9 and 13 | [100,200) | 857 | 80 | 1 | 0.0747 | 0.1148 | ✓ |
| 9 and 13 | | 857 | 78 | 2 | 0.0726 | 0.1123 | ✓ |
| 9 and 13 | | 857 | 328 | 3 | 0.3501 | 0.4162 | ✓ |
| 9 and 13 | | 857 | 371 | 4 | 0.3994 | 0.4668 | ✓ |
| 9 and 13 | [200,500) | 1700 | 230 | 1 | 0.1194 | 0.1525 | ✓ |
| 9 and 13 | | 1700 | 208 | 2 | 0.1071 | 0.1389 | ✓ |
| 9 and 13 | | 1700 | 527 | 3 | 0.2881 | 0.3326 | ✓ |
| 9 and 13 | | 1700 | 735 | 4 | 0.4086 | 0.4563 | ✓ |
| 9 and 13 | [500,1000) | 1369 | 222 | 1 | 0.143 | 0.1828 | ✓ |
| 9 and 13 | | 1369 | 214 | 2 | 0.1375 | 0.1767 | ✓ |
| 9 and 13 | | 1369 | 276 | 3 | 0.1806 | 0.2239 | ✓ |
| 9 and 13 | | 1369 | 657 | 4 | 0.4531 | 0.5068 | ✓ |
| 9 and 13 | [1000,2000) | 1064 | 211 | 1 | 0.1747 | 0.2236 | ✓ |
| 9 and 13 | | 1064 | 187 | 2 | 0.1533 | 0.2 | ✓ |
| 9 and 13 | | 1064 | 176 | 3 | 0.1436 | 0.1891 | ✓ |
| 9 and 13 | | 1064 | 490 | 4 | 0.4303 | 0.491 | ✓ |
| 9 and 13 | [2000,3000) | 742 | 143 | 1 | 0.1649 | 0.223 | ✓ |
| 9 and 13 | | 742 | 170 | 2 | 0.1993 | 0.2611 | |
| 9 and 13 | | 742 | 153 | 3 | 0.1776 | 0.2371 | ✓ |
| 9 and 13 | | 742 | 276 | 4 | 0.3371 | 0.4079 | ✓ |
| 9 and 13 | [3000,5000) | 997 | 216 | 1 | 0.1914 | 0.2435 | ✓ |
| 9 and 13 | | 997 | 232 | 2 | 0.2068 | 0.2602 | |
| 9 and 13 | | 997 | 280 | 3 | 0.2531 | 0.3099 | ✓ |
| 9 and 13 | | 997 | 269 | 4 | 0.2425 | 0.2985 | |
| 9 and 13 | [5000,10000) | 2144 | 559 | 1 | 0.2422 | 0.2799 | |
| 9 and 13 | | 2144 | 512 | 2 | 0.2209 | 0.2574 | |
| 9 and 13 | | 2144 | 517 | 3 | 0.2232 | 0.2598 | |
| 9 and 13 | | 2144 | 556 | 4 | 0.2409 | 0.2784 | |

| Pair | Distance interval (m) | Total | Count | Type | Lower CI | Upper CI | Sign. diff. |
|---|---|---|---|---|---|---|---|
| 10 and 11 | [0,50) | 14 | 1 | 1 | 0.0018 | 0.3387 | |
| 10 and 11 | | 14 | 0 | 2 | 0 | 0.2316 | ✓ |
| 10 and 11 | | 14 | 6 | 3 | 0.1766 | 0.7114 | |
| 10 and 11 | | 14 | 7 | 4 | 0.2304 | 0.7696 | |
| 10 and 11 | [50,100) | 25 | 1 | 1 | 0.001 | 0.2035 | ✓ |
| 10 and 11 | | 25 | 3 | 2 | 0.0255 | 0.3122 | |
| 10 and 11 | | 25 | 14 | 3 | 0.3493 | 0.756 | ✓ |
| 10 and 11 | | 25 | 7 | 4 | 0.1207 | 0.4939 | |
| 10 and 11 | [100,200) | 60 | 8 | 1 | 0.0594 | 0.2459 | ✓ |
| 10 and 11 | | 60 | 10 | 2 | 0.0829 | 0.2852 | |
| 10 and 11 | | 60 | 30 | 3 | 0.3681 | 0.6319 | ✓ |
| 10 and 11 | | 60 | 12 | 4 | 0.1078 | 0.3233 | |
| 10 and 11 | [200,500) | 152 | 27 | 1 | 0.1204 | 0.2478 | ✓ |
| 10 and 11 | | 152 | 31 | 2 | 0.143 | 0.2768 | |
| 10 and 11 | | 152 | 41 | 3 | 0.201 | 0.3476 | |
| 10 and 11 | | 152 | 53 | 4 | 0.2733 | 0.4301 | ✓ |
| 10 and 11 | [500,1000) | 206 | 38 | 1 | 0.134 | 0.2443 | ✓ |
| 10 and 11 | | 206 | 50 | 2 | 0.1858 | 0.3072 | |
| 10 and 11 | | 206 | 42 | 3 | 0.1511 | 0.2654 | |
| 10 and 11 | | 206 | 76 | 4 | 0.3029 | 0.4388 | ✓ |
| 10 and 11 | [1000,2000) | 384 | 80 | 1 | 0.1688 | 0.2524 | |
| 10 and 11 | | 384 | 111 | 2 | 0.2442 | 0.3372 | |
| 10 and 11 | | 384 | 83 | 3 | 0.176 | 0.2607 | |
| 10 and 11 | | 384 | 110 | 4 | 0.2417 | 0.3345 | |
| 10 and 11 | [2000,3000) | 453 | 93 | 1 | 0.169 | 0.2455 | ✓ |
| 10 and 11 | | 453 | 107 | 2 | 0.1978 | 0.2781 | |
| 10 and 11 | | 453 | 96 | 3 | 0.1752 | 0.2525 | |
| 10 and 11 | | 453 | 157 | 4 | 0.3028 | 0.3924 | ✓ |
| 10 and 11 | [3000,5000) | 936 | 210 | 1 | 0.198 | 0.2525 | |
| 10 and 11 | | 936 | 249 | 2 | 0.238 | 0.2956 | |
| 10 and 11 | | 936 | 234 | 3 | 0.2225 | 0.279 | |
| 10 and 11 | | 936 | 243 | 4 | 0.2318 | 0.289 | |
| 10 and 11 | [5000,10000) | 2545 | 549 | 1 | 0.1999 | 0.2322 | ✓ |
| 10 and 11 | | 2545 | 742 | 2 | 0.2739 | 0.3096 | ✓ |
| 10 and 11 | | 2545 | 587 | 3 | 0.2144 | 0.2475 | ✓ |
| 10 and 11 | | 2545 | 667 | 4 | 0.2451 | 0.2796 | |

| Pair | Distance interval (m) | Total | Count | Type | Lower CI | Upper CI | Sign. diff. |
|---|---|---|---|---|---|---|---|
| 10 and 12 | [0,50) | 9 | 0 | 1 | 0 | 0.3363 | |
| 10 and 12 | | 9 | 2 | 2 | 0.0281 | 0.6001 | |
| 10 and 12 | | 9 | 3 | 3 | 0.0749 | 0.7007 | |
| 10 and 12 | | 9 | 4 | 4 | 0.137 | 0.788 | |
| 10 and 12 | [50,100) | 21 | 2 | 1 | 0.0117 | 0.3038 | |
| 10 and 12 | | 21 | 2 | 2 | 0.0117 | 0.3038 | |
| 10 and 12 | | 21 | 10 | 3 | 0.2571 | 0.7022 | ✓ |
| 10 and 12 | | 21 | 7 | 4 | 0.1459 | 0.5697 | |
| 10 and 12 | [100,200) | 46 | 1 | 1 | 6e-04 | 0.1153 | ✓ |
| 10 and 12 | | 46 | 5 | 2 | 0.0362 | 0.2357 | ✓ |
| 10 and 12 | | 46 | 24 | 3 | 0.3695 | 0.6711 | ✓ |
| 10 and 12 | | 46 | 16 | 4 | 0.2135 | 0.5025 | |
| 10 and 12 | [200,500) | 133 | 17 | 1 | 0.0763 | 0.1967 | ✓ |
| 10 and 12 | | 133 | 19 | 2 | 0.0883 | 0.2141 | ✓ |
| 10 and 12 | | 133 | 47 | 3 | 0.2725 | 0.4409 | ✓ |
| 10 and 12 | | 133 | 50 | 4 | 0.2935 | 0.464 | ✓ |
| 10 and 12 | [500,1000) | 124 | 19 | 1 | 0.0948 | 0.2289 | ✓ |
| 10 and 12 | | 124 | 35 | 2 | 0.2051 | 0.3701 | |
| 10 and 12 | | 124 | 36 | 3 | 0.2123 | 0.3786 | |
| 10 and 12 | | 124 | 34 | 4 | 0.1979 | 0.3615 | |
| 10 and 12 | [1000,2000) | 210 | 29 | 1 | 0.0945 | 0.1923 | ✓ |
| 10 and 12 | | 210 | 59 | 2 | 0.2213 | 0.3469 | |
| 10 and 12 | | 210 | 68 | 3 | 0.261 | 0.3916 | ✓ |
| 10 and 12 | | 210 | 54 | 4 | 0.1995 | 0.3218 | |
| 10 and 12 | [2000,3000) | 188 | 35 | 1 | 0.1332 | 0.2493 | ✓ |
| 10 and 12 | | 188 | 48 | 2 | 0.1946 | 0.3239 | |
| 10 and 12 | | 188 | 61 | 3 | 0.2581 | 0.3964 | ✓ |
| 10 and 12 | | 188 | 44 | 4 | 0.1755 | 0.3012 | |
| 10 and 12 | [3000,5000) | 351 | 67 | 1 | 0.1511 | 0.236 | ✓ |
| 10 and 12 | | 351 | 85 | 2 | 0.1983 | 0.2905 | |
| 10 and 12 | | 351 | 98 | 3 | 0.2329 | 0.3293 | |
| 10 and 12 | | 351 | 101 | 4 | 0.2409 | 0.3382 | |
| 10 and 12 | [5000,10000) | 416 | 106 | 1 | 0.2136 | 0.2995 | |
| 10 and 12 | | 416 | 118 | 2 | 0.2408 | 0.3296 | |
| 10 and 12 | | 416 | 106 | 3 | 0.2136 | 0.2995 | |
| 10 and 12 | | 416 | 86 | 4 | 0.1688 | 0.2489 | ✓ |

| Pair | Distance interval (m) | Total | Count | Type | Lower CI | Upper CI | Sign. diff. |
|---|---|---|---|---|---|---|---|
| 10 and 13 | [0,50) | 613 | 48 | 1 | 0.0583 | 0.1025 | ✓ |
| 10 and 13 | | 613 | 31 | 2 | 0.0346 | 0.071 | ✓ |
| 10 and 13 | | 613 | 278 | 3 | 0.4136 | 0.4939 | ✓ |
| 10 and 13 | | 613 | 256 | 4 | 0.3782 | 0.4578 | ✓ |
| 10 and 13 | [50,100) | 550 | 28 | 1 | 0.0341 | 0.0727 | ✓ |
| 10 and 13 | | 550 | 32 | 2 | 0.0401 | 0.0811 | ✓ |
| 10 and 13 | | 550 | 245 | 3 | 0.4034 | 0.4881 | ✓ |
| 10 and 13 | | 550 | 245 | 4 | 0.4034 | 0.4881 | ✓ |
| 10 and 13 | [100,200) | 952 | 78 | 1 | 0.0653 | 0.1012 | ✓ |
| 10 and 13 | | 952 | 81 | 2 | 0.0681 | 0.1046 | ✓ |
| 10 and 13 | | 952 | 368 | 3 | 0.3555 | 0.4183 | ✓ |
| 10 and 13 | | 952 | 425 | 4 | 0.4145 | 0.4787 | ✓ |
| 10 and 13 | [200,500) | 1827 | 218 | 1 | 0.1048 | 0.1351 | ✓ |
| 10 and 13 | | 1827 | 231 | 2 | 0.1115 | 0.1426 | ✓ |
| 10 and 13 | | 1827 | 497 | 3 | 0.2517 | 0.2931 | ✓ |
| 10 and 13 | | 1827 | 881 | 4 | 0.4591 | 0.5054 | ✓ |
| 10 and 13 | [500,1000) | 1645 | 239 | 1 | 0.1286 | 0.1633 | ✓ |
| 10 and 13 | | 1645 | 259 | 2 | 0.1402 | 0.176 | ✓ |
| 10 and 13 | | 1645 | 399 | 3 | 0.222 | 0.264 | |
| 10 and 13 | | 1645 | 748 | 4 | 0.4304 | 0.4791 | ✓ |
| 10 and 13 | [1000,2000) | 1417 | 278 | 1 | 0.1758 | 0.2178 | ✓ |
| 10 and 13 | | 1417 | 269 | 2 | 0.1697 | 0.2112 | ✓ |
| 10 and 13 | | 1417 | 269 | 3 | 0.1697 | 0.2112 | ✓ |
| 10 and 13 | | 1417 | 601 | 4 | 0.3982 | 0.4504 | ✓ |
| 10 and 13 | [2000,3000) | 980 | 174 | 1 | 0.1541 | 0.2029 | ✓ |
| 10 and 13 | | 980 | 213 | 2 | 0.1919 | 0.2445 | ✓ |
| 10 and 13 | | 980 | 231 | 3 | 0.2095 | 0.2636 | |
| 10 and 13 | | 980 | 362 | 4 | 0.3391 | 0.4005 | ✓ |
| 10 and 13 | [3000,5000) | 1358 | 285 | 1 | 0.1885 | 0.2325 | ✓ |
| 10 and 13 | | 1358 | 335 | 2 | 0.224 | 0.2705 | |
| 10 and 13 | | 1358 | 380 | 3 | 0.2561 | 0.3045 | ✓ |
| 10 and 13 | | 1358 | 358 | 4 | 0.2404 | 0.2879 | |
| 10 and 13 | [5000,10000) | 2605 | 680 | 1 | 0.2443 | 0.2784 | |
| 10 and 13 | | 2605 | 619 | 2 | 0.2214 | 0.2544 | |
| 10 and 13 | | 2605 | 662 | 3 | 0.2375 | 0.2713 | |
| 10 and 13 | | 2605 | 644 | 4 | 0.2308 | 0.2643 | |

| Pair | Distance interval (m) | Total | Count | Type | Lower CI | Upper CI | Sign. diff. |
|---|---|---|---|---|---|---|---|
| 13 and 15 | [0,50) | 289 | 32 | 1 | 0.077 | 0.1527 | ✓ |
| 13 and 15 | | 289 | 24 | 2 | 0.0539 | 0.121 | ✓ |
| 13 and 15 | | 289 | 101 | 3 | 0.2946 | 0.4075 | ✓ |
| 13 and 15 | | 289 | 132 | 4 | 0.3983 | 0.5161 | ✓ |
| 13 and 15 | [50,100) | 202 | 19 | 1 | 0.0576 | 0.143 | ✓ |
| 13 and 15 | | 202 | 14 | 2 | 0.0384 | 0.1136 | ✓ |
| 13 and 15 | | 202 | 81 | 3 | 0.3328 | 0.4721 | ✓ |
| 13 and 15 | | 202 | 88 | 4 | 0.3662 | 0.507 | ✓ |
| 13 and 15 | [100,200) | 228 | 25 | 1 | 0.0722 | 0.1576 | ✓ |
| 13 and 15 | | 228 | 29 | 2 | 0.0869 | 0.1775 | ✓ |
| 13 and 15 | | 228 | 75 | 3 | 0.2684 | 0.3941 | ✓ |
| 13 and 15 | | 228 | 99 | 4 | 0.3689 | 0.5012 | ✓ |
| 13 and 15 | [200,500) | 364 | 63 | 1 | 0.1356 | 0.2159 | ✓ |
| 13 and 15 | | 364 | 65 | 2 | 0.1406 | 0.2219 | ✓ |
| 13 and 15 | | 364 | 151 | 3 | 0.3637 | 0.4673 | ✓ |
| 13 and 15 | | 364 | 85 | 4 | 0.191 | 0.2804 | |
| 13 and 15 | [500,1000) | 453 | 80 | 1 | 0.1426 | 0.2149 | ✓ |
| 13 and 15 | | 453 | 103 | 2 | 0.1896 | 0.2688 | |
| 13 and 15 | | 453 | 142 | 3 | 0.271 | 0.3584 | ✓ |
| 13 and 15 | | 453 | 128 | 4 | 0.2415 | 0.3265 | |
| 13 and 15 | [1000,2000) | 785 | 146 | 1 | 0.1594 | 0.215 | ✓ |
| 13 and 15 | | 785 | 198 | 2 | 0.2222 | 0.2841 | |
| 13 and 15 | | 785 | 234 | 3 | 0.2663 | 0.3314 | ✓ |
| 13 and 15 | | 785 | 207 | 4 | 0.2332 | 0.296 | |
| 13 and 15 | [2000,3000) | 746 | 182 | 1 | 0.2135 | 0.2764 | |
| 13 and 15 | | 746 | 188 | 2 | 0.2212 | 0.2848 | |
| 13 and 15 | | 746 | 207 | 3 | 0.2456 | 0.3111 | |
| 13 and 15 | | 746 | 169 | 4 | 0.197 | 0.2583 | |
| 13 and 15 | [3000,5000) | 1469 | 381 | 1 | 0.2371 | 0.2826 | |
| 13 and 15 | | 1469 | 392 | 2 | 0.2444 | 0.2903 | |
| 13 and 15 | | 1469 | 392 | 3 | 0.2444 | 0.2903 | |
| 13 and 15 | | 1469 | 304 | 4 | 0.1865 | 0.2286 | ✓ |
| 13 and 15 | [5000,10000) | 2843 | 792 | 1 | 0.2622 | 0.2955 | ✓ |
| 13 and 15 | | 2843 | 711 | 2 | 0.2343 | 0.2664 | |
| 13 and 15 | | 2843 | 832 | 3 | 0.276 | 0.3098 | ✓ |
| 13 and 15 | | 2843 | 508 | 4 | 0.1648 | 0.1933 | ✓ |

| Pair | Distance interval (m) | Total | Count | Type | Lower CI | Upper CI | Sign. diff. |
|---|---|---|---|---|---|---|---|
| 14 and 15 | [0,50) | 92 | 17 | 1 | 0.1115 | 0.2793 | |
| 14 and 15 | | 92 | 13 | 2 | 0.0774 | 0.2295 | ✓ |
| 14 and 15 | | 92 | 26 | 3 | 0.1936 | 0.3861 | |
| 14 and 15 | | 92 | 36 | 4 | 0.2912 | 0.4986 | ✓ |
| 14 and 15 | [50,100) | 51 | 8 | 1 | 0.0702 | 0.2859 | |
| 14 and 15 | | 51 | 3 | 2 | 0.0123 | 0.1624 | ✓ |
| 14 and 15 | | 51 | 17 | 3 | 0.2076 | 0.4792 | |
| 14 and 15 | | 51 | 23 | 4 | 0.3113 | 0.5966 | ✓ |
| 14 and 15 | [100,200) | 55 | 4 | 1 | 0.0202 | 0.1759 | ✓ |
| 14 and 15 | | 55 | 6 | 2 | 0.0411 | 0.2225 | ✓ |
| 14 and 15 | | 55 | 22 | 3 | 0.2702 | 0.5409 | ✓ |
| 14 and 15 | | 55 | 23 | 4 | 0.2865 | 0.5589 | ✓ |
| 14 and 15 | [200,500) | 106 | 12 | 1 | 0.0599 | 0.1894 | ✓ |
| 14 and 15 | | 106 | 12 | 2 | 0.0599 | 0.1894 | ✓ |
| 14 and 15 | | 106 | 50 | 3 | 0.374 | 0.5711 | ✓ |
| 14 and 15 | | 106 | 32 | 4 | 0.2165 | 0.3987 | |
| 14 and 15 | [500,1000) | 109 | 14 | 1 | 0.072 | 0.2061 | ✓ |
| 14 and 15 | | 109 | 27 | 2 | 0.17 | 0.3396 | |
| 14 and 15 | | 109 | 36 | 3 | 0.2432 | 0.4269 | |
| 14 and 15 | | 109 | 32 | 4 | 0.2102 | 0.3885 | |
| 14 and 15 | [1000,2000) | 185 | 28 | 1 | 0.103 | 0.2113 | ✓ |
| 14 and 15 | | 185 | 49 | 2 | 0.2028 | 0.3346 | |
| 14 and 15 | | 185 | 60 | 3 | 0.2575 | 0.3969 | ✓ |
| 14 and 15 | | 185 | 48 | 4 | 0.1979 | 0.3289 | |
| 14 and 15 | [2000,3000) | 215 | 43 | 1 | 0.1487 | 0.2598 | |
| 14 and 15 | | 215 | 53 | 2 | 0.1904 | 0.3097 | |
| 14 and 15 | | 215 | 78 | 3 | 0.2985 | 0.4309 | ✓ |
| 14 and 15 | | 215 | 41 | 4 | 0.1405 | 0.2497 | ✓ |
| 14 and 15 | [3000,5000) | 269 | 52 | 1 | 0.1479 | 0.2456 | ✓ |
| 14 and 15 | | 269 | 70 | 2 | 0.2088 | 0.317 | |
| 14 and 15 | | 269 | 52 | 3 | 0.1479 | 0.2456 | ✓ |
| 14 and 15 | | 269 | 95 | 4 | 0.2961 | 0.4135 | ✓ |
| 14 and 15 | [5000,10000) | 1333 | 310 | 1 | 0.2101 | 0.2562 | |
| 14 and 15 | | 1333 | 299 | 2 | 0.2022 | 0.2477 | ✓ |
| 14 and 15 | | 1333 | 365 | 3 | 0.25 | 0.2986 | |
| 14 and 15 | | 1333 | 359 | 4 | 0.2457 | 0.294 | |

### *6.3. 30-min pairs*

| Pair | Distance interval (m) | Total | Count | Type | Lower CI | Upper CI | Sign. diff. |
|---|---|---|---|---|---|---|---|
| 1 and 4 | [0,50) | 3 | 0 | 1 | 0 | 0.7076 | |
| 1 and 4 | | 3 | 1 | 2 | 0.0084 | 0.9057 | |
| 1 and 4 | | 3 | 1 | 3 | 0.0084 | 0.9057 | |
| 1 and 4 | | 3 | 1 | 4 | 0.0084 | 0.9057 | |
| 1 and 4 | [50,100) | 10 | 2 | 1 | 0.0252 | 0.5561 | |
| 1 and 4 | | 10 | 2 | 2 | 0.0252 | 0.5561 | |
| 1 and 4 | | 10 | 3 | 3 | 0.0667 | 0.6525 | |
| 1 and 4 | | 10 | 3 | 4 | 0.0667 | 0.6525 | |
| 1 and 4 | [100,200) | 22 | 6 | 1 | 0.1073 | 0.5022 | |
| 1 and 4 | | 22 | 2 | 2 | 0.0112 | 0.2916 | |
| 1 and 4 | | 22 | 9 | 3 | 0.2071 | 0.6365 | |
| 1 and 4 | | 22 | 5 | 4 | 0.0782 | 0.4537 | |
| 1 and 4 | [200,500) | 132 | 16 | 1 | 0.0709 | 0.1894 | ✓ |
| 1 and 4 | | 132 | 23 | 2 | 0.1138 | 0.2499 | ✓ |
| 1 and 4 | | 132 | 51 | 3 | 0.3029 | 0.475 | ✓ |
| 1 and 4 | | 132 | 42 | 4 | 0.2399 | 0.4049 | |
| 1 and 4 | [500,1000) | 166 | 22 | 1 | 0.085 | 0.1938 | ✓ |
| 1 and 4 | | 166 | 42 | 2 | 0.1888 | 0.3262 | |
| 1 and 4 | | 166 | 53 | 3 | 0.2492 | 0.396 | |
| 1 and 4 | | 166 | 49 | 4 | 0.227 | 0.3708 | |
| 1 and 4 | [1000,2000) | 189 | 47 | 1 | 0.1888 | 0.3166 | |
| 1 and 4 | | 189 | 49 | 2 | 0.1984 | 0.3279 | |
| 1 and 4 | | 189 | 54 | 3 | 0.2225 | 0.3558 | |
| 1 and 4 | | 189 | 39 | 4 | 0.151 | 0.2711 | |
| 1 and 4 | [2000,3000) | 169 | 48 | 1 | 0.2174 | 0.3584 | |
| 1 and 4 | | 169 | 42 | 2 | 0.1854 | 0.3207 | |
| 1 and 4 | | 169 | 52 | 3 | 0.2391 | 0.3832 | |
| 1 and 4 | | 169 | 27 | 4 | 0.108 | 0.2239 | ✓ |
| 1 and 4 | [3000,5000) | 377 | 106 | 1 | 0.2363 | 0.3295 | |
| 1 and 4 | | 377 | 91 | 2 | 0.199 | 0.2878 | |
| 1 and 4 | | 377 | 88 | 3 | 0.1916 | 0.2795 | |
| 1 and 4 | | 377 | 92 | 4 | 0.2015 | 0.2906 | |
| 1 and 4 | [5000,10000) | 933 | 227 | 1 | 0.2161 | 0.2722 | |
| 1 and 4 | | 933 | 198 | 2 | 0.1864 | 0.2399 | ✓ |
| 1 and 4 | | 933 | 263 | 3 | 0.2532 | 0.3119 | ✓ |
| 1 and 4 | | 933 | 245 | 4 | 0.2346 | 0.2921 | |

| Pair | Distance interval (m) | Total | Count | Type | Lower CI | Upper CI | Sign. diff. |
|---|---|---|---|---|---|---|---|
| 1 and 6 | [0,50) | 176 | 17 | 1 | 0.0573 | 0.1501 | ✓ |
| 1 and 6 | | 176 | 10 | 2 | 0.0276 | 0.102 | ✓ |
| 1 and 6 | | 176 | 78 | 3 | 0.3685 | 0.5198 | ✓ |
| 1 and 6 | | 176 | 71 | 4 | 0.3303 | 0.4799 | ✓ |
| 1 and 6 | [50,100) | 357 | 33 | 1 | 0.0645 | 0.1274 | ✓ |
| 1 and 6 | | 357 | 20 | 2 | 0.0346 | 0.0852 | ✓ |
| 1 and 6 | | 357 | 181 | 3 | 0.4539 | 0.56 | ✓ |
| 1 and 6 | | 357 | 123 | 4 | 0.2953 | 0.3964 | ✓ |
| 1 and 6 | [100,200) | 873 | 70 | 1 | 0.063 | 0.1002 | ✓ |
| 1 and 6 | | 873 | 42 | 2 | 0.0349 | 0.0645 | ✓ |
| 1 and 6 | | 873 | 451 | 3 | 0.4829 | 0.5502 | ✓ |
| 1 and 6 | | 873 | 310 | 4 | 0.3233 | 0.3879 | ✓ |
| 1 and 6 | [200,500) | 1066 | 112 | 1 | 0.0873 | 0.125 | ✓ |
| 1 and 6 | | 1066 | 68 | 2 | 0.0499 | 0.0802 | ✓ |
| 1 and 6 | | 1066 | 517 | 3 | 0.4546 | 0.5155 | ✓ |
| 1 and 6 | | 1066 | 369 | 4 | 0.3176 | 0.3756 | ✓ |
| 1 and 6 | [500,1000) | 252 | 36 | 1 | 0.1021 | 0.1922 | ✓ |
| 1 and 6 | | 252 | 21 | 2 | 0.0523 | 0.1246 | ✓ |
| 1 and 6 | | 252 | 113 | 3 | 0.386 | 0.5121 | ✓ |
| 1 and 6 | | 252 | 82 | 4 | 0.2679 | 0.387 | ✓ |
| 1 and 6 | [1000,2000) | 86 | 22 | 1 | 0.1678 | 0.3613 | |
| 1 and 6 | | 86 | 10 | 2 | 0.0572 | 0.2035 | ✓ |
| 1 and 6 | | 86 | 29 | 3 | 0.2388 | 0.4472 | |
| 1 and 6 | | 86 | 25 | 4 | 0.1978 | 0.3986 | |
| 1 and 6 | [2000,3000) | 20 | 7 | 1 | 0.1539 | 0.5922 | |
| 1 and 6 | | 20 | 2 | 2 | 0.0123 | 0.317 | |
| 1 and 6 | | 20 | 5 | 3 | 0.0866 | 0.491 | |
| 1 and 6 | | 20 | 6 | 4 | 0.1189 | 0.5428 | |
| 1 and 6 | [3000,5000) | 39 | 6 | 1 | 0.0586 | 0.3053 | |
| 1 and 6 | | 39 | 6 | 2 | 0.0586 | 0.3053 | |
| 1 and 6 | | 39 | 7 | 3 | 0.0754 | 0.3353 | |
| 1 and 6 | | 39 | 20 | 4 | 0.3478 | 0.6758 | ✓ |
| 1 and 6 | [5000,10000) | 39 | 13 | 1 | 0.1909 | 0.5022 | |
| 1 and 6 | | 39 | 12 | 2 | 0.1702 | 0.4757 | |
| 1 and 6 | | 39 | 12 | 3 | 0.1702 | 0.4757 | |
| 1 and 6 | | 39 | 2 | 4 | 0.0063 | 0.1732 | ✓ |

| Pair | Distance interval (m) | Total | Count | Type | Lower CI | Upper CI | Sign. diff. |
|---|---|---|---|---|---|---|---|
| 2 and 3 | [0,50) | 12 | 1 | 1 | 0.0021 | 0.3848 | |
| 2 and 3 | | 12 | 2 | 2 | 0.0209 | 0.4841 | |
| 2 and 3 | | 12 | 5 | 3 | 0.1517 | 0.7233 | |
| 2 and 3 | | 12 | 4 | 4 | 0.0992 | 0.6511 | |
| 2 and 3 | [50,100) | 14 | 2 | 1 | 0.0178 | 0.4281 | |
| 2 and 3 | | 14 | 1 | 2 | 0.0018 | 0.3387 | |
| 2 and 3 | | 14 | 8 | 3 | 0.2886 | 0.8234 | ✓ |
| 2 and 3 | | 14 | 3 | 4 | 0.0466 | 0.508 | |
| 2 and 3 | [100,200) | 42 | 5 | 1 | 0.0398 | 0.2563 | |
| 2 and 3 | | 42 | 6 | 2 | 0.0543 | 0.2854 | |
| 2 and 3 | | 42 | 17 | 3 | 0.2563 | 0.5672 | ✓ |
| 2 and 3 | | 42 | 14 | 4 | 0.1957 | 0.4955 | |
| 2 and 3 | [200,500) | 135 | 23 | 1 | 0.1112 | 0.2446 | ✓ |
| 2 and 3 | | 135 | 22 | 2 | 0.105 | 0.2363 | ✓ |
| 2 and 3 | | 135 | 42 | 3 | 0.2343 | 0.3964 | |
| 2 and 3 | | 135 | 48 | 4 | 0.2751 | 0.4425 | ✓ |
| 2 and 3 | [500,1000) | 266 | 53 | 1 | 0.153 | 0.2524 | |
| 2 and 3 | | 266 | 34 | 2 | 0.0902 | 0.174 | ✓ |
| 2 and 3 | | 266 | 57 | 3 | 0.1665 | 0.2685 | |
| 2 and 3 | | 266 | 122 | 4 | 0.3976 | 0.5206 | ✓ |
| 2 and 3 | [1000,2000) | 498 | 114 | 1 | 0.1927 | 0.2684 | |
| 2 and 3 | | 498 | 85 | 2 | 0.1387 | 0.2067 | ✓ |
| 2 and 3 | | 498 | 94 | 3 | 0.1553 | 0.2259 | ✓ |
| 2 and 3 | | 498 | 205 | 4 | 0.3681 | 0.4563 | ✓ |
| 2 and 3 | [2000,3000) | 455 | 121 | 1 | 0.2259 | 0.3091 | |
| 2 and 3 | | 455 | 109 | 2 | 0.201 | 0.2815 | |
| 2 and 3 | | 455 | 101 | 3 | 0.1846 | 0.263 | |
| 2 and 3 | | 455 | 124 | 4 | 0.2321 | 0.3159 | |
| 2 and 3 | [3000,5000) | 1055 | 302 | 1 | 0.2591 | 0.3146 | ✓ |
| 2 and 3 | | 1055 | 250 | 2 | 0.2116 | 0.2638 | |
| 2 and 3 | | 1055 | 244 | 3 | 0.2061 | 0.2579 | |
| 2 and 3 | | 1055 | 259 | 4 | 0.2198 | 0.2726 | |
| 2 and 3 | [5000,10000) | 2202 | 567 | 1 | 0.2393 | 0.2763 | |
| 2 and 3 | | 2202 | 470 | 2 | 0.1965 | 0.2312 | ✓ |
| 2 and 3 | | 2202 | 522 | 3 | 0.2194 | 0.2554 | |
| 2 and 3 | | 2202 | 643 | 4 | 0.2731 | 0.3115 | ✓ |

| Pair | Distance interval (m) | Total | Count | Type | Lower CI | Upper CI | Sign. diff. |
|---|---|---|---|---|---|---|---|
| 2 and 6 | [0,50) | 6 | 0 | 1 | 0 | 0.4593 | |
| 2 and 6 | | 6 | 2 | 2 | 0.0433 | 0.7772 | |
| 2 and 6 | | 6 | 1 | 3 | 0.0042 | 0.6412 | |
| 2 and 6 | | 6 | 3 | 4 | 0.1181 | 0.8819 | |
| 2 and 6 | [50,100) | 20 | 3 | 1 | 0.0321 | 0.3789 | |
| 2 and 6 | | 20 | 2 | 2 | 0.0123 | 0.317 | |
| 2 and 6 | | 20 | 6 | 3 | 0.1189 | 0.5428 | |
| 2 and 6 | | 20 | 9 | 4 | 0.2306 | 0.6847 | |
| 2 and 6 | [100,200) | 40 | 5 | 1 | 0.0419 | 0.268 | |
| 2 and 6 | | 40 | 4 | 2 | 0.0279 | 0.2366 | ✓ |
| 2 and 6 | | 40 | 12 | 3 | 0.1656 | 0.4653 | |
| 2 and 6 | | 40 | 19 | 4 | 0.3151 | 0.6387 | ✓ |
| 2 and 6 | [200,500) | 98 | 19 | 1 | 0.121 | 0.2861 | |
| 2 and 6 | | 98 | 21 | 2 | 0.1378 | 0.3087 | |
| 2 and 6 | | 98 | 20 | 3 | 0.1293 | 0.2974 | |
| 2 and 6 | | 98 | 38 | 4 | 0.291 | 0.4915 | ✓ |
| 2 and 6 | [500,1000) | 149 | 32 | 1 | 0.1518 | 0.2894 | |
| 2 and 6 | | 149 | 29 | 2 | 0.1344 | 0.2674 | |
| 2 and 6 | | 149 | 33 | 3 | 0.1576 | 0.2967 | |
| 2 and 6 | | 149 | 55 | 4 | 0.2916 | 0.452 | ✓ |
| 2 and 6 | [1000,2000) | 324 | 71 | 1 | 0.1753 | 0.2682 | |
| 2 and 6 | | 324 | 86 | 2 | 0.2181 | 0.3171 | |
| 2 and 6 | | 324 | 65 | 3 | 0.1584 | 0.2484 | ✓ |
| 2 and 6 | | 324 | 102 | 4 | 0.2646 | 0.3685 | ✓ |
| 2 and 6 | [2000,3000) | 410 | 103 | 1 | 0.2099 | 0.2961 | |
| 2 and 6 | | 410 | 100 | 2 | 0.2031 | 0.2885 | |
| 2 and 6 | | 410 | 99 | 3 | 0.2008 | 0.2859 | |
| 2 and 6 | | 410 | 108 | 4 | 0.2214 | 0.3089 | |
| 2 and 6 | [3000,5000) | 1003 | 273 | 1 | 0.2448 | 0.3009 | |
| 2 and 6 | | 1003 | 234 | 2 | 0.2074 | 0.2607 | |
| 2 and 6 | | 1003 | 210 | 3 | 0.1846 | 0.2359 | ✓ |
| 2 and 6 | | 1003 | 286 | 4 | 0.2574 | 0.3142 | ✓ |
| 2 and 6 | [5000,10000) | 2266 | 527 | 1 | 0.2153 | 0.2505 | |
| 2 and 6 | | 2266 | 540 | 2 | 0.2209 | 0.2564 | |
| 2 and 6 | | 2266 | 580 | 3 | 0.2381 | 0.2745 | |
| 2 and 6 | | 2266 | 619 | 4 | 0.2549 | 0.292 | ✓ |

| Pair | Distance interval (m) | Total | Count | Type | Lower CI | Upper CI | Sign. diff. |
|---|---|---|---|---|---|---|---|
| 2 and 7 | [0,50) | 427 | 62 | 1 | 0.1132 | 0.1822 | ✓ |
| 2 and 7 | | 427 | 33 | 2 | 0.0538 | 0.1068 | ✓ |
| 2 and 7 | | 427 | 131 | 3 | 0.2634 | 0.3529 | ✓ |
| 2 and 7 | | 427 | 201 | 4 | 0.4226 | 0.5193 | ✓ |
| 2 and 7 | [50,100) | 533 | 41 | 1 | 0.0558 | 0.1029 | ✓ |
| 2 and 7 | | 533 | 34 | 2 | 0.0446 | 0.088 | ✓ |
| 2 and 7 | | 533 | 203 | 3 | 0.3395 | 0.4236 | ✓ |
| 2 and 7 | | 533 | 255 | 4 | 0.4353 | 0.5218 | ✓ |
| 2 and 7 | [100,200) | 1032 | 68 | 1 | 0.0515 | 0.0828 | ✓ |
| 2 and 7 | | 1032 | 66 | 2 | 0.0498 | 0.0806 | ✓ |
| 2 and 7 | | 1032 | 401 | 3 | 0.3587 | 0.4191 | ✓ |
| 2 and 7 | | 1032 | 497 | 4 | 0.4507 | 0.5126 | ✓ |
| 2 and 7 | [200,500) | 2200 | 222 | 1 | 0.0886 | 0.1143 | ✓ |
| 2 and 7 | | 2200 | 219 | 2 | 0.0873 | 0.1128 | ✓ |
| 2 and 7 | | 2200 | 847 | 3 | 0.3646 | 0.4057 | ✓ |
| 2 and 7 | | 2200 | 912 | 4 | 0.3939 | 0.4355 | ✓ |
| 2 and 7 | [500,1000) | 1615 | 259 | 1 | 0.1428 | 0.1792 | ✓ |
| 2 and 7 | | 1615 | 247 | 2 | 0.1357 | 0.1714 | ✓ |
| 2 and 7 | | 1615 | 533 | 3 | 0.3071 | 0.3536 | ✓ |
| 2 and 7 | | 1615 | 576 | 4 | 0.3333 | 0.3806 | ✓ |
| 2 and 7 | [1000,2000) | 1377 | 291 | 1 | 0.19 | 0.2339 | ✓ |
| 2 and 7 | | 1377 | 258 | 2 | 0.1671 | 0.209 | ✓ |
| 2 and 7 | | 1377 | 435 | 3 | 0.2914 | 0.3412 | ✓ |
| 2 and 7 | | 1377 | 393 | 4 | 0.2617 | 0.3101 | ✓ |
| 2 and 7 | [2000,3000) | 823 | 220 | 1 | 0.2373 | 0.299 | |
| 2 and 7 | | 823 | 195 | 2 | 0.2083 | 0.2675 | |
| 2 and 7 | | 823 | 212 | 3 | 0.228 | 0.2889 | |
| 2 and 7 | | 823 | 196 | 4 | 0.2094 | 0.2688 | |
| 2 and 7 | [3000,5000) | 1462 | 407 | 1 | 0.2555 | 0.3021 | ✓ |
| 2 and 7 | | 1462 | 330 | 2 | 0.2045 | 0.248 | ✓ |
| 2 and 7 | | 1462 | 378 | 3 | 0.2363 | 0.2818 | |
| 2 and 7 | | 1462 | 347 | 4 | 0.2157 | 0.26 | |
| 2 and 7 | [5000,10000) | 2279 | 626 | 1 | 0.2564 | 0.2935 | ✓ |
| 2 and 7 | | 2279 | 459 | 2 | 0.1851 | 0.2185 | ✓ |
| 2 and 7 | | 2279 | 613 | 3 | 0.2509 | 0.2877 | ✓ |
| 2 and 7 | | 2279 | 581 | 4 | 0.2371 | 0.2734 | |

| Pair | Distance interval (m) | Total | Count | Type | Lower CI | Upper CI | Sign. diff. |
|---|---|---|---|---|---|---|---|
| 3 and 7 | [0,50) | 6 | 0 | 1 | 0 | 0.4593 | |
| 3 and 7 | | 6 | 1 | 2 | 0.0042 | 0.6412 | |
| 3 and 7 | | 6 | 2 | 3 | 0.0433 | 0.7772 | |
| 3 and 7 | | 6 | 3 | 4 | 0.1181 | 0.8819 | |
| 3 and 7 | [50,100) | 8 | 0 | 1 | 0 | 0.3694 | |
| 3 and 7 | | 8 | 1 | 2 | 0.0032 | 0.5265 | |
| 3 and 7 | | 8 | 2 | 3 | 0.0319 | 0.6509 | |
| 3 and 7 | | 8 | 5 | 4 | 0.2449 | 0.9148 | |
| 3 and 7 | [100,200) | 28 | 1 | 1 | 9e-04 | 0.1835 | ✓ |
| 3 and 7 | | 28 | 7 | 2 | 0.1069 | 0.4487 | |
| 3 and 7 | | 28 | 11 | 3 | 0.215 | 0.5942 | |
| 3 and 7 | | 28 | 9 | 4 | 0.1588 | 0.5235 | |
| 3 and 7 | [200,500) | 104 | 22 | 1 | 0.1376 | 0.3026 | |
| 3 and 7 | | 104 | 21 | 2 | 0.1296 | 0.2919 | |
| 3 and 7 | | 104 | 37 | 3 | 0.2643 | 0.4557 | ✓ |
| 3 and 7 | | 104 | 24 | 4 | 0.1538 | 0.3236 | |
| 3 and 7 | [500,1000) | 240 | 43 | 1 | 0.1328 | 0.2336 | ✓ |
| 3 and 7 | | 240 | 47 | 2 | 0.1476 | 0.2518 | |
| 3 and 7 | | 240 | 107 | 3 | 0.3819 | 0.5111 | ✓ |
| 3 and 7 | | 240 | 43 | 4 | 0.1328 | 0.2336 | ✓ |
| 3 and 7 | [1000,2000) | 535 | 116 | 1 | 0.1826 | 0.2542 | |
| 3 and 7 | | 535 | 107 | 2 | 0.1669 | 0.2364 | ✓ |
| 3 and 7 | | 535 | 183 | 3 | 0.3019 | 0.384 | ✓ |
| 3 and 7 | | 535 | 129 | 4 | 0.2055 | 0.2797 | |
| 3 and 7 | [2000,3000) | 541 | 121 | 1 | 0.1892 | 0.2612 | |
| 3 and 7 | | 541 | 137 | 2 | 0.2171 | 0.2921 | |
| 3 and 7 | | 541 | 176 | 3 | 0.286 | 0.3666 | ✓ |
| 3 and 7 | | 541 | 107 | 4 | 0.165 | 0.2339 | ✓ |
| 3 and 7 | [3000,5000) | 968 | 244 | 1 | 0.225 | 0.2807 | |
| 3 and 7 | | 968 | 222 | 2 | 0.2032 | 0.2571 | |
| 3 and 7 | | 968 | 236 | 3 | 0.217 | 0.2721 | |
| 3 and 7 | | 968 | 266 | 4 | 0.2469 | 0.3041 | |
| 3 and 7 | [5000,10000) | 1931 | 487 | 1 | 0.233 | 0.2722 | |
| 3 and 7 | | 1931 | 404 | 2 | 0.1913 | 0.2281 | ✓ |
| 3 and 7 | | 1931 | 560 | 3 | 0.2698 | 0.3108 | ✓ |
| 3 and 7 | | 1931 | 480 | 4 | 0.2294 | 0.2685 | |

| Pair | Distance interval (m) | Total | Count | Type | Lower CI | Upper CI | Sign. diff. |
|---|---|---|---|---|---|---|---|
| 4 and 6 | [0,50) | 27 | 3 | 1 | 0.0235 | 0.2916 | |
| 4 and 6 | | 27 | 5 | 2 | 0.063 | 0.3808 | |
| 4 and 6 | | 27 | 10 | 3 | 0.194 | 0.5763 | |
| 4 and 6 | | 27 | 9 | 4 | 0.1652 | 0.5396 | |
| 4 and 6 | [50,100) | 74 | 12 | 1 | 0.0867 | 0.2661 | |
| 4 and 6 | | 74 | 7 | 2 | 0.0389 | 0.1852 | ✓ |
| 4 and 6 | | 74 | 32 | 3 | 0.3177 | 0.5528 | ✓ |
| 4 and 6 | | 74 | 23 | 4 | 0.2083 | 0.429 | |
| 4 and 6 | [100,200) | 228 | 25 | 1 | 0.0722 | 0.1576 | ✓ |
| 4 and 6 | | 228 | 28 | 2 | 0.0832 | 0.1726 | ✓ |
| 4 and 6 | | 228 | 85 | 3 | 0.3099 | 0.4391 | ✓ |
| 4 and 6 | | 228 | 90 | 4 | 0.3308 | 0.4614 | ✓ |
| 4 and 6 | [200,500) | 762 | 94 | 1 | 0.1008 | 0.1488 | ✓ |
| 4 and 6 | | 762 | 101 | 2 | 0.1093 | 0.1587 | ✓ |
| 4 and 6 | | 762 | 266 | 3 | 0.3152 | 0.3841 | ✓ |
| 4 and 6 | | 762 | 301 | 4 | 0.3601 | 0.4307 | ✓ |
| 4 and 6 | [500,1000) | 933 | 166 | 1 | 0.1539 | 0.204 | ✓ |
| 4 and 6 | | 933 | 162 | 2 | 0.1499 | 0.1995 | ✓ |
| 4 and 6 | | 933 | 324 | 3 | 0.3167 | 0.3788 | ✓ |
| 4 and 6 | | 933 | 281 | 4 | 0.2719 | 0.3317 | ✓ |
| 4 and 6 | [1000,2000) | 1057 | 235 | 1 | 0.1976 | 0.2486 | ✓ |
| 4 and 6 | | 1057 | 236 | 2 | 0.1985 | 0.2496 | ✓ |
| 4 and 6 | | 1057 | 294 | 3 | 0.2513 | 0.3062 | ✓ |
| 4 and 6 | | 1057 | 292 | 4 | 0.2495 | 0.3043 | |
| 4 and 6 | [2000,3000) | 946 | 242 | 1 | 0.2283 | 0.2849 | |
| 4 and 6 | | 946 | 206 | 2 | 0.1918 | 0.2454 | ✓ |
| 4 and 6 | | 946 | 258 | 3 | 0.2446 | 0.3023 | |
| 4 and 6 | | 946 | 240 | 4 | 0.2262 | 0.2827 | |
| 4 and 6 | [3000,5000) | 1656 | 458 | 1 | 0.2551 | 0.2988 | ✓ |
| 4 and 6 | | 1656 | 378 | 2 | 0.2082 | 0.2492 | ✓ |
| 4 and 6 | | 1656 | 389 | 3 | 0.2147 | 0.2561 | |
| 4 and 6 | | 1656 | 431 | 4 | 0.2393 | 0.2821 | |
| 4 and 6 | [5000,10000) | 3478 | 943 | 1 | 0.2564 | 0.2862 | ✓ |
| 4 and 6 | | 3478 | 781 | 2 | 0.2108 | 0.2388 | ✓ |
| 4 and 6 | | 3478 | 925 | 3 | 0.2513 | 0.281 | ✓ |
| 4 and 6 | | 3478 | 829 | 4 | 0.2243 | 0.2529 | |

| Pair | Distance interval (m) | Total | Count | Type | Lower CI | Upper CI | Sign. diff. |
|---|---|---|---|---|---|---|---|
| 8 and 12 | [0,50) | 6 | 2 | 1 | 0.0433 | 0.7772 | |
| 8 and 12 | | 6 | 0 | 2 | 0 | 0.4593 | |
| 8 and 12 | | 6 | 1 | 3 | 0.0042 | 0.6412 | |
| 8 and 12 | | 6 | 3 | 4 | 0.1181 | 0.8819 | |
| 8 and 12 | [50,100) | 11 | 1 | 1 | 0.0023 | 0.4128 | |
| 8 and 12 | | 11 | 0 | 2 | 0 | 0.2849 | |
| 8 and 12 | | 11 | 5 | 3 | 0.1675 | 0.7662 | |
| 8 and 12 | | 11 | 5 | 4 | 0.1675 | 0.7662 | |
| 8 and 12 | [100,200) | 39 | 6 | 1 | 0.0586 | 0.3053 | |
| 8 and 12 | | 39 | 6 | 2 | 0.0586 | 0.3053 | |
| 8 and 12 | | 39 | 13 | 3 | 0.1909 | 0.5022 | |
| 8 and 12 | | 39 | 14 | 4 | 0.212 | 0.5282 | |
| 8 and 12 | [200,500) | 142 | 18 | 1 | 0.0769 | 0.1929 | ✓ |
| 8 and 12 | | 142 | 15 | 2 | 0.0603 | 0.1682 | ✓ |
| 8 and 12 | | 142 | 52 | 3 | 0.287 | 0.4511 | ✓ |
| 8 and 12 | | 142 | 57 | 4 | 0.3201 | 0.4869 | ✓ |
| 8 and 12 | [500,1000) | 254 | 35 | 1 | 0.0979 | 0.1864 | ✓ |
| 8 and 12 | | 254 | 53 | 2 | 0.1604 | 0.2639 | |
| 8 and 12 | | 254 | 100 | 3 | 0.3332 | 0.4567 | ✓ |
| 8 and 12 | | 254 | 66 | 4 | 0.207 | 0.3184 | |
| 8 and 12 | [1000,2000) | 455 | 100 | 1 | 0.1826 | 0.2607 | |
| 8 and 12 | | 455 | 95 | 2 | 0.1723 | 0.2491 | ✓ |
| 8 and 12 | | 455 | 148 | 3 | 0.2824 | 0.3705 | ✓ |
| 8 and 12 | | 455 | 112 | 4 | 0.2072 | 0.2884 | |
| 8 and 12 | [2000,3000) | 430 | 91 | 1 | 0.174 | 0.2533 | |
| 8 and 12 | | 430 | 103 | 2 | 0.1999 | 0.2828 | |
| 8 and 12 | | 430 | 100 | 3 | 0.1934 | 0.2754 | |
| 8 and 12 | | 430 | 136 | 4 | 0.2726 | 0.3625 | ✓ |
| 8 and 12 | [3000,5000) | 873 | 220 | 1 | 0.2235 | 0.2822 | |
| 8 and 12 | | 873 | 220 | 2 | 0.2235 | 0.2822 | |
| 8 and 12 | | 873 | 207 | 3 | 0.2093 | 0.2668 | |
| 8 and 12 | | 873 | 226 | 4 | 0.2301 | 0.2893 | |
| 8 and 12 | [5000,10000) | 2630 | 652 | 1 | 0.2315 | 0.2649 | |
| 8 and 12 | | 2630 | 701 | 2 | 0.2497 | 0.2839 | |
| 8 and 12 | | 2630 | 647 | 3 | 0.2296 | 0.2629 | |
| 8 and 12 | | 2630 | 630 | 4 | 0.2233 | 0.2563 | |

| Pair | Distance interval (m) | Total | Count | Type | Lower CI | Upper CI | Sign. diff. |
|---|---|---|---|---|---|---|---|
| 8 and 13 | [0,50) | 3 | 0 | 1 | 0 | 0.7076 | |
| 8 and 13 | | 3 | 1 | 2 | 0.0084 | 0.9057 | |
| 8 and 13 | | 3 | 1 | 3 | 0.0084 | 0.9057 | |
| 8 and 13 | | 3 | 1 | 4 | 0.0084 | 0.9057 | |
| 8 and 13 | [50,100) | 11 | 1 | 1 | 0.0023 | 0.4128 | |
| 8 and 13 | | 11 | 4 | 2 | 0.1093 | 0.6921 | |
| 8 and 13 | | 11 | 6 | 3 | 0.2338 | 0.8325 | |
| 8 and 13 | | 11 | 0 | 4 | 0 | 0.2849 | |
| 8 and 13 | [100,200) | 31 | 4 | 1 | 0.0363 | 0.2983 | |
| 8 and 13 | | 31 | 4 | 2 | 0.0363 | 0.2983 | |
| 8 and 13 | | 31 | 10 | 3 | 0.1668 | 0.5137 | |
| 8 and 13 | | 31 | 13 | 4 | 0.2455 | 0.6092 | |
| 8 and 13 | [200,500) | 111 | 16 | 1 | 0.0847 | 0.2235 | ✓ |
| 8 and 13 | | 111 | 22 | 2 | 0.1286 | 0.2846 | |
| 8 and 13 | | 111 | 36 | 3 | 0.2385 | 0.4197 | |
| 8 and 13 | | 111 | 37 | 4 | 0.2467 | 0.4291 | |
| 8 and 13 | [500,1000) | 169 | 23 | 1 | 0.0883 | 0.1972 | ✓ |
| 8 and 13 | | 169 | 37 | 2 | 0.1591 | 0.2889 | |
| 8 and 13 | | 169 | 65 | 3 | 0.3109 | 0.4624 | ✓ |
| 8 and 13 | | 169 | 44 | 4 | 0.196 | 0.3333 | |
| 8 and 13 | [1000,2000) | 428 | 85 | 1 | 0.1618 | 0.2396 | ✓ |
| 8 and 13 | | 428 | 85 | 2 | 0.1618 | 0.2396 | ✓ |
| 8 and 13 | | 428 | 138 | 3 | 0.2783 | 0.369 | ✓ |
| 8 and 13 | | 428 | 120 | 4 | 0.2383 | 0.3255 | |
| 8 and 13 | [2000,3000) | 369 | 76 | 1 | 0.1658 | 0.2509 | |
| 8 and 13 | | 369 | 105 | 2 | 0.2391 | 0.3335 | |
| 8 and 13 | | 369 | 95 | 3 | 0.2136 | 0.3053 | |
| 8 and 13 | | 369 | 93 | 4 | 0.2085 | 0.2996 | |
| 8 and 13 | [3000,5000) | 885 | 212 | 1 | 0.2118 | 0.2691 | |
| 8 and 13 | | 885 | 210 | 2 | 0.2096 | 0.2667 | |
| 8 and 13 | | 885 | 228 | 3 | 0.2291 | 0.2878 | |
| 8 and 13 | | 885 | 235 | 4 | 0.2367 | 0.2959 | |
| 8 and 13 | [5000,10000) | 2543 | 645 | 1 | 0.2368 | 0.271 | |
| 8 and 13 | | 2543 | 654 | 2 | 0.2403 | 0.2746 | |
| 8 and 13 | | 2543 | 587 | 3 | 0.2146 | 0.2477 | ✓ |
| 8 and 13 | | 2543 | 657 | 4 | 0.2414 | 0.2758 | |

| Pair | Distance interval (m) | Total | Count | Type | Lower CI | Upper CI | Sign. diff. |
|---|---|---|---|---|---|---|---|
| 8 and 14 | [0,50) | 50 | 9 | 1 | 0.0858 | 0.3144 | |
| 8 and 14 | | 50 | 1 | 2 | 5e-04 | 0.1065 | ✓ |
| 8 and 14 | | 50 | 22 | 3 | 0.2999 | 0.5875 | ✓ |
| 8 and 14 | | 50 | 18 | 4 | 0.2292 | 0.5081 | |
| 8 and 14 | [50,100) | 92 | 6 | 1 | 0.0243 | 0.1366 | ✓ |
| 8 and 14 | | 92 | 9 | 2 | 0.0457 | 0.1776 | ✓ |
| 8 and 14 | | 92 | 45 | 3 | 0.3834 | 0.5956 | ✓ |
| 8 and 14 | | 92 | 32 | 4 | 0.2515 | 0.4543 | ✓ |
| 8 and 14 | [100,200) | 208 | 18 | 1 | 0.0521 | 0.1333 | ✓ |
| 8 and 14 | | 208 | 20 | 2 | 0.0597 | 0.1446 | ✓ |
| 8 and 14 | | 208 | 100 | 3 | 0.4112 | 0.5509 | ✓ |
| 8 and 14 | | 208 | 70 | 4 | 0.2727 | 0.4051 | ✓ |
| 8 and 14 | [200,500) | 638 | 50 | 1 | 0.0587 | 0.102 | ✓ |
| 8 and 14 | | 638 | 68 | 2 | 0.0837 | 0.1332 | ✓ |
| 8 and 14 | | 638 | 310 | 3 | 0.4465 | 0.5254 | ✓ |
| 8 and 14 | | 638 | 210 | 4 | 0.2928 | 0.3671 | ✓ |
| 8 and 14 | [500,1000) | 762 | 108 | 1 | 0.1177 | 0.1685 | ✓ |
| 8 and 14 | | 762 | 118 | 2 | 0.1299 | 0.1825 | ✓ |
| 8 and 14 | | 762 | 297 | 3 | 0.355 | 0.4254 | ✓ |
| 8 and 14 | | 762 | 239 | 4 | 0.2808 | 0.3479 | ✓ |
| 8 and 14 | [1000,2000) | 981 | 191 | 1 | 0.1704 | 0.2209 | ✓ |
| 8 and 14 | | 981 | 189 | 2 | 0.1684 | 0.2188 | ✓ |
| 8 and 14 | | 981 | 307 | 3 | 0.284 | 0.343 | ✓ |
| 8 and 14 | | 981 | 294 | 4 | 0.2712 | 0.3294 | ✓ |
| 8 and 14 | [2000,3000) | 744 | 171 | 1 | 0.2001 | 0.2618 | |
| 8 and 14 | | 744 | 142 | 2 | 0.1632 | 0.221 | ✓ |
| 8 and 14 | | 744 | 199 | 3 | 0.236 | 0.3008 | |
| 8 and 14 | | 744 | 232 | 4 | 0.2787 | 0.3465 | ✓ |
| 8 and 14 | [3000,5000) | 1394 | 361 | 1 | 0.2361 | 0.2828 | |
| 8 and 14 | | 1394 | 326 | 2 | 0.2119 | 0.257 | |
| 8 and 14 | | 1394 | 364 | 3 | 0.2382 | 0.285 | |
| 8 and 14 | | 1394 | 343 | 4 | 0.2236 | 0.2695 | |
| 8 and 14 | [5000,10000) | 3395 | 908 | 1 | 0.2526 | 0.2827 | ✓ |
| 8 and 14 | | 3395 | 767 | 2 | 0.2119 | 0.2404 | ✓ |
| 8 and 14 | | 3395 | 798 | 3 | 0.2209 | 0.2497 | ✓ |
| 8 and 14 | | 3395 | 922 | 4 | 0.2567 | 0.2869 | ✓ |

| Pair | Distance interval (m) | Total | Count | Type | Lower CI | Upper CI | Sign. diff. |
|---|---|---|---|---|---|---|---|
| 11 and 14 | [0,50) | 9 | 1 | 1 | 0.0028 | 0.4825 | |
| 11 and 14 | | 9 | 2 | 2 | 0.0281 | 0.6001 | |
| 11 and 14 | | 9 | 4 | 3 | 0.137 | 0.788 | |
| 11 and 14 | | 9 | 2 | 4 | 0.0281 | 0.6001 | |
| 11 and 14 | [50,100) | 6 | 2 | 1 | 0.0433 | 0.7772 | |
| 11 and 14 | | 6 | 0 | 2 | 0 | 0.4593 | |
| 11 and 14 | | 6 | 2 | 3 | 0.0433 | 0.7772 | |
| 11 and 14 | | 6 | 2 | 4 | 0.0433 | 0.7772 | |
| 11 and 14 | [100,200) | 6 | 2 | 1 | 0.0433 | 0.7772 | |
| 11 and 14 | | 6 | 0 | 2 | 0 | 0.4593 | |
| 11 and 14 | | 6 | 2 | 3 | 0.0433 | 0.7772 | |
| 11 and 14 | | 6 | 2 | 4 | 0.0433 | 0.7772 | |
| 11 and 14 | [200,500) | 91 | 9 | 1 | 0.0462 | 0.1795 | ✓ |
| 11 and 14 | | 91 | 17 | 2 | 0.1128 | 0.2822 | |
| 11 and 14 | | 91 | 33 | 3 | 0.2644 | 0.4701 | ✓ |
| 11 and 14 | | 91 | 32 | 4 | 0.2544 | 0.4588 | ✓ |
| 11 and 14 | [500,1000) | 191 | 26 | 1 | 0.0909 | 0.1931 | ✓ |
| 11 and 14 | | 191 | 39 | 2 | 0.1494 | 0.2684 | |
| 11 and 14 | | 191 | 74 | 3 | 0.318 | 0.4605 | ✓ |
| 11 and 14 | | 191 | 52 | 4 | 0.2105 | 0.3412 | |
| 11 and 14 | [1000,2000) | 510 | 93 | 1 | 0.1498 | 0.2187 | ✓ |
| 11 and 14 | | 510 | 105 | 2 | 0.1716 | 0.2436 | ✓ |
| 11 and 14 | | 510 | 147 | 3 | 0.2493 | 0.3297 | |
| 11 and 14 | | 510 | 165 | 4 | 0.2831 | 0.366 | ✓ |
| 11 and 14 | [2000,3000) | 556 | 115 | 1 | 0.1739 | 0.2429 | ✓ |
| 11 and 14 | | 556 | 141 | 2 | 0.2179 | 0.2919 | |
| 11 and 14 | | 556 | 128 | 3 | 0.1958 | 0.2675 | |
| 11 and 14 | | 556 | 172 | 4 | 0.2711 | 0.3496 | ✓ |
| 11 and 14 | [3000,5000) | 1016 | 275 | 1 | 0.2436 | 0.2991 | |
| 11 and 14 | | 1016 | 279 | 2 | 0.2474 | 0.3032 | |
| 11 and 14 | | 1016 | 222 | 3 | 0.1934 | 0.2452 | ✓ |
| 11 and 14 | | 1016 | 240 | 4 | 0.2104 | 0.2636 | |
| 11 and 14 | [5000,10000) | 2285 | 693 | 1 | 0.2845 | 0.3226 | ✓ |
| 11 and 14 | | 2285 | 525 | 2 | 0.2126 | 0.2476 | ✓ |
| 11 and 14 | | 2285 | 540 | 3 | 0.219 | 0.2543 | |
| 11 and 14 | | 2285 | 527 | 4 | 0.2135 | 0.2485 | ✓ |

| Pair | Distance interval (m) | Total | Count | Type | Lower CI | Upper CI | Sign. diff. |
|---|---|---|---|---|---|---|---|
| 12 and 13 | [0,50) | 9 | 0 | 1 | 0 | 0.3363 | |
| 12 and 13 | | 9 | 4 | 2 | 0.137 | 0.788 | |
| 12 and 13 | | 9 | 1 | 3 | 0.0028 | 0.4825 | |
| 12 and 13 | | 9 | 4 | 4 | 0.137 | 0.788 | |
| 12 and 13 | [50,100) | 45 | 9 | 1 | 0.0958 | 0.346 | |
| 12 and 13 | | 45 | 2 | 2 | 0.0054 | 0.1515 | ✓ |
| 12 and 13 | | 45 | 17 | 3 | 0.2377 | 0.5346 | |
| 12 and 13 | | 45 | 17 | 4 | 0.2377 | 0.5346 | |
| 12 and 13 | [100,200) | 126 | 15 | 1 | 0.0682 | 0.1887 | ✓ |
| 12 and 13 | | 126 | 12 | 2 | 0.0502 | 0.1605 | ✓ |
| 12 and 13 | | 126 | 50 | 3 | 0.3108 | 0.4878 | ✓ |
| 12 and 13 | | 126 | 49 | 4 | 0.3034 | 0.4798 | ✓ |
| 12 and 13 | [200,500) | 437 | 52 | 1 | 0.0902 | 0.1531 | ✓ |
| 12 and 13 | | 437 | 46 | 2 | 0.0781 | 0.1379 | ✓ |
| 12 and 13 | | 437 | 138 | 3 | 0.2724 | 0.3617 | ✓ |
| 12 and 13 | | 437 | 201 | 4 | 0.4125 | 0.508 | ✓ |
| 12 and 13 | [500,1000) | 669 | 106 | 1 | 0.1316 | 0.1884 | ✓ |
| 12 and 13 | | 669 | 97 | 2 | 0.1192 | 0.174 | ✓ |
| 12 and 13 | | 669 | 239 | 3 | 0.3209 | 0.3949 | ✓ |
| 12 and 13 | | 669 | 227 | 4 | 0.3035 | 0.3766 | ✓ |
| 12 and 13 | [1000,2000) | 895 | 170 | 1 | 0.1647 | 0.2172 | ✓ |
| 12 and 13 | | 895 | 172 | 2 | 0.1669 | 0.2195 | ✓ |
| 12 and 13 | | 895 | 257 | 3 | 0.2577 | 0.318 | ✓ |
| 12 and 13 | | 895 | 296 | 4 | 0.2999 | 0.3626 | ✓ |
| 12 and 13 | [2000,3000) | 701 | 147 | 1 | 0.1801 | 0.2417 | ✓ |
| 12 and 13 | | 701 | 132 | 2 | 0.16 | 0.2192 | ✓ |
| 12 and 13 | | 701 | 225 | 3 | 0.2865 | 0.3569 | ✓ |
| 12 and 13 | | 701 | 197 | 4 | 0.248 | 0.3159 | |
| 12 and 13 | [3000,5000) | 1469 | 355 | 1 | 0.22 | 0.2644 | |
| 12 and 13 | | 1469 | 340 | 2 | 0.2101 | 0.2539 | |
| 12 and 13 | | 1469 | 377 | 3 | 0.2345 | 0.2798 | |
| 12 and 13 | | 1469 | 397 | 4 | 0.2477 | 0.2937 | |
| 12 and 13 | [5000,10000) | 3441 | 948 | 1 | 0.2606 | 0.2908 | ✓ |
| 12 and 13 | | 3441 | 836 | 2 | 0.2287 | 0.2576 | |
| 12 and 13 | | 3441 | 834 | 3 | 0.2281 | 0.257 | |
| 12 and 13 | | 3441 | 823 | 4 | 0.225 | 0.2538 | |

| Pair | Distance interval (m) | Total | Count | Type | Lower CI | Upper CI | Sign. diff. |
|---|---|---|---|---|---|---|---|
| 13 and 14 | [0,50) | 4 | 0 | 1 | 0 | 0.6024 | |
| 13 and 14 | | 4 | 1 | 2 | 0.0063 | 0.8059 | |
| 13 and 14 | | 4 | 2 | 3 | 0.0676 | 0.9324 | |
| 13 and 14 | | 4 | 1 | 4 | 0.0063 | 0.8059 | |
| 13 and 14 | [50,100) | 9 | 1 | 1 | 0.0028 | 0.4825 | |
| 13 and 14 | | 9 | 1 | 2 | 0.0028 | 0.4825 | |
| 13 and 14 | | 9 | 3 | 3 | 0.0749 | 0.7007 | |
| 13 and 14 | | 9 | 4 | 4 | 0.137 | 0.788 | |
| 13 and 14 | [100,200) | 28 | 3 | 1 | 0.0227 | 0.2823 | |
| 13 and 14 | | 28 | 3 | 2 | 0.0227 | 0.2823 | |
| 13 and 14 | | 28 | 11 | 3 | 0.215 | 0.5942 | |
| 13 and 14 | | 28 | 11 | 4 | 0.215 | 0.5942 | |
| 13 and 14 | [200,500) | 86 | 9 | 1 | 0.049 | 0.1894 | ✓ |
| 13 and 14 | | 86 | 12 | 2 | 0.0742 | 0.2311 | ✓ |
| 13 and 14 | | 86 | 37 | 3 | 0.3239 | 0.5415 | ✓ |
| 13 and 14 | | 86 | 28 | 4 | 0.2284 | 0.4352 | |
| 13 and 14 | [500,1000) | 151 | 28 | 1 | 0.1269 | 0.2567 | |
| 13 and 14 | | 151 | 21 | 2 | 0.0882 | 0.2047 | ✓ |
| 13 and 14 | | 151 | 54 | 3 | 0.2814 | 0.4396 | ✓ |
| 13 and 14 | | 151 | 48 | 4 | 0.2446 | 0.3985 | |
| 13 and 14 | [1000,2000) | 314 | 68 | 1 | 0.1723 | 0.2663 | |
| 13 and 14 | | 314 | 67 | 2 | 0.1694 | 0.2629 | |
| 13 and 14 | | 314 | 78 | 3 | 0.2016 | 0.3 | |
| 13 and 14 | | 314 | 101 | 4 | 0.2703 | 0.3764 | ✓ |
| 13 and 14 | [2000,3000) | 354 | 77 | 1 | 0.1756 | 0.2642 | |
| 13 and 14 | | 354 | 88 | 2 | 0.2044 | 0.297 | |
| 13 and 14 | | 354 | 77 | 3 | 0.1756 | 0.2642 | |
| 13 and 14 | | 354 | 112 | 4 | 0.2682 | 0.3676 | ✓ |
| 13 and 14 | [3000,5000) | 758 | 174 | 1 | 0.2001 | 0.2612 | |
| 13 and 14 | | 758 | 213 | 2 | 0.2492 | 0.3145 | |
| 13 and 14 | | 758 | 156 | 3 | 0.1776 | 0.2364 | ✓ |
| 13 and 14 | | 758 | 215 | 4 | 0.2518 | 0.3172 | ✓ |
| 13 and 14 | [5000,10000) | 2141 | 569 | 1 | 0.2471 | 0.285 | |
| 13 and 14 | | 2141 | 554 | 2 | 0.2403 | 0.2779 | |
| 13 and 14 | | 2141 | 503 | 3 | 0.2171 | 0.2535 | |
| 13 and 14 | | 2141 | 515 | 4 | 0.2226 | 0.2592 | |
